# Integrated design, execution, and analysis of arrayed and pooled CRISPR genome editing experiments

**DOI:** 10.1101/125245

**Authors:** Matthew C. Canver, Maximilian Haeussler, Daniel E. Bauer, Stuart H. Orkin, Neville E. Sanjana, Ophir Shalem, Guo-Cheng Yuan, Feng Zhang, Jean-Paul Concordet, Luca Pinello

**Affiliations:** Department of Molecular Pathology, Massachusetts General Hospital, Harvard Medical School, Boston, Massachusetts, USA; Santa Cruz Genomics Institute, MS CBSE, University of California, Santa Cruz, California, USA; Division of Hematology/Oncology, Boston Children’s Hospital, Department of Pediatric Oncology, Dana-Farber Cancer Institute, Harvard Stem Cell Institute, Department of Pediatrics, Harvard Medical School, Boston, Massachusetts, USA; Howard Hughes Medical Institute, Boston, Massachusetts, USA; New York Genome Center and Department of Biology, New York University, New York City, New York, USA; Raymond G. Perelman Center for Cellular and Molecular Therapeutics, Children’s Hospital of Philadelphia, Department of Genetics, University of Pennsylvania, Philadelphia, Pennsylvania, USA; Department of Biostatistics and Computational Biology, Dana-Farber Cancer Institute and Harvard T.H. Chan School of Public Health, Boston, Massachusetts, USA; The Broad Institute, Cambridge, Massachusetts, USA.; McGovern Institute for Brain Research, Department of Brain and Cognitive Sciences, Department of Biological Engineering, Massachusetts Institute of Technology, Cambridge, Massachusetts, USA.; INSERM U1154, CNRS UMR 7196, Muséum National d’Histoire Naturelle, Paris, France.

**Author notes:** Co-first authors (Matthew C. Canver, Maximilian Haeussler). Correspondence to Luca Pinello.

**Keywords:** CRISPR genome editing, Cas9/Cpf1, CRISPR pooled screening, Saturating mutagenesis, sgRNA design, Lentivirus, CRISPOR, CRISPResso, Docker, Locus-specific deep sequencing, Off-target effect, On-target effect, Amplicon sequencing, Targeted sequencing, Sequence analysis, Computational tools, Software pipeline

## Abstract

CRISPR genome editing experiments offer enormous potential for the evaluation of genomic loci using arrayed single guide RNAs (sgRNAs) or pooled sgRNA libraries. Numerous computational tools are available to help design sgRNAs with optimal on-target efficiency and minimal off-target potential. In addition, computational tools have been developed to analyze deep sequencing data resulting from genome editing experiments. However, these tools are typically developed in isolation and oftentimes not readily translatable into laboratory-based experiments. Here we present a protocol that describes in detail both the computational and benchtop implementation of an arrayed and/or pooled CRISPR genome editing experiment. This protocol provides instructions for sgRNA design with CRISPOR, experimental implementation, and analysis of the resulting high-throughput sequencing data with CRISPResso. This protocol allows for design and execution of arrayed and pooled CRISPR experiments in 4-5 weeks by non-experts as well as computational data analysis in 1-2 days that can be performed by both computational and non-computational biologists alike.

## INTRODUCTION

The clustered regularly interspaced short palindromic repeats (CRISPR) nuclease system is a facile and robust genome editing system^1,2^. The CRISPR nuclease system was identified as the driver of prokaryotic adaptive immunity to allow for resistance to bacteriophages^3^. This system has been subsequently repurposed for eukaryotic genome editing by heterologous expression of the CRISPR components in eukaryotic cells. Site-specific cleavage by Cas9 requires an RNA molecule to guide nucleases to specific genomic loci to initiate double-strand breaks (DSBs)^1,2,4^. Site-specific cleavage requires Watson-Crick base pairing of the RNA molecule to a corresponding genomic sequence upstream of a protospacer adjacent motif (PAM)^1,2^. The required RNA molecule for genome editing experiments consists of a synthetic fusion of the prokaryotic tracrRNA and crRNA to create a single chimeric guide RNA (sgRNA)^5^. In contrast, the Cpf1 nuclease does not require a tracrRNA and engenders DSBs downstream of its PAM sequence. Cpf1 requires a CRISPR RNA (crRNA) to create DSBs that include 5’ overhangs^4^.

CRISPR mutagenesis relies on engagement of endogenous DNA repair pathways after nuclease-mediated DSB induction has occurred. The principal repair pathways include non-homologous end joining (NHEJ) and homology-directed repair (HDR). NHEJ repair is an error-prone pathway, which results in a heterogeneous spectrum of insertions/deletions (indels) primarily in the range of 1-10 bp^1,2,6–8^. HDR relies on the co-delivery of an extrachromosomal template to be used as a template for DNA repair as opposed to an endogenous template such as a sister chromatid. This allows for the insertion of customized sequence into the genome^1,2^.

### Applications of the method and development of the protocol

A variety of computational tools have been developed for the design and analysis of CRISPR-based experiments. However, these tools are typically developed in isolation without features for facile integration with one another and/or without sufficient consideration to facilitate implementation in a laboratory setting. Here, we offer a protocol to integrate robust, publicly available tools for the design, execution, and analysis of CRISPR genome editing experiments. Specifically, we have adapted CRISPOR^9^ and CRISPResso^10^ to be integrated with one another as well as streamlined for experimental implementation (Figs. 1-3). This protocol has been used in previously published works^11,12^.

**Fig. 1:**
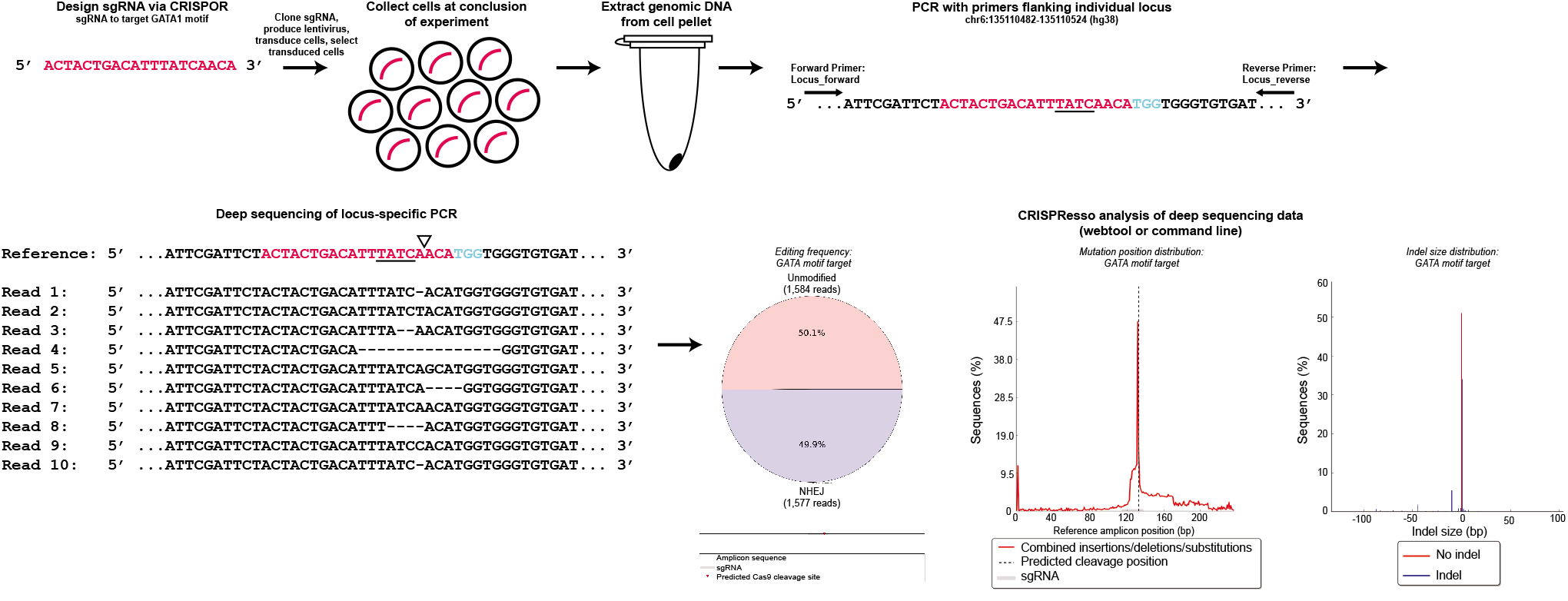
Schematic of an arrayed genome editing experiment. Arrayed genome editing experiments are performed by designing one sgRNA using CRISPOR. This schematic demonstrates design of an sgRNA to mutagenize a GATA motif. After designing the optimal sgRNA, it is cloned into pLentiGuide-puro, lentivirus produced, cells transduced, and successful transductants are selected (successful transduction is indicated by red curved lines). After conclusion of the experiment, cells are pelleted and genomic DNA is extracted. Locus-specific PCR primers are used to amplify regions flanking the double-strand break site. Deep sequencing of the amplicon generated by locus-specific PCR is subsequently performed. Quantification of editing frequency and indel distribution is determined by CRISPResso. The GATA motif is underlined, the triangle indicates the double-strand break position, and the PAM sequence is shown in blue.

CRISPR mutagenesis allows for the study of both coding and non-coding regions of the genome^13^. This involves usage of one sgRNA for arrayed experiments or multiple sgRNAs for pooled experiments. Arrayed experiments are useful when a target can be mutagenized by one sgRNA. Pooled screening allows for targeting of a handful of genes up to genome-scale gene targeting^14–17^. It also allows for saturating mutagenesis (tiling of sgRNA) experiments to identify functional sequence within non^11,12,18^ coding regions. This protocol can be used to design and execute arrayed or pooled genome editing experiments. Furthermore, it can also be used for the design, implementation, and analysis of pooled screens for gene-targeting, saturating mutagenesis of non-coding elements, or any other targeting^11,12,19,20^ strategies.

### Comparison with other methods

Numerous computational tools are freely available to aid sgRNA design for a wide spectrum of PAM sequences as well as on-target efficiency and off-target cleavage predictions: Broad GPP Portal^21^, Cas-database^22^, Cas-OFFinder^23^, CasOT^24^, CCTop^25^, COSMID^26^, CHOPCHOP^27,28^, CRISPRdirect^29^, CRISPR-DO^30^, CRISPR-ERA^31^, CRISPR-P^32^, CROP-IT^33^, DNA Striker^12^, E-CRISP^34^, flyCRISPR^35^, GuideScan^36^, GT-scan^37^, MIT CRISPR design tool^8^, WU-CRISPR^38^, CRISPRseek^39^, sgRNAcas9^40^, CRISPR multiTargeter^41^, as well as others offered by companies such as Deskgen^42^ and Benchling^42^. CRISPOR (http://crispor.org) is a computational tool that predicts off-target cleavage sites and offers a variety of on-target efficiency scoring systems to assist sgRNA selection for more than 120 genomes using many different CRISPR nucleases (Table 1)^9^. CRISPOR offers several unique advantages for designing sgRNA for genome editing experiments. First, CRISPOR integrates multiple published on-target sgRNA efficiency scores including from Fusi et al^43^, Chari et al^44^, Xu et al^45^, Doench et al (Doench 2014 and 2016)^21,46^, Wang et al^47^, Moreno-Mateos et al^48^, Housden et al^49^, Prox. GC^50^, -GG^51^, and Out-of-Frame^52^. It also offers previously published off-target prediction^8^. Second, CRISPOR has been optimized to facilitate experimental implementation by providing automated primer design for both on-target and off-target deep sequencing analysis. The primers and output files are further designed to be compatible for subsequent analysis by CRISPResso after the experiments have been completed. Added after the initial publication, CRISPOR now offers features to automate oligonucleotide design for cloning pooled sgRNA libraries targeting both genes and non-coding regions. Taken together, CRISPOR provides an sgRNA design platform to facilitate experimental execution and downstream data analysis.

**Table 1.**
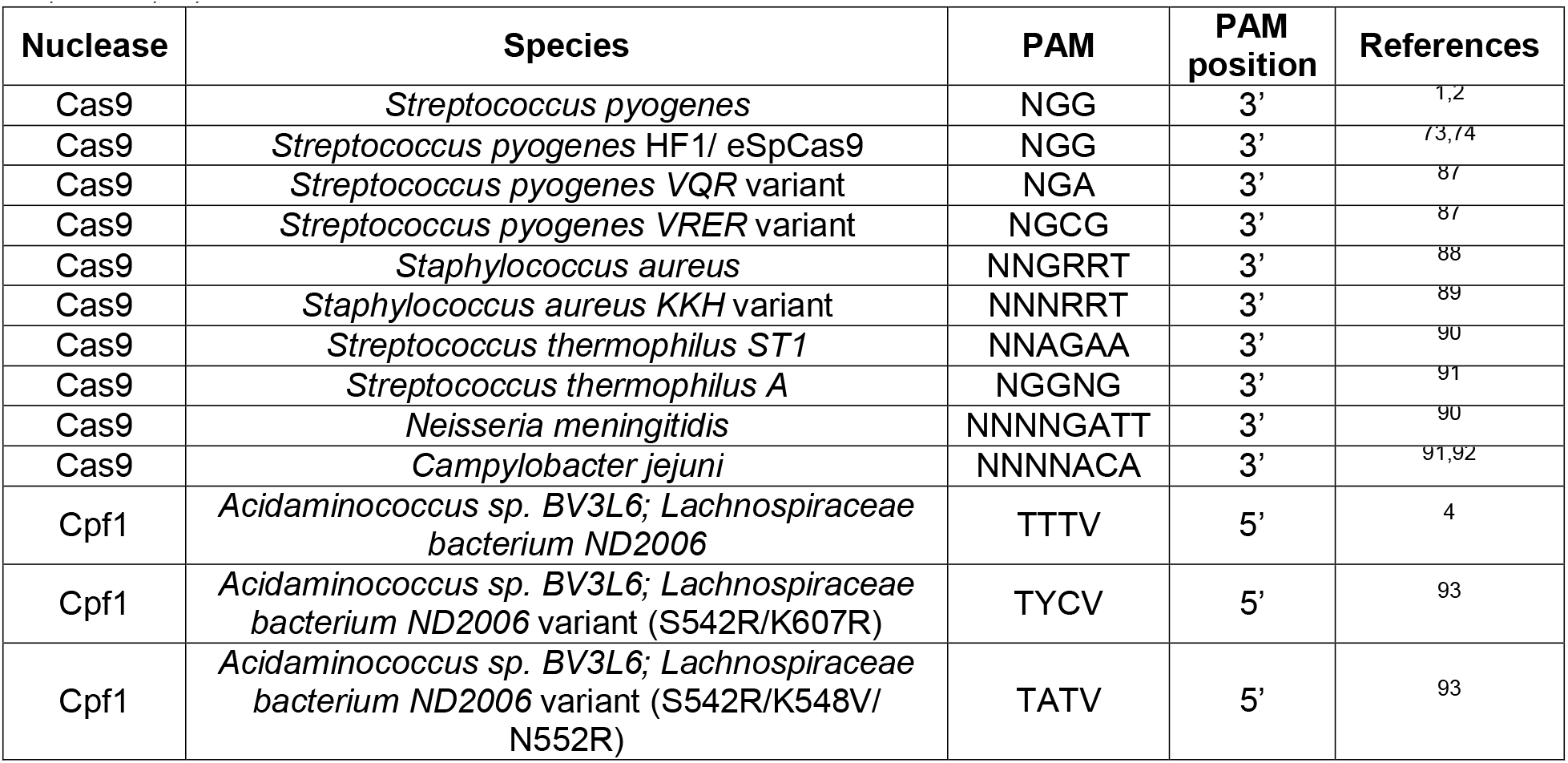
|List of available PAM sequences for CRISPOR analysis. R = A or G; Y = C or T; V = A, C, or G; N = A, C, G or T

CRISPResso is a computational pipeline that enables accurate quantification and visualization of CRISPR genome editing outcomes, as well as comprehensive evaluation of effects on coding sequences, non-coding elements, and off-target sites from individual loci, pooled amplicons, and whole genome deep-sequencing data^10^. The CRISPResso suite involves multiple tools for analysis, including the CRISPResso webtool and command line version of CRISPResso. There are also multiple additional command line tools: CRISPRessoPooled, CRISPRessoWGS, CRISPRessoCount, CRISPRessoCompare, and CRISPRessoPooledWGSCompare. The applications and features of these tools are summarized in Table 2. CRISPResso analysis offers many unique features such as splice-site analysis or frameshift analysis to quantify the proportion of engendered mutations that result in a frameshift when targeting coding sequence. In addition, features for indel visualization have been added to CRISPResso since its initial publication^10^. Alternative computational tools exist to evaluate genome editing outcomes from deep sequencing data^53–57^; however, these tools offer limited analysis functionality for pooled amplicon sequencing or WGS data as compared to the CRISPResso suite. CrispRVariants is another tool that offers functionality to analyze deep sequencing data by quantifying mosaicism and allele-specific gene editing as well as multi-sequence alignment views^56^.

**Table 2.**
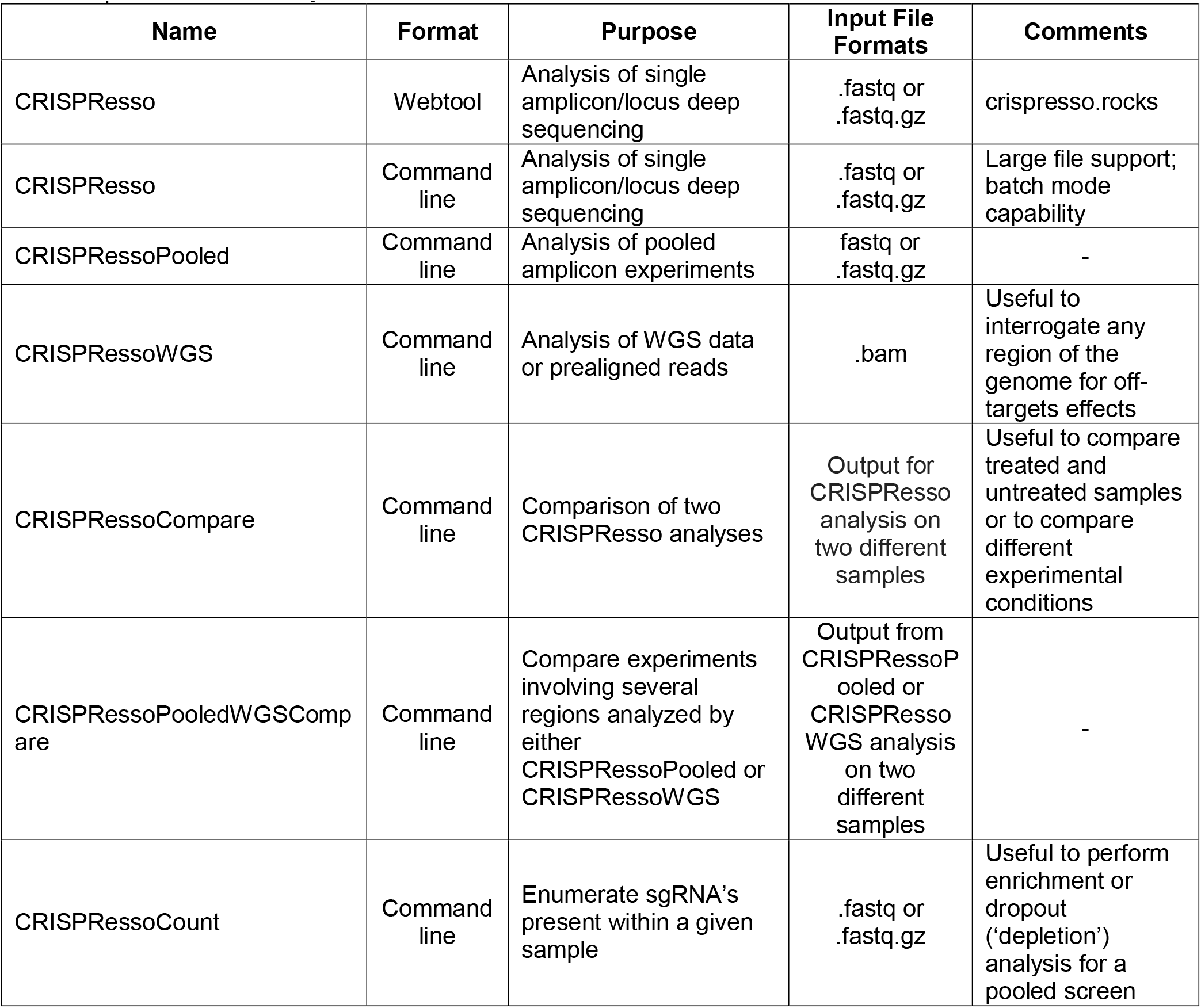
|CRISPResso analysis suite

CRISPR genome editing reagents have taken many forms, including DNA, RNA, protein, or various combinations of each^58,59^. Furthermore, delivery of these reagents has also been attempted using a variety of methods, including electroporation, lipid-based transfection, and viral-mediated delivery^58,59^. Pooled screening relies on the ability to deliver individual reagents to individual cells in batches. Electroporation and lipid-based transfection methods offer limited ability to control the number of reagents (i.e., sgRNA) delivered per cell; however, lentiviral transduction at low transduction rates (~30-50%) results in single lentiviral integrants per cell in the majority of cases (see Box 4 for further discussion)^11^. Furthermore, lentivirus offers stable integration of the CRISPR reagents into each cell’s genome. These features of lentivirus allow for pooled CRISPR experiments. Therefore, this protocol uses lentivirus for both arrayed and pooled CRISPR experiments.

### Limitations of on-target and off-target prediction

As described above, there are many tools available for both on-target sgRNA efficiency and off-target cleavage prediction. Progress has been made towards enhancing the predictive value of these scores; however, while these predictions are useful to focus sgRNA selection for experimental design, experimental validation provides the definitive analysis of on-target and off-target mutagenesis. Alternatively, experimental approaches have also been developed for unbiased genome-wide detection of off-target cleavages^60–66^. Continued investigation is necessary to more completely understand the rules governing sgRNA efficiency and off-target mutagenesis (see Box 1 for further discussion).

### Experimental design

#### Target identification and nuclease choice

CRISPR genome editing experiments require appropriate target identification to fit experimental objectives, which can include functional analysis of gene-or noncoding sequence (e.g., enhancers, CTCF or other transcription factor binding sites). Gene knockout is usually accomplished by targeting exonic sequence, but it can also be achieved through promoter disruption. Gene knockout is often complicated by alternative splicing and/or expression of multiple isoforms for a given gene of interest. It is difficult to predict the relevant isoform without prior knowledge; however, it is often possible to target exonic sequence common to all isoforms or to design sgRNA targeting unique isoforms to aid in the identification of a relevant/functional isoform. In the case of gene families, conserved domains can be identified for disruption^34^. While exon targeting toward the 5’ end of the gene has been shown to be more effective for functional disruption than the 3’ end^46^, it is also possible to target functional protein domains. In this case, even in-frame mutations frequently disrupt protein function^67^. Similar considerations can be applied to CRISPR interference (CRISPRi) or CRISPR activation (CRISPRa) approaches; however, CRISPRi/CRISPRa require sgRNA targeting in close proximity to the transcriptional start state for maximal repression/activation^17,68–72^.

Each CRISPR nuclease offers a unique PAM sequence, which has varying frequency within the genome^12^. PAM frequency can change depending on the location of targeting such as exons, introns, promoters, DNase hypersensitive sites, enhancers, or repressed regions^12^. The optimal nuclease can be chosen based on the density of available PAMs such as for saturating mutagenesis or proximity of PAMs to a particular genomic position such a transcription factor binding motif. It can be useful to use a nuclease with a GC-rich PAM for GC-rich sequences or a nuclease with an AT-rich PAM for AT-rich sequences (Table 1). Therefore, choosing the optimal nuclease depends on the region(s) of interest to be analyzed. Finally, high fidelity nucleases can be used to minimize the probability of off-target mutagenesis^73,74^.

#### Positive, negative, and editing controls for arrayed and pooled screen CRISPR experiments

Positive and negative controls are essential for both arrayed and pooled screen experiments. Positive controls are generally experiment-specific; however, some widely used positive controls include targeting known essential genes (e.g., ribosomal genes, housekeeping genes) when performing experiments to identify novel essential genes or when performing a dropout (‘depletion’) screen (see Box 4 for further discussion of dropout screens). Another example includes targeting exonic sequences as a positive control when targeting non-coding sequences potentially affects gene expression^11,12^. While a single positive control sgRNA is often used for an arrayed experiment, multiple positive control sgRNAs are typically used for pooled screen experiments, oftentimes comprising 1-5% of the number of sgRNA in the library (see Box 4 for further discussion)^11,12^.

Negative controls typically take the form of non-targeting controls, which are sgRNAs without any perfect matches in a given genome and with low potential for cleavage at genomic loci with imperfect matches (selected by minimization of sites with few mismatches, which is similar to minimizing off-target effects for targeted sgRNA). Non-targeting controls are genome build-, species-and PAM-specific (unless designed to be compatible with multiple genomes and/or PAMs). Other options for negative controls include targeting a safe harbor locus such as *AAVS1* or a gene/region known to have no effect on cellular function or the phenotype of interest. While a single negative control sgRNA is often used for an arrayed experiment, multiple negative control sgRNAs are typically used for pooled screen experiments, oftentimes comprising 1-5% of the number of sgRNA in the library (see Box 4 for further discussion)^11,12^. Non-targeting sgRNA can be generated “by-hand” via a guess and check approach to ensure no genomic matches (and minimal sites with few mismatches) or can be designed using previously published tools^12^.

An editing control can also be included to ensure proper functioning of CRISPR reagents, particularly nuclease function, in the cells under study (see Boxes 3-4 for further discussion). One possibility is to include a construct that expresses GFP together with an sgRNA targeting GFP to assess for functional Cas9 expression by flow cytometry^12,46^. This is particularly informative when using cell lines with stable Cas9 expression and can be helpful for troubleshooting low or absent editing rates. Typically, only one reporter system/sgRNA is needed for both arrayed and pooled experiments.

#### sgRNA and PCR primer design for arrayed and pooled screen experiments using CRISPOR

Arrayed experiments involve the use of an sgRNA for editing of a single target. CRISPOR offers a variety of on- and off-target prediction scores that can aid in the optimal sgRNA selection (Box 1)^9^. Analysis of on-target sgRNA efficiency can be predicted based on available scores and/or investigated experimentally by analysis of editing frequency. In addition to the numerous sgRNA efficiency prediction scores, CRISPOR offers automated primer design to facilitate polymerase chain reaction (PCR) amplification of regions for deep sequencing analysis to quantitate editing frequency. This involves PCR amplification of sequences flanking the DSB site for a given sgRNA. Similarly, CRISPOR offers computational prediction of off-target sites as well as PCR primers for deep sequencing analysis of mutagenesis at these predicted sites.

CRISPOR can also be used for the design of gene-targeted pooled screens by including exonic regions for sgRNA design as well as saturating mutagenesis screens. Alternatively, a list of gene names can be input into CRISPOR (“CRISPOR Batch”) to aid in the design of large scale gene-targeted libraries, which includes the inclusion of non-targeting controls. This analysis includes automated design of the required oligonucleotides for library cloning. Saturating mutagenesis involves utilizing all PAM-restricted^11,12,18^ sgRNA within a given region(s) in a pooled screening format to identify functional sequences^11,12,18^. Saturating mutagenesis can be used to analyze coding and non-coding elements in the genome or a combination of the two. Screen resolution is a function of PAM frequency and can be enhanced by PAM choice and/or combination of nucleases with unique PAM sequences^12^. CRISPOR can be utilized to design saturating mutagenesis libraries by simply selecting all sgRNA within the inputted region(s). This analysis includes automated design of the required oligonucleotides for library cloning. It is particularly important to consider off-target prediction scores offered by CRISPOR for saturating mutagenesis screens as repetitive sequences can confound screen results^12^. sgRNAs with high probability of off-target mutagenesis can be excluded at the library design stage or can be appropriately handled at the analysis stage (Box 1).

#### Analysis of deep sequencing from arrayed or pooled sgRNA experiments using CRISPResso

CRISPResso offers the ability to quantitate and visualize genome editing outcomes at individual loci^10^. CRISPResso provides a variety of features to offer users the opportunity to optimize analysis of sequencing data (Box 2). This analysis can be performed using the CRISPResso webtool or command line version (Table 2). CRISPResso analysis of an individual locus requires PCR amplification of the sequences flanking the genomic position of the DSB for a given sgRNA. The resulting deep sequencing FASTQ file can be analyzed by CRISPResso to quantitate the indel spectrum as well as visualize individual alleles (Fig. 4). This analysis can be performed when targeting coding (Fig. 4a-d) or non-coding sequences. When targeting coding (exonic) sequence, CRISPResso can determine the frequency of inframe and out-of-frame (frameshift) mutations produced (Fig. 4d). This type of analysis may suggest toxicity upon gene knockout by identifying an increased frequency of in-frame mutations^12^. Two separate CRISPResso analyses with the same amplicon can be directly compared using the CRISPRessoCompare tool (Table 2). CRISPRessoCompare is useful in situations such as comparing “treated” and “untreated” groups as well as to compare different experimental conditions. It can also be used to compare indel distributions created by two different sgRNA within the same region/amplicon^11^.

CRISPResso analysis can be extended from a single amplicon to multiple amplicons using the CRISPRessoPooled tool (Table 2). This is useful for individual locus experiments that require multiple amplicons for analysis of the full region or when multiple genes are disrupted. The CRISPResso suite can also be used to analyze whole genome sequencing (WGS) data through CRISPRessoWGS. This requires pre-aligned WGS data in BAM format, which can be created using publicly available aligners (e.g., Bowtie2^75^ or Burrows-Wheeler Aligner^76,77^). Similar to CRISPRessoCompare, two analyses using either CRISPRessoPooled or CRISPRessoWGS can be directly compared using CRISPRessoPooledWGSCompare (Table 2). This can be particularly useful when using multiple amplicon sequencing data or WGS data to evaluate off-target cleavages by two different sgRNA to identify sgRNAs with lower off-target activity.

CRISPResso also offers the ability to analyze deep sequencing data from CRISPR pooled screens. Sequence/indel based analysis in a pooled screening format is confounded by non-edited reads from cells containing sgRNA targeting other regions/loci. Therefore, pooled screens are typically analyzed by enumeration of sgRNAs by PCR amplifying sequences containing the cloned sgRNA within the lentiviral construct that has been integrated into the cell's genome (using PCR primers specific to the lentiviral construct). Deep sequencing data generated using PCR primers specific to the lentiviral construct can be analyzed by CRISPRessoCount for sgRNA enumeration for the purposes of calculating enrichment and/or dropout of sgRNAs in differing experimental conditions^11,12^ (Fig. 2, 4e; Box 4).

**Fig. 2:**
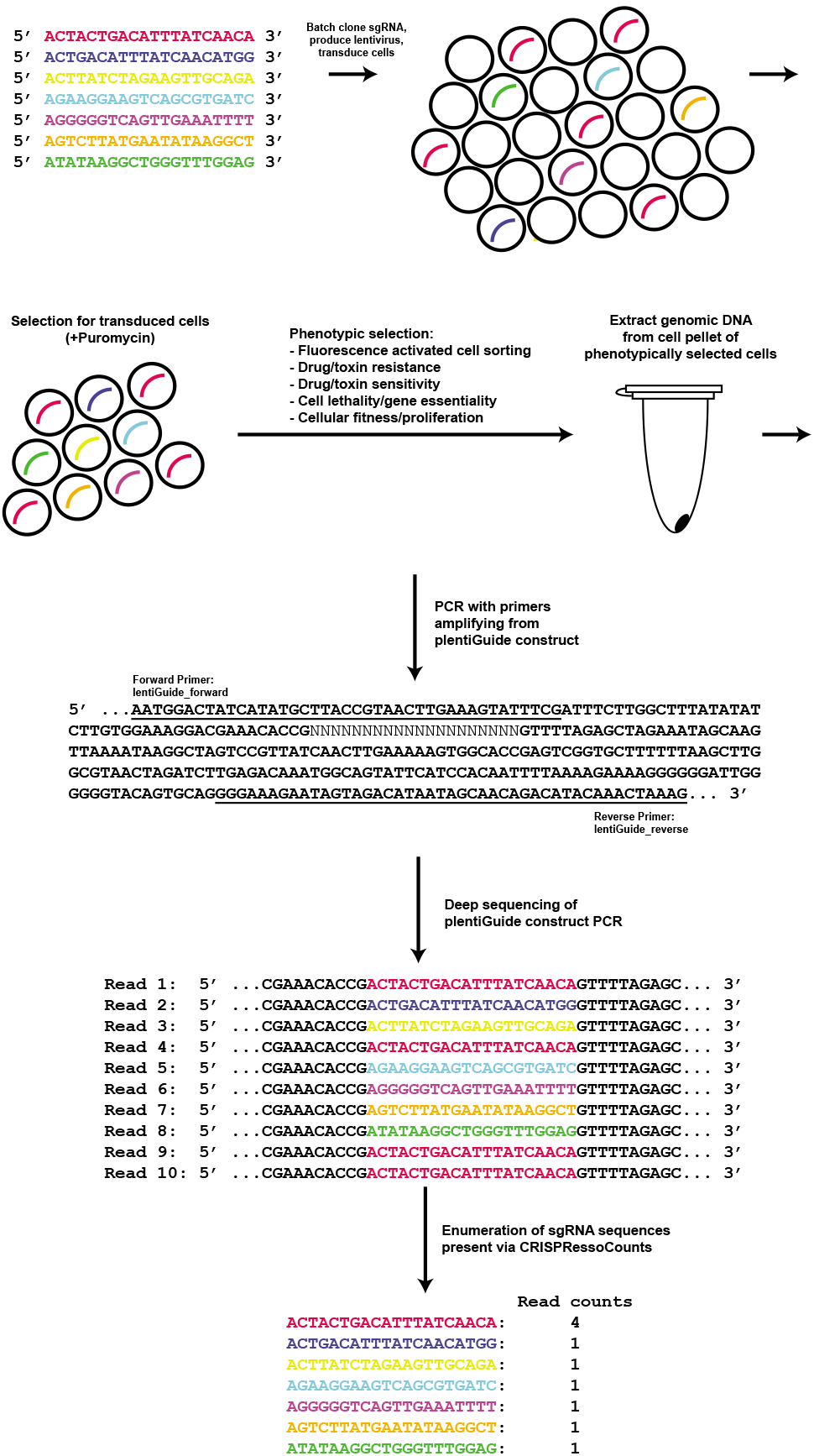
Schematic of a pooled genome editing experiment. Pooled genome editing experiments are performed by designing multiple sgRNA using CRISPOR. After designing the sgRNAs, they are batch cloned into pLentiGuide-puro, lentivirus produced, cells transduced at low multiplicity, and successful transductants are selected (successful transduction is indicated by curved lines). Phenotypic selection is performed (e.g., fluorescence activated cell sorting, drug/toxin resistance, drug/toxin sensitivity, cell lethality/gene essentiality, cellular fitness/proliferation). After conclusion of the experiment, cells are pelleted and genomic DNA is extracted. PCR primers specific (primer sequence is underlined) to the pLentiGuide-Puro construct are used to amplify regions flanking the cloned sgRNA sequence. Deep sequencing of the amplicon generated by construct-specific PCR is subsequently performed. sgRNAs present within the sample are enumerated by CRISPRessoCount.

#### Experimental designs and workflows

All CRISPR experiments must begin with target identification as described above, which may include a single locus, genome-wide targeting, and everything in between (Fig. 3). Arrayed experiments are useful when a target can be mutagenized by one sgRNA (e.g., a transcription factor binding motif, gene-knockout via exon mutagenesis) (Fig. 1). For an arrayed experiment, the next step after target identification is sgRNA design using CRISPOR. It is essential to consider on- and off-target potential at this stage, which can be done using the on- and off-target prediction scores described above. Further detail regarding the sgRNA selection process is described in Box 1. After an sgRNA is selected, CRISPOR offers automated primer design to allow for deep sequencing of on- and off-target sites at the conclusion of the experiment. Notably, this allows for targeted evaluation of off-target mutagenesis while genome-wide approaches can be accomplished by previously published methods such as GUIDE-seq, Digenome-seq, BLISS, CIRCLE-seq, SITE-seq or HTGTS^60–66^. CRISPResso indel analysis can supplement other common analyses, such as gene expression or protein level changes. CRISPResso is useful to quantify editing frequency to demonstrate that editing has successfully occurred, demonstration of the full indel spectrum (substitutions, insertions, and deletions), and offers other analyses that may be applicable depending on the targeted genomic region (e.g., splice-site analysis, frameshift vs. in-frame analysis). This workflow for an arrayed experiment is summarized in Figures 1 and 3. A representative example of an arrayed experiment is shown in Figure 4a-d, which demonstrates successful mutagenesis of exon 2 of *BCL11A* for the purposes of gene knockout. See Box 3 for further discussion of arrayed experiments.

**Fig. 3:**
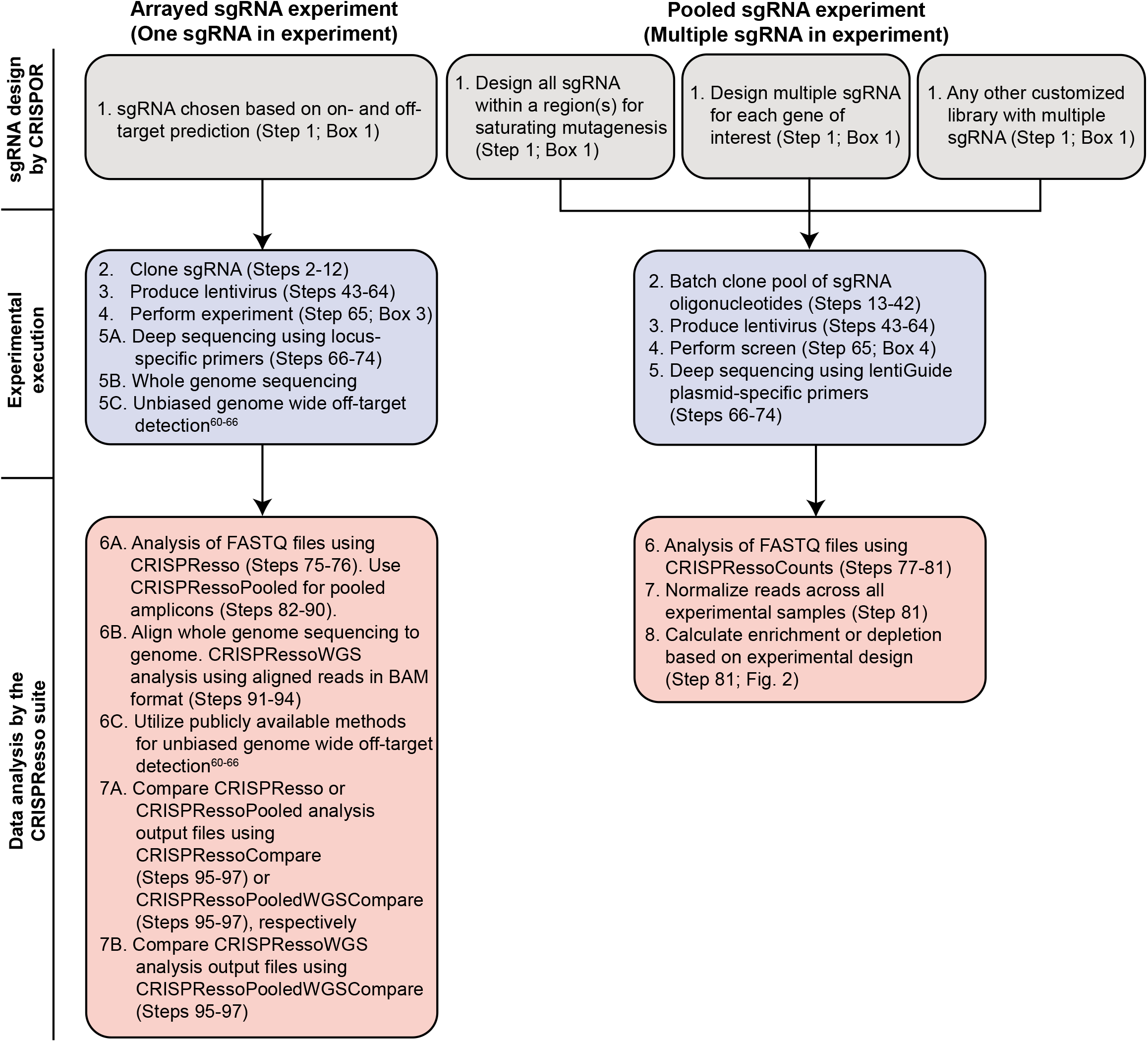
Design, experimental execution, and data analysis workflows for arrayed and pooled genome editing experiments. sgRNA design steps by CRISPOR are shown in gray, experimental execution steps are shown in blue, and data analysis steps by CRISPResso are shown in red.

Pooled screen experiments are useful when a target(s) requires >1 sgRNA for mutagenesis (Fig. 2). Both arrayed and pooled experiments are described in this protocol as arrayed experiments are typically required to validate the results of pooled experiments. As with arrayed experiments, the first step is target identification. This can include targeting a single locus with multiple sgRNAs (e.g., saturating^11,12,18^ mutagenesis (so-called tiling sgRNA) experiments)^11,12,18^, targeting multiple loci by saturating mutagenesis (e.g., saturating mutagenesis of multiple DNase hypersensitive sites)^11,12^, or gene-targeted pooled screens (e.g., targeting multiple genes with multiple sgRNAs per gene)^14–17^. In addition, libraries combining gene-based targeting and saturating mutagenesis approaches can also be designed. CRISPOR offers the ability to design saturating mutagenesis screens including designing oligonucleotides ready for cloning as described in this protocol. Furthermore, CRISPOR offers a streamlined approach for designing gene targeted screens with similar automation of oligonucleotides ready for cloning. After successful cloning and execution of a pooled screen, CRISPRessoCount can be used to enumerate sgRNAs present. Workflows for a pooled experiment are summarized in Figures 2-3. An example of a pooled experiment analyzed by CRISPRessoCount is shown in Figure 4e. Specifically, Figure 4e demonstrates saturating mutagenesis of a DNase hypersensitive site within an enhancer of *BCL11A*. It is important to note that validation of saturating mutagenesis screens typically involves using the top scoring sgRNAs in arrayed experiments while validation of gene-targeted screens is commonly performed using unique sgRNAs (not included in the library design) targeting the identified genes in arrayed experiments. Targeted genomic deletions using unique sgRNAs can also be used to validate saturating mutagenesis experiments. See “Anticipated Results” for further discussion of pooled screen data and Box 4 for additional details on pooled experiments.

**Fig. 4:**
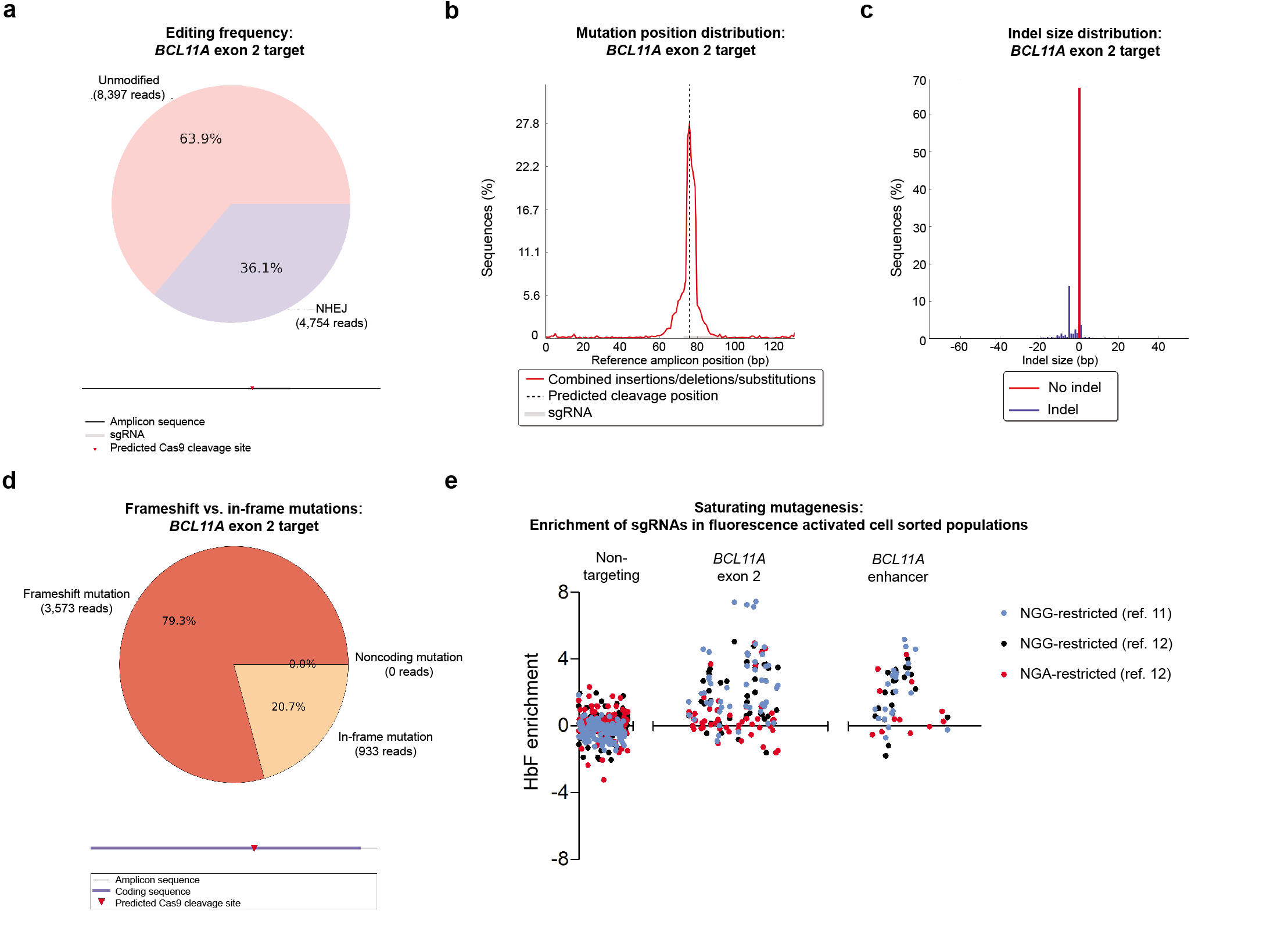
Locus-specific deep sequencing analysis of coding and non-coding targeting by CRISPResso. **a**, Frequency distribution of alleles with indels (shown in blue) and without indels (shown in red) for an sgRNA *BCL11A* exon 2. b, All reads with sequence modifications (insertions, deletions, and substitutions) with modification mapped to position within the *BCL11A* exon 2 reference amplicon. The vertical dashed line indicates the position of predicted Cas9 cleavage. The position of the sgRNA is shown in gray. c, Distribution of indel sizes when targeting *BCL11A* exon 2. Percentage of unmodified sequences is shown in red and percentages of modified sequences are shown in blue. d, Frameshift analysis of *BCL11A* exon 2 coding sequence targeted reads. Frameshift mutations are shown in red and in-frame mutations are shown in tan. e, sgRNA enrichment based on analysis of fetal hemoglobin (HbF) levels when performing saturating mutagenesis of *BCL11A* exon 2 and the functional core of the *BCL11A* enhancer using NGG-and NGA-restricted sgRNA using data from two previously published studies^11,12^. Non-targeting sgRNA are pseudo-mapped with 5 bp spacing.

## MATERIALS

REAGENTS

### Cloning/transformation

E. cloni 10G ELITE electrocompentent cells with recovery medium (Lucigen, cat. no. 60052)

Endura electrocompetent cells with recovery medium (Lucigen, cat. no. 60242)

Gibson Assembly Master Mix (New England Biolabs, cat. no. E2611S)

Gene Pulser/MicroPulser Electroporation Cuvettes, 0.1 cm gap (Biorad, cat. no. 1652089)

Corning Untreated 245mm Square BioAssay Dishes (Fisher Scientific, cat. no. 431111)

Fast Digest Esp3I (ThermoFisher, cat. no. FD0454)

NEB Stable Competent E. coli (High Efficiency) (New England Biolabs, cat. no. C3040H)

Pyrex solid glass beads (Fisher Scientific, cat. no. 11-312-10B)

LB broth base (ThermoFisher, cat. no. 12780052)

BD bacto dehydrated agar (Fisher Scientific, cat. no. DF0140-01-0)

Corning Falcon Bacteriological Petri Dishes with Lid (Fisher Scientific, cat. no. 08-757-100D) Thermosensitive alkaline phosphatase (TSAP) (Promega, cat. no. M9910)

Nuclease-Free Water (Fisher Scientific, AM9937)

Falcon Cell Scraper with 40cm Handle and 3.0cm Blade (Corning, cat. no. 353087)

T4 polynucleotide kinase (New England Biolabs, cat. no. M0201S)

Ampicillin sodium salt (Sigma, cat. no. A9518)

S.O.C. Medium (ThermoFisher, cat. no. 15544034)

Quick Ligation Kit (New England Biolabs, cat. no. M2200S)

Individual oligonucleotides (e.g., Integrated DNA Technologies, Bio-Rad)

Oligonucleotide pool synthesis (e.g., CustomArray Inc., Twist Bioscience)

QIAquick PCR purification kit (Qiagen, cat. no. 28104)

### Gel electrophoresis

SYBR Safe DNA Gel Stain (ThermoFisher, cat. no. S33102)

VWR Life Science AMRESCO Agarose I (VWR, cat. no. 97062-250)

1 Kb Plus DNA Ladder (ThermoFisher, cat. no. 10787018)

50x TAE buffer (Boston Bioproducts, cat. no. BM-250)

### PCR

Phusion Hot Start Flex DNA Polymerase (New England Biolabs, cat. no. M0535S)

Q5 High-Fidelity DNA Polymerase (New England Biolabs, cat. no. M0491S)

Herculase II Fusion DNA Polymerase (Agilent Genomics, cat. no. 600675)

Dimethyl sulfoxide (Sigma, cat. no. D8418)

QIAquick Gel Extraction Kit (Qiagen, cat. no. 28704)

MinElute PCR Purification Kit (Qiagen, cat. no. 28004)

Qubit dsDNA HS Assay Kit (ThermoFisher, cat. no. Q32854)

DNeasy Blood & Tissue Kit (Qiagen, cat. no. 69504)

### Plasmids/plasmid preparation

lentiGuide-Puro (Addgene plasmid ID: 52963)^19^

lentiCas9-Blast (Addgene plasmid ID: 52962)^19^

pXPR_011 (Addgene plasmid ID: 59702)^46^

lenti-Cas9-VQR-Blast (Addgene plasmid ID: 87155)^12^

lenti-Cas9-VQR-GFP_activity_reporter (Addgene plasmid ID: 87156)^12^

Qiagen Plasmid Maxi Kit (Qiagen, cat. no. 12163)

AccuPrep Plasmid Mini Extraction Kit (Bioneer, cat. no. K-3030)

### Lentivirus Production

pCMV-VSV-G (Addgene plasmid ID: 8454)

psPAX2 (Addgene plasmid ID: 12260)

Polyethylenimine, Branched (Sigma, cat no. 408727)

Sucrose (Sigma, cat no. S0389)

Steriflip-HV, 0.45 µm, PVDF, radio-sterilized (Millipore, cat. no SE1M003M00)

Stericup-GP, 0.22 µm, polyethersulfone, 500 mL, radio-sterilized (Millipore, cat no. SCGPU05RE)

Phosphate buffered saline 1x, w/o calcium, magnesium (Lonza, cat. no 17-516F)

DMEM (Life Technologies, cat no. 11995-073)

Ultracentrifuge Tube, Thinwall, Polypropylene, 38.5 mL, 25 x 89 mm (Beckman Coulter, cat no. 326823) Falcon 50mL Conical Centrifuge Tubes (Fisher Scientific, cat no. 14-432-22)

Corning TC-Treated Culture Dishes, 15cm, round (Fisher, cat no. 08-772-24)

Penicillin-streptomycin (Life Technologies, cat no. 15140122)

HEK293FT cells (Thermo Fisher Scientific, cat. no. R70007)

Lenti-X qRT-PCR Titration Kit (Takara Bio USA, Inc., cat. no. 631235)

## REAGENT SETUP

**Preparing sgRNA cloning, sequencing, and primer oligonucleotides:** Dissolve oligonucleotides in nuclease-free water at a concentration of 100 µM. Store at -20 °C for up to two years. This does not require protection from light.

**TAE electrophoresis solution:** Dilute 50x TAE buffer stock with dH2O for a 1x working solution. Store at room temperature (25 °C) for up to 6 months. This does not require protection from light.

**Ampicillin solution:** Dissolve ampicillin in a one-to-one mix of pure ethanol and dH2O to a final concentration of 100 mg/mL. Store at -20^o^C for up to 4-6 months. This does require protection from light.

**HEK293FT cell culture medium:** DMEM with glucose/sodium pyruvate supplemented with 10% (vol/vol) fetal bovine serum (FBS) and 1% (vol/vol) penicillin–streptomycin. Sterilize with 0.22 µm filter store it at 4°C for up to 4 weeks. This does not require protection from light.

CAUTION: The HEK293 cells used should be regularly checked to ensure that they are not infected by mycoplasma.

**Polyethylenimine solution:** Dissolve in nuclease free water at 10 µg/µL with pH 7.4. This can be stored at 4 ^o^C for years and does not require protection from light.

**20% sucrose solution:** Dissolve sucrose in PBS to create 20% (wt/vol) solution by mass. Sterilize with 0.22 µm filter store it at 4 °C for up to 1 year. This does not require protection from light.

## EQUIPMENT

Sequence analysis software (e.g. free software such as ApE, SerialCloner or other such as Thermo Scientific Vector NTI, Lasergene DNAstar, CLC Main Workbench, or MacVector)

Electroporation system (e.g., Bio-Rad’s Gene Pulser MXcel, Gene Pulser Xcell, and MicroPulser Electroporators)

MiSeq or HiSeq sequencing system (Illumina) or equivalent sequencing platform

Qubit Fluorometer (Thermo Fisher Scientific)

UV-Vis spectrophotometer (NanoDrop)

Gel visualization system (Alpha Innotech)

Thermocycler (Bio-Rad)

UV-light transilluminator and UV-filter face-mask

CAUTION: Wear gloves, lab coat and face shield to avoid harm to eyes and skin caused by UV light.

VWR Microcentrifuge Tubes, Polypropylene (VWR, cat. no. 87003-294)

8-Strip PCR Tubes, 0.2mL, (Fisher Scientific, cat. no. 14-222-250)

Sterile Culture Tube with Attached Dual Position Cap (VWR, cat. no. 89497-812)

Ultracentrifuge (Beckman Coulter)

SW32Ti rotor with compatible swinging buckets (Beckman Coulter, cat no. 369694)

## COMPUTING EQUIPMENT

Computer with at least 8 GB of RAM, 1 TB of disk space and a recent CPU with 2 or more cores (command line and Docker version only). Windows (Docker only), OSX, and Linux platforms are supported.

Anaconda Python 2.7: https://www.continuum.io/downloads

Docker: https://www.docker.com/community-edition#/download

CRISPOR command line software: http://crispor.org/downloads

CRISPResso command line software: https://github.com/lucapinello/CRISPResso

FASTQC: http://www.bioinformatics.babraham.ac.uk/projects/fastqc/

Refer to Steps 1C (CRISPOR) 75-97 (CRISPResso) for detailed installation instructions and troubleshooting.

**Computational skills necessary:** For the command line and Docker versions of this protocol, it is required to have a basic understanding of how to execute a command in a terminal and how to navigate a Unix-based filesystem. For the online version, no particular skills are required.

## PROTOCOL

### sgRNA design using CRISPOR | Timing 1-4 h

1) The CRISPOR webtool allows for the design of sgRNAs from a single genomic locus (option A). sgRNAs can be selected manually from the CRISPOR output using on- and off-target predictions (Box 1). Alternatively, the “CRISPOR Batch” gene targeting assistant webtool offers gene-targeted pooled library generation from a gene list (option B). If multiple genomic loci are required for sgRNA design or command line is preferred, the command line version of CRISPOR can be used (option C). CRISPOR does not offer the ability to design repair templates for HDR experiments. If the experimental design requires a repair template, other protocols can provide further instruction^58^.

**(A) sgRNA design for arrayed and pooled experiments using the CRISPOR webtool | Timing 1-4 h**

i) Use the CRISPOR webtool: http://crispor-beta.tefor.net

ii) Input target DNA sequence. The webtool requires sequence input (<1 kilobase) or genomic coordinates. The input sequence must be a genome sequence as opposed to a cDNA, which can include sequence that is not in the genome due to splicing. You can obtain genomic sequences

using a website such as the UCSC Genome Browser (https://genome.ucsc.edu, click “View - DNA”) or Ensembl (http://www.ensembl.org, click “Export data– Text”). From the UCSC Genome Browser, the current sequence in view can be sent directly to CRISPOR via the menu entry “View– In external tools”

? TROUBLESHOOTING

iii) Select the relevant assembly (e.g. hg19, mm9) and PAM sequence for the relevant nuclease (e.g., NGG for *S. pyogenes* Cas9).

CRITICAL STEP: It is important to pick the appropriate assembly to be consistent with the genomic coordinates of the DNA sequences provided in Step 1A(ii) and it also essential for accurate prediction of off-target sites.

iv) Run CRISPOR analysis to identify optimal sgRNA based on double-strand break position as well as on- and off-target prediction (Box 1). Select the “Cloning/PCR primers” link underneath each sgRNA sequence for automated primer design for future indel analysis by CRISPResso at on- and off-target loci.

If you are conducting a saturating mutagenesis screen, the list of the oligonucleotide pool sequences for Gibson assembly to be ordered, sequencing primers for validation, sequencing amplicons for CRISPResso, and the full list of guide sequences can be downloaded by following the link “Saturating Mutagenesis Assistant” at the top of the CRISPOR output page. Alternatively, the full list of sgRNA sequences for saturating mutagenesis can also be found by following the “Cloning/PCR primers” link under each sgRNA sequence and then following the link provided under the “Saturating mutagenesis using all guides” heading. This webtool allows for filtering based on a minimum specificity score for off-target concerns and/or the Doench 2016 score (Box 1) for on-target activity. It is also possible to filter based on these or other characteristics using Excel or command line procedures as illustrated in Step 1C xvi). The four files provided by the Saturating Mutagenesis Assistant can be downloaded by selecting from the dropdown menu adjacent to “output file”. The outputs are described in Table 3. If you are conducting a gene-targeted screen, it is possible to provide CRISPOR with the relevant exonic sequences. Alternatively, a list of gene names can be provided using the “CRISPOR Batch” gene targeting assistant feature (see Step 1B).

**Table 3.**
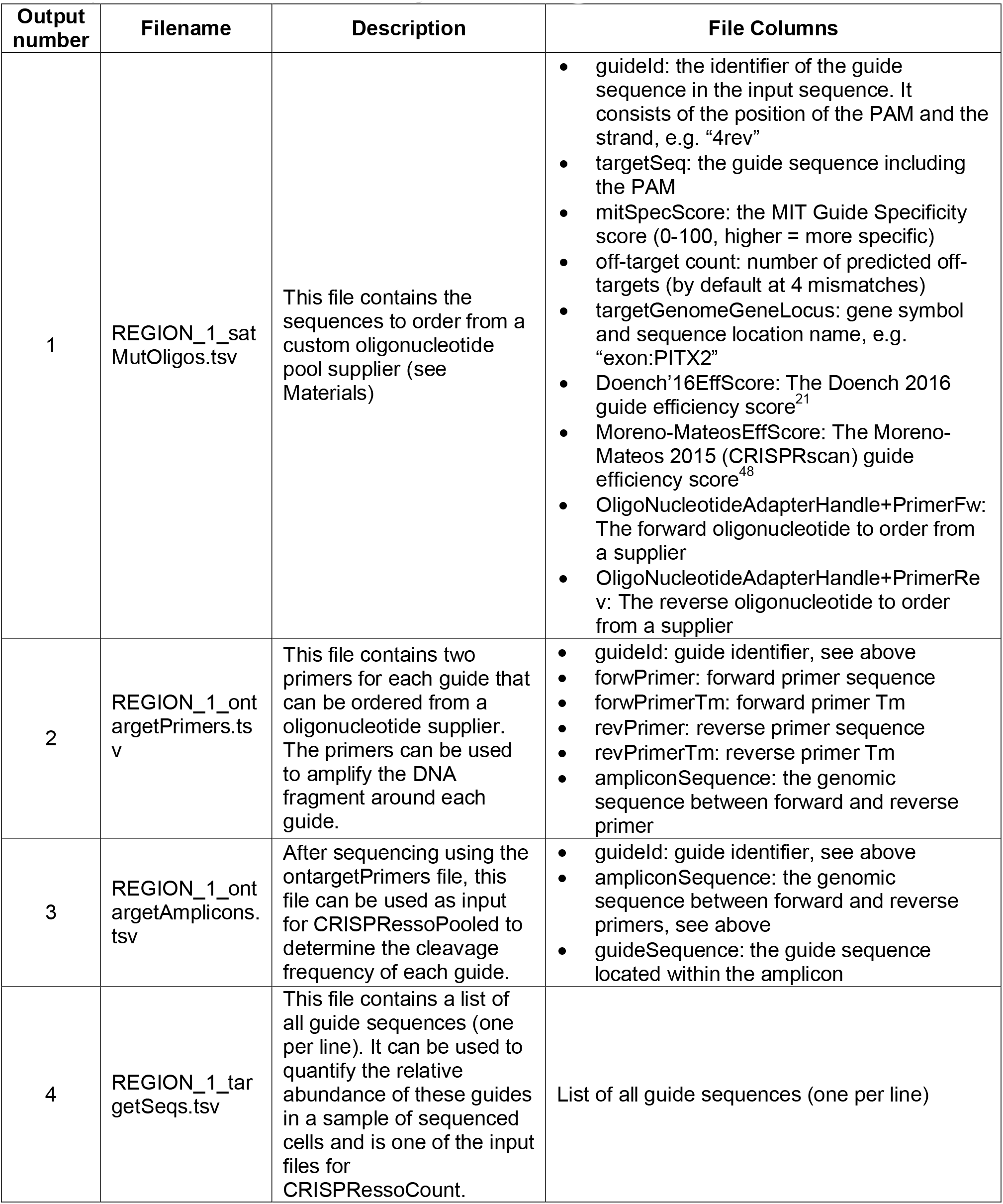
|Four output files from CRISPOR sgRNA design analysis

CRITICAL STEP: CRISPOR allows for PCR amplicon lengths of 50-600 bp and the melting temperature (Tm) is calculated for each primer pair. Amplicon length should be selected based on deep sequencing read length. 100-200 bp amplicons are reasonable for paired-end deep sequencing with 75 or 150 bp reads, respectively. Because sequencing quality is lower towards the end of the reads, there must be some overlap of read pairs to allow reliable merging. For example, for a 200 bp amplicon it is suggested to use 125-150 bp reads in order to have sufficient overlap. Some considerations for determining amplicon length include the cost of sequencing for a given read length, the size in base pairs of the region to be sequenced, and the length of reliable sequencing (due to unreliable sequence at read ends). If an amplicon length is too short, it may result in inadequate coverage of the target region as well as missing larger indels. In contrast, if the amplicon is longer than twice the read length, no overlap will exist between reads making the merging steps of paired end reads impossible.

CRITICAL STEP: When designing a pooled screen, it is important to consider the inclusion of positive and negative controls (e.g., non-targeting or safe harbor targeting sgRNA) in the library design. It is important to note that non-targeting sgRNAs are often genome build-, species-and PAM-specific. For example, a non-targeting sgRNA can be designed for human hg19 reference genome using the NGG PAM. Non-targeting sgRNAs can be created by generating random sgRNA sequences and analyzing their on- and off-target potential, which can be automated by previously published tools^12^. In addition to ensuring no perfect matches in the genome restricted by the relevant PAM sequence, it is also important to ensure limited potential for cleavage at loci with few mismatches (similar to minimization off-target potential for targeted sgRNA). Safe harbor (e.g., *AAVS1* locus) targeted sgRNA can be designed using the instructions found in Step 1A or 1C. There is no strict requirement for the number of positive and negative controls to include in a pooled experiment. It is reasonable for both positive and negative controls to each comprise 1-5% of the total number of sgRNAs in the library (2-10% of the library comprising both positive and negative controls).

**(B) Gene-targeted library design generated from a gene list using the CRISPOR Batch gene targeting assistant webtool | Timing 1-4 h**

i) The “CRISPOR Batch” gene targeting assistant generates gene-targeted libraries from a gene list using sgRNA sequences from previously validated genome-wide libraries^19,21^. This feature is useful when designing a gene-targeted library that is not genome-wide, but rather contains a small subset of gene targets (e.g., library targeting all kinases in the genome). If a gene list is provided, the number of sgRNA per gene and the source genome-wide library can be selected. In addition, a user-set number of non-targeting sgRNA can be added to the library. Finally, the list of oligonucleotide pool sequences for Gibson assembly cloning can be downloaded. To generate a gene-targeted library using CRISPOR Batch gene targeting assistant, go to: http://crispor-beta.tefor.net

ii) Select the source genome-wide library for the extraction of sgRNA sequences using the dropdown menu adjacent to “Lentiviral screen library”. References are provided for each library to assist with appropriate source library selection.

iii) Select the desired “number of guides per gene” (maximum of 6 sgRNAs/gene) using the dropdown menu. Note that some libraries have fewer than 6 sgRNA per gene. If more sgRNAs are requested than are available for a given gene with the library, the maximum number of sgRNA sequences available within the library will be output instead.

iv) Enter the “number of non-targeting control guides” to include in the library (maximum 1,000). It is reasonable for non-targeting sgRNAs to comprise 1-5% of the total number of sgRNAs in the library.

v) Select a barcode to allow for library preparation as described in Step 13. Refer to Step 13 for further discussion of barcodes.

vi) Copy and paste gene list into the entry field. Genes should be entered as gene symbols, Entrez Gene IDs, or Refseq IDs with only one entry per line. This entry is case insensitive due to libraries already being species specific. If an entry is not found in the database of genome-wide sgRNAs, sgRNAs for the entered gene will display a warning and will not be included in the output.

? TROUBLESHOOTING

vii) Click “Submit” for gene-targeted library generation and download the output library.

**(C) sgRNA design using command line CRISPOR | Timing 1-4 h**

i) Installation of Docker: Docker is a virtualization technology that allows packaging software with all dependencies into files called containers to be executed on either Windows, Linux or OSX. This allows creating and distributing a “frozen” version of the software that will always run independent of updates or changes to libraries or the required dependencies on the host machine. In this case, you only need to install Docker and do not need to install any dependencies. Download and install Docker from this link: https://docs.docker.com/engine/installation/.

ii) Sharing disk volumes with Docker: By default, Docker containers cannot share data with the machine where they run. For this reason, it is necessary to map local folders to folders inside the container to allow the container to read the input files with the data to process and to write the output files in a folder on the disk of the local machine. First, it is necessary to check if Docker has permissions to access the disk(s) on your machine where analysis data is to be stored and processed. By default, any subfolder within your home directory is automatically shared; however, this behavior may change with future versions of Docker. Check that that the drive(s) you want to be available to the container is(are) selected in the Settings…/ Shared Drives panel (See Supplementary Fig. 1).

To map a local folder (i.e. a folder on the local disk) to a container folder, Docker has a special option with this syntax:

~~~
-v local_folder:container_folder
~~~

You can specify this option multiple times if it is necessary to map more than one folder. In this example, the folders to map are all subfolders of the home folder of the user “user”, i.e. /home/user/. In addition, some of the Docker commands will be used with the option -w /DATA to specify the working directory (i.e. where the command will be executed) to allow for the use of relative paths and to shorten the commands.

iii) Allocation of memory for Docker containers: It is necessary to allocate enough memory to the container. The default assigned by Docker may depend on the version and on the machine upon which it is run. To run the CRISPOR/CRISPResso containers, we suggest assigning Docker 6-8GB of RAM. For analysis involving large sequences or many sites, it may be necessary to increase the amount of allocated memory under the Settings…/ Advanced panel (See Supplementary Fig. 2).

CRITICAL STEP: The execution may halt with an error if enough memory is not allocated to the container.

? TROUBLESHOOTING

iv) Type the following command to download the latest version of the container for this protocol:

~~~
docker pull lucapinello/crispor_crispresso_nat_prot
~~~

v) Verify that the container was downloaded successfully by running the command:

~~~
docker run lucapinello/crispor_crispresso_nat_prot crispor.py
~~~

vi) Create a folder to store the genomes to use and download a pre-indexed genome for CRISPOR. Obtain the CRISPOR assembly identifier of the genome (e.g. hg19 or mm9). If you are unsure, the full list of assemblies is available at http://crispor.tefor.net/genomes/genomeInfo.all.tab.

vii) Execute the following commands. Here we are assuming that the user will store the pre-indexed genomes in the folder:

~~~
/home/user/crispor_genomes
~~~

~~~
mkdir -p /home/user/crispor_genomes
~~~

~~~
docker run -v /home/user/crispor_genomes:/crisporWebsite/genomes lucapinello/crispor_crispresso_nat_prot downloadGenome hg19 /crisporWebsite/genomes
~~~

viii) Prepare genomic input sequences in FASTA format (see a description of the format here https://www.ncbi.nlm.nih.gov/blast/fasta.shtml).

CRITICAL STEP: It is important to pick the appropriate assembly to be consistent with the genomic coordinates of the DNA sequences provided in Step 1C(vi) and it also essential for accurate prediction of off-target sites.

ix) Generate FASTA file with exonic sequences (optional): If you have a list of genes and you want to target their exons, use an internet browser and go to the UCSC Genome Browser page http://genome.ucsc.edu/cgi-bin/hgTables

x) Set the following options (adjust entries as appropriate for each experiment and refer to the description of each item in the “Table Browser” provided directly below the “Table Browser” under the heading “Using the Table Browser”). Of note, the “Table” parameter will change depending on the chosen “Track”:

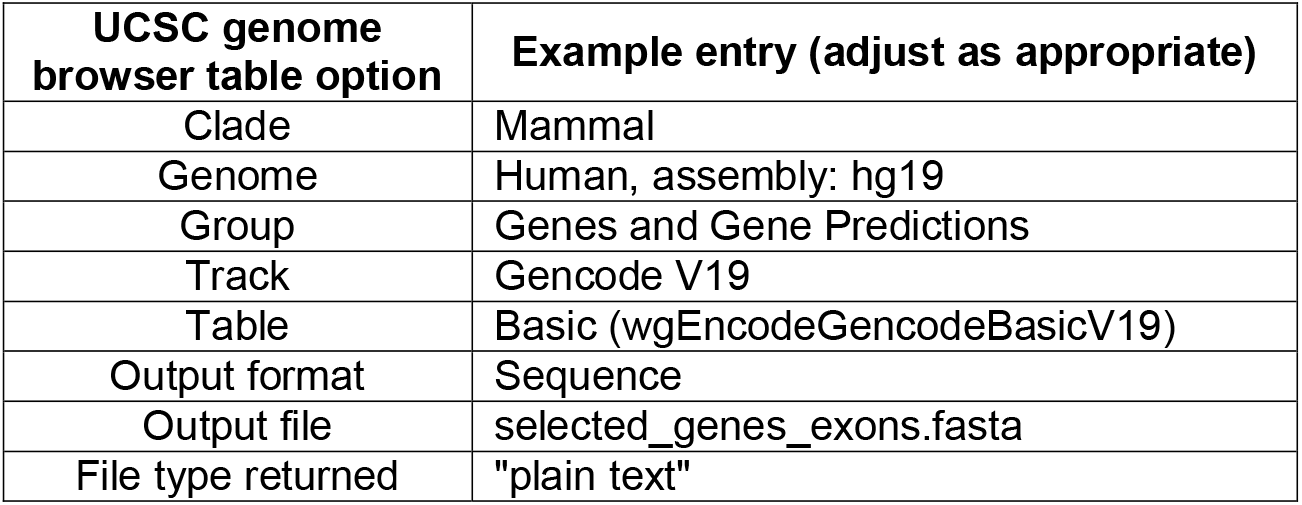

xi) Press the button “paste list” and enter the list of transcript identifiers for each gene.

xii) Press the button “get output”. A new page will open.

xiii) Select the option “genomic” and press submit. A new page will open

xiv) Select “CDS: Exons and “One FASTA record per region (exon, intron, etc.)”

xv) Press the button: “get sequence” and save the file as selected_genes_exons.fasta

CRITICAL STEP: The “Send output to” Galaxy, GREAT, or GenomeSpace checkboxes are off by default; however, they will remain checked if you have ever checked any of them in the past. Make sure that none of the checkboxes are selected.

### Run CRISPOR analysis on single- or multi-FASTA input file | Timing 15 min

xvi) Run CRISPOR over your single- or multi-fasta input file. Here we assume this file is stored in

~~~
/home/user/crispor_data/crispor_input.fasta and contains a sequence with id >REGION_1.
~~~

~~~
docker run \
~~~

~~~
-v /home/user/crispor_data:/DATA \
~~~

~~~
-v /home/user/crispor_genomes:/crisporWebsite/genomes \
~~~

~~~
-w /DATA \
~~~

~~~
lucapinello/crispor_crispresso_nat_prot \
~~~

~~~
crispor.py hg19 crispor_input.fasta crispor_output.tsv–satMutDir=./
~~~

The sgRNA sequences will be written to the file crispor_output.tsv and 4 files for each sequence ID in the FASTA file will be created like in the web version into the folder `/home/user/crispor_data/` (REGION_1_satMutOligos.tsv,

REGION_1_ontargetPrimers.tsv, REGION_1_ontargetAmplicons.tsv, REGION_1_targetSeqs.tsv; see Table 3).

? TROUBLESHOOTING

### Selecting a subset of sgRNAs and primers for analysis | Timing 5 min

xvii) sgRNA(s) can be filtered based on a minimum specificity score for off-target concerns and/or the Doench 2016 score (Box 1) for on-target activity at the design stage (see Step 1A(iv)). If removal of any sgRNA(s) is desired after the library has been designed (e.g., due to high off-target potential), a subset of sgRNAs and their associated information as described in Table 3 can be selected. To select a subset of sgRNAs from the file REGION_1_satMutOligos.tsv as generated in Step 1A(iv) or 1C(xvi) (Table 3), select the lines corresponding to sgRNA with desired scores/attributes (see Box 1) and save them to a new file called REGION_1_satMutOligos_filtered.tsv. This operation can be performed using such programs as Excel or the *awk* utility. Filtering within Excel can be performed using the “Filter” functionality. Please refer to Excel support and online help solutions for guidance on filtering within Excel. Then run in order the following commands:

~~~
docker run -v /home/user/crispor_data/:/DATA -w /DATA lucapinello/crispor_crispresso_nat_prot bash -c “join -t $’\t’ - 1 1 -2 1 REGION_1_satMutOligos_filtered.tsv REGION_1_ontargetAmplicons.tsv -o 2.1,2.2,2.3 > CRISPRessoPooled_amplicons.tsv”
~~~

~~~
docker run -v /home/user/crispor_data/:/DATA -w /DATA lucapinello/crispor_crispresso_nat_prot bash -c “join -t $’\t’ -1 1 -2 1 REGION_1_satMutOligos_filtered.tsv REGION_1_ontargetAmplicons.tsv -o 2.3 | sed 1d > CRISPRessoCounts_sgRNA.tsv”
~~~

~~~
docker run -v /home/user/crispor_data/:/DATA -w /DATA lucapinello/crispor_crispresso_nat_prot bash -c “join -t $’\t’ -1 1 -2 1 REGION_1_satMutOligos_filtered.tsv REGION_1_ontargetPrimers.tsv -o 2.1,2.2,2.3,2.4,2.5,2.6,2.7 > REGION_1_ontargetPrimers_filtered.tsv”
~~~

This will create a filtered version of the files described in Table 3 to use to design amplicons and to perform CRISPResso analysis. If the input file to CRISPOR was a multi-fasta (Step 1C(xvi)), repeat this step for the other regions as appropriate.

### Synthesis and cloning of individual sgRNA into a lentiviral vector | Timing 3 d to obtain sequence-confirmed, cloned sgRNA lentiviral plasmid

2) When ordering individual oligonucleotides for sgRNA cloning, add “CACC” onto the sense strand of the sgRNA (5’ CACCNNNNNNNNNNNNNNNNNNNN 3’) and “AAAC” onto the antisense strand of the sgRNA (5’ AAACNNNNNNNNNNNNNNNNNNNN 3’). The two oligonucleotides are designed as reverse complements because they will be annealed together in Step 4. The “CACC” and “AAAC” sequence are required for cloning of the phosphorylated/annealed oligonucleotides into the Esp3l-digested/dephosphorylated lentiGuide-Puro plasmid (Steps 3-8). The addition of a “G” at the 5’ end of the sgRNA sequence can increase transcription from the U6 promoter (5’ CACCGNNNNNNNNNNNNNNNNNNNN 3’ and 5’ AAACNNNNNNNNNNNNNNNNNNNNC 3’); this is not required if the 5’ end of the sgRNA sequence is already a “G”. Individually resuspend lyophilized primer oligonucleotides at 100 µM in nuclease-free water. Using CRISPOR, the sequences for sgRNA cloning into lentiGuide-Puro can be downloaded by following the “Cloning / PCR primers” link below the sgRNA sequence. The ready-to-order oligonucleotide sequences are available from the sgRNA results table by selecting the “lentiGuide-Puro (Zhang lab)” plasmid under the heading of “U6 expression from an Addgene plasmid”.

PAUSE POINT: Resuspended oligonucleotides can be stored at -20 ^o^C for years.

3) Set up phosphorylation reaction:

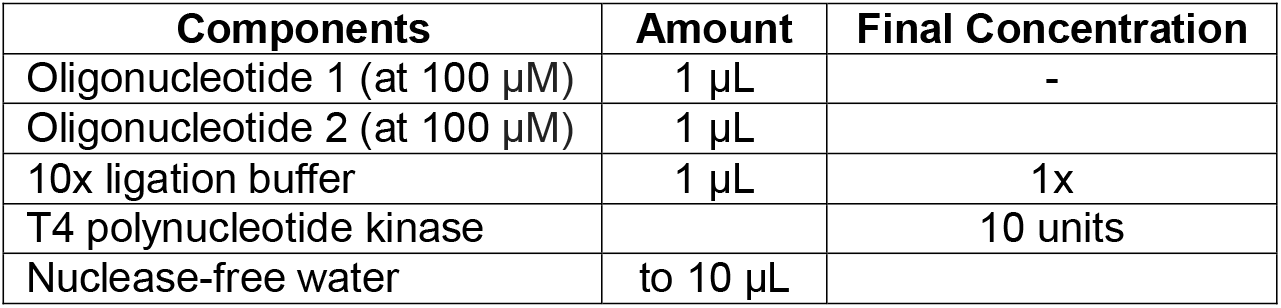

4) Perform phosphorylation and annealing of the oligonucleotides in a thermocycler:

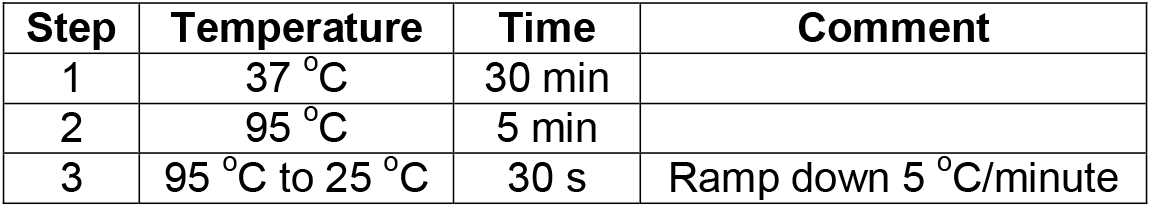

5) Digest sgRNA expressing plasmid (lentiGuide-Puro) with Esp3l (an isoschizomer of BsmBI) restriction enzyme for 15 minutes at 37 ^o^C.

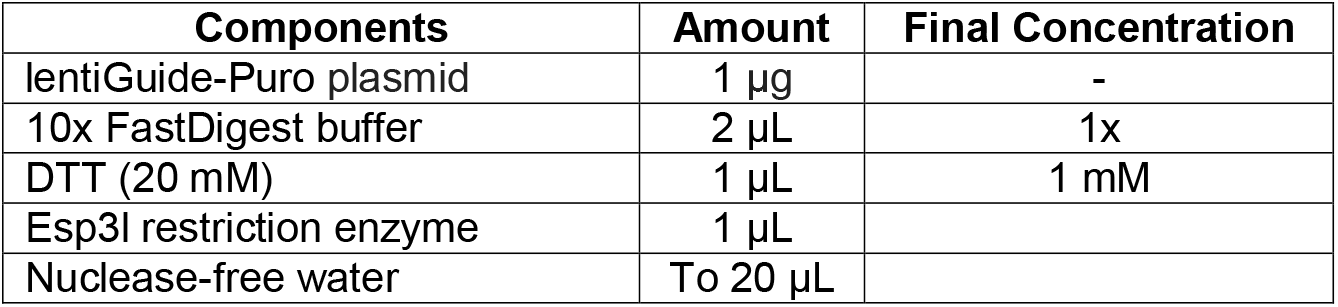

6) Add 1 µL of TSAP to dephosphorylate reaction for 15 minute-1 hour at 37 ^o^C following by heat inactivation for 15 minutes at 74 ^o^C.

7) Perform gel purification of the Esp3l-digested/dephosphorylated lentiGuide-Puro plasmid. See Materials for gel purification kit and follow manufacturer instructions. For further information on gel purification, refer to this previously published protocol^78^.

8) Perform ligation reaction for 5-15 minutes at room temperature (25 °C) using 30-70 ng of Esp3l-digested/dephosphorylated lentiGuide-Puro plasmid with the phosphorylated-annealed oligonucleotides produced in Steps 2-4 diluted 1:500 with nuclease-free water.

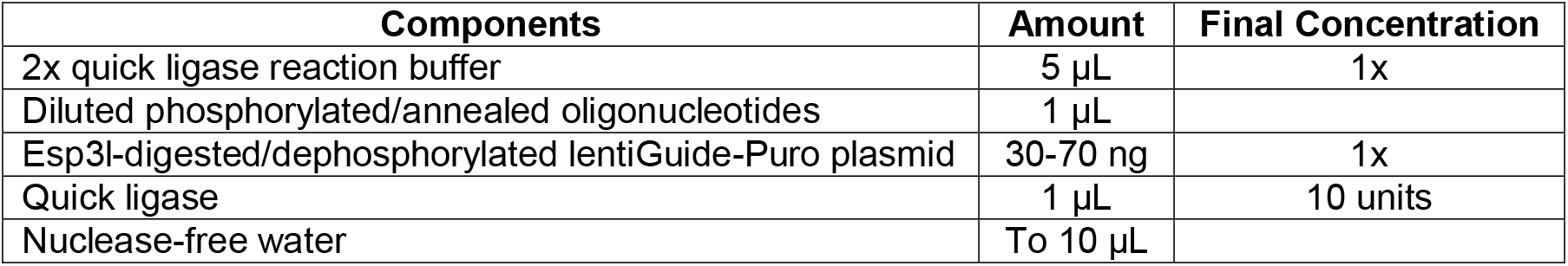

9) Heat-shock transform the ligation reaction in stable competent *E. coli*. Using recombination-deficient *E. coli* is important to minimize recombination events of repetitive elements in lentiviral plasmid. For more information about performing heat-shock transformation of competent *E. coli*, refer to these previously published protocols^79,80^.

10) Plate transformation on a 10-cm ampicillin-resistant (0.1 mg/mL ampicillin) agar plate. Incubate at 32 ^o^C for 14-16 hours. Growth at 32 ^o^C reduces recombination between the lentiviral long-terminal repeats.

CRITICAL STEP: It is recommended to incubate transformations at 32 ^o^C, but is not essential. Incubation at 37 ^o^C can also be performed.

11) Pick colonies (3-5 colonies is reasonable) and individually incubate each colony in 2-mL of ampicillin-resistant (0.1 mg/mL ampicillin) LB broth at 32 ^o^C for 14-16 hours with shaking at 250 rpm.

PAUSE POINT: Successfully cloned sgRNA plasmids can be stored long term (years) at -20 ^o^C prior to creating lentivirus (Steps 43-64).

CRITICAL STEP: It is recommended to incubate transformations at 32 ^o^C, but is not essential. Incubation at 37^o^C can also be performed. Growth at 32 ^o^C reduces recombination between the lentiviral long-terminal repeats.

? TROUBLESHOOTING

12) Perform mini-scale plasmid preparation of each 2-mL culture from Step 11 (see Materials for a miniscale plasmid preparation kit and follow manufacturer instructions). Sanger sequence each colony-derived plasmid with the U6 sequencing primer to identify correctly cloned plasmids^58^:

CGTAACTTGAAAGTATTTCGATTTCTTGGC

### Synthesis and cloning of pooled sgRNA libraries into a lentiviral vector | Timing 2-4 d to obtain cloned sgRNA lentiviral plasmid library

13) Use sgRNA’s identified/chosen in Step 1 to design full-length oligonucleotides (96-99 bp) flanked by barcodes and homologous sequence to the lentiGuide-Puro plasmid (Table 4). This step describes how to design the full-length oligonucleotides; however, CRISPOR automates this step for applications such as saturating mutagenesis designs (Steps 1A and 1C) and gene-targeted libraries generated from a gene list (Step 1B). Barcodes allow for oligonucleotides for multiple unique libraries to be synthesized on the same programmable microarray (Step 14), which reduces cost by avoiding purchase of multiple microarrays for multiple libraries. Specifically, the barcodes offer the ability for individual PCR-amplification of unique libraries from a single batch of oligonucleotides synthesized on the same microarray. The number of libraries that can be included on a single programmable microarray is limited by the microarray’s oligonucleotide capacity; however, there is no limit to the number of possible barcodes that can be utilized (and thus no limit to the number of libraries that can be generated from a single pool of oligonucleotides). Barcodes for 10 libraries are provided in Table 4. To create additional or new barcodes, generate 10-13 bp of sequence distinct from the sequences to be amplified and the other utilized barcodes. It is also important to ensure that the barcode results in a primer melting temperature (Tm) compatible with the lsPCR1 reaction (see Table 5 for examples). The homologous sequence to the lentiGuide-Puro plasmid is required because batch sgRNA library cloning is performed using Gibson assembly (Steps 23-24), which relies on homologous flanking sequence for successful cloning.

**Table 4.**
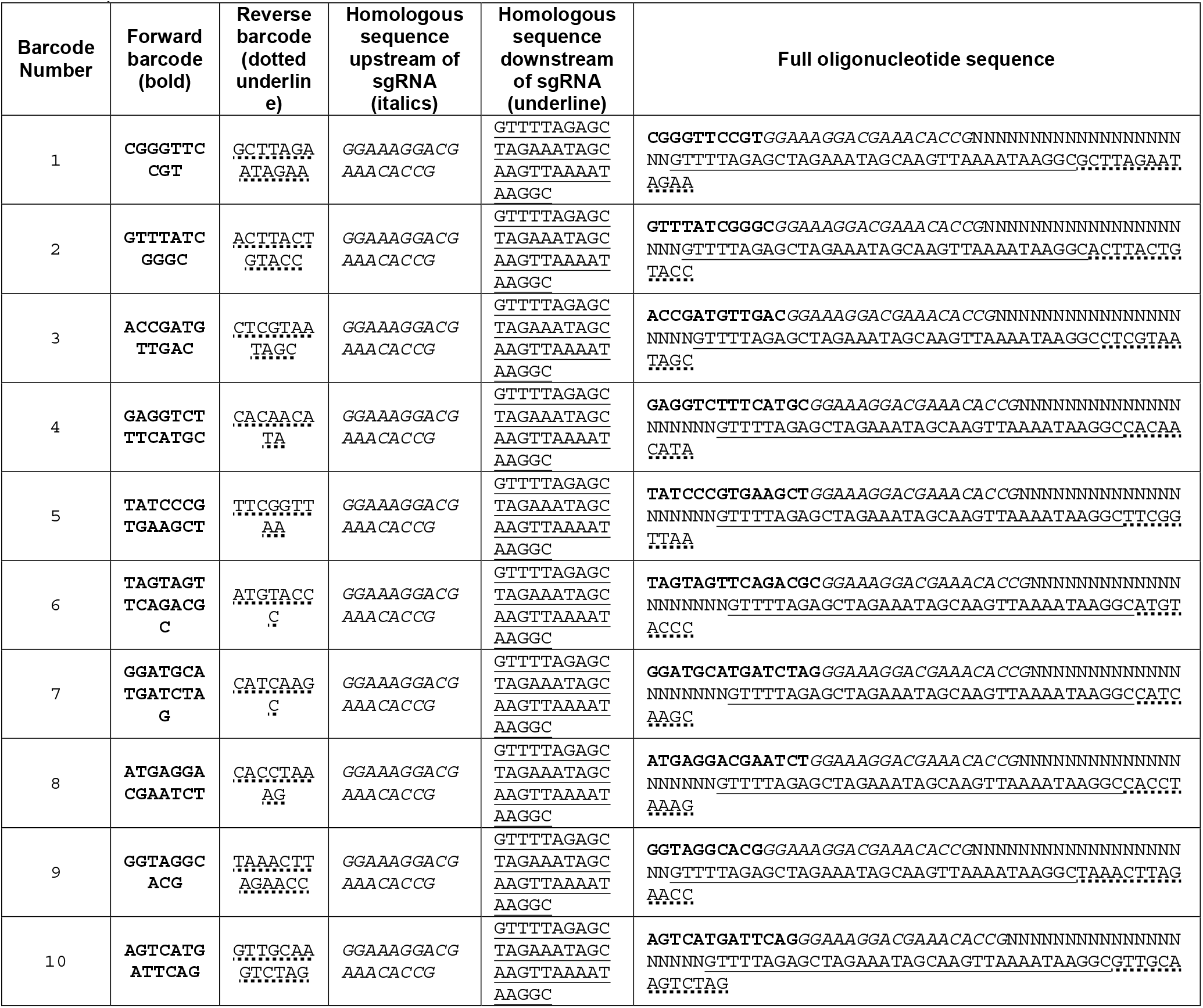
|Barcoding strategy for oligonucleotide pool synthesis. The sgRNA sequence is denoted with N’s

**Table 5.**
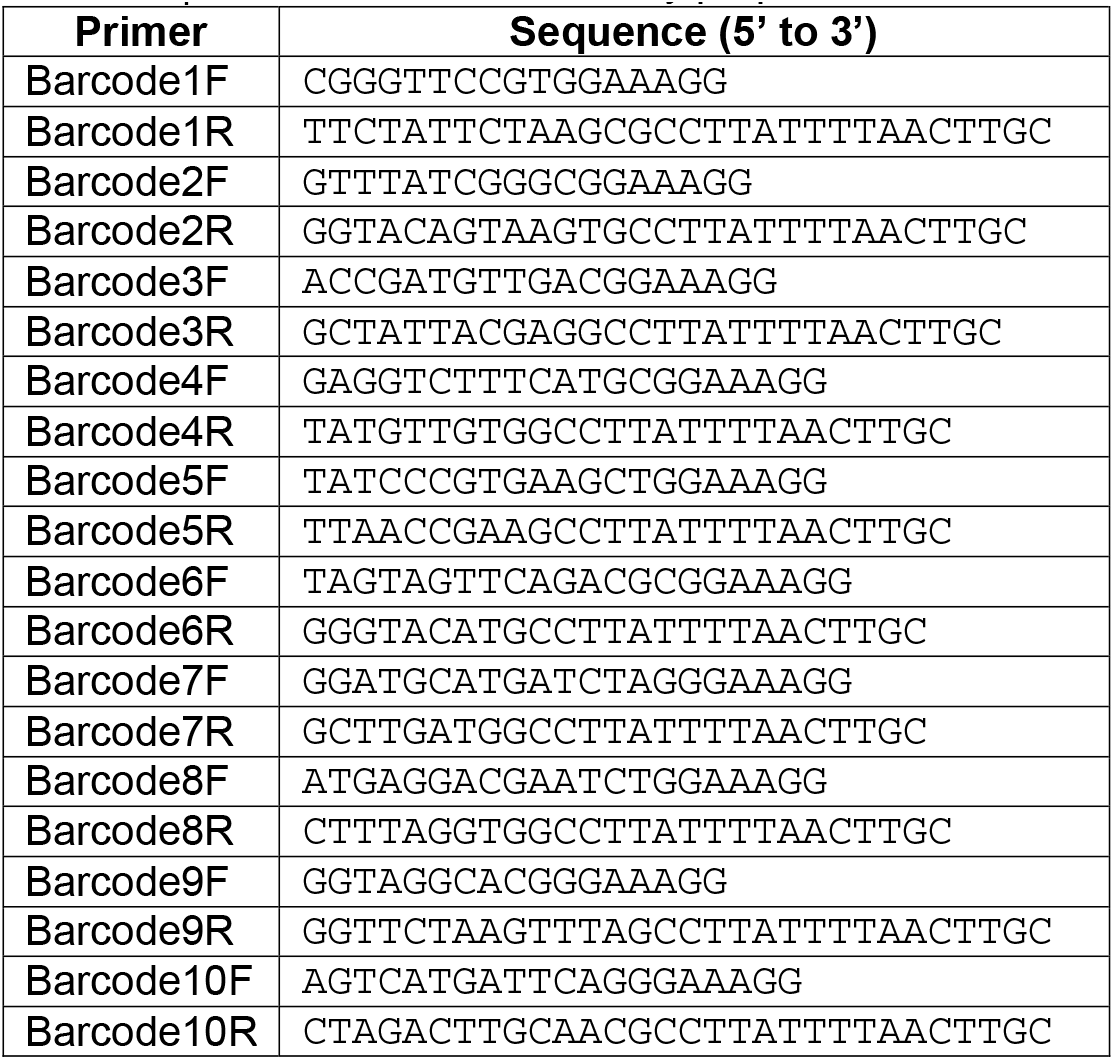
|Primers for lsPCR1 library preparation

14) Order synthesized DNA oligonucleotide pool synthesis on a programmable microarray (See Materials).

15) Obtain resuspended oligonucleotides for use as the template for the PCR reaction in Step 16. Consider freezing subset of oligonucleotides to save as a backup.

PAUSE POINT: Oligonucleotide pools can be stored at -20 ^o^C for short term storage (months to years) and -80^o^C for long term storage (years).

16) Set up barcode-specific PCR to amplify using barcode-specific primers (Table 5). This PCR will herein be referred to as “Library Synthesis PCR1” (lsPCR1).

CRITICAL STEP: It is important to use a proofreading polymerase to minimize introduction of PCR errors.

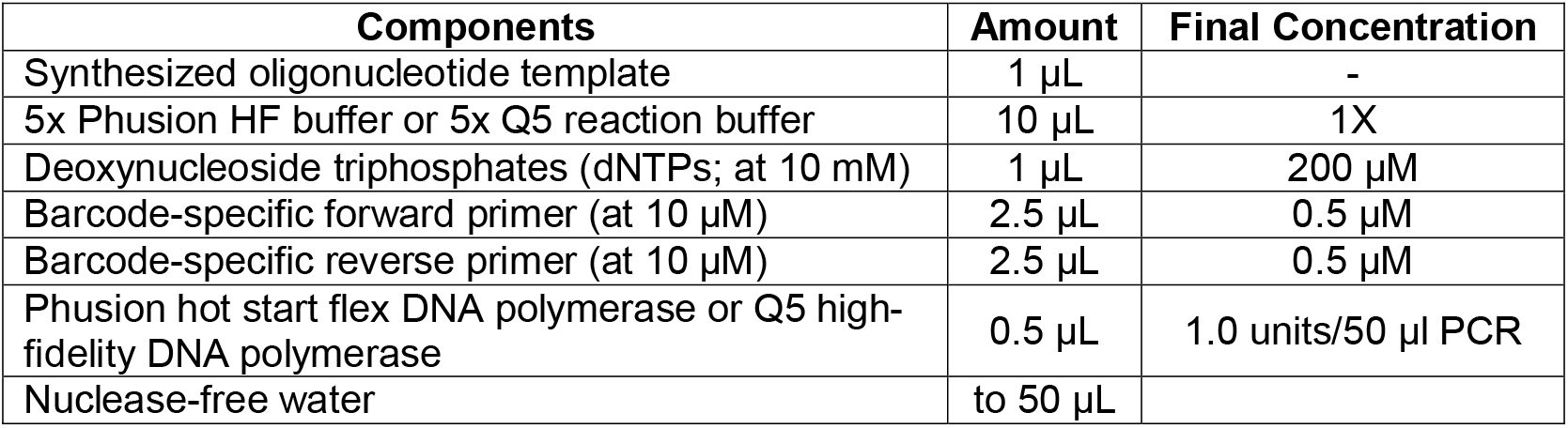

? TROUBLESHOOTING

17) Perform multiple cycles (e.g., 10, 15, 20 cycles) of lsPCR1 in a thermocycler as follows (Supplementary Fig. 3a):

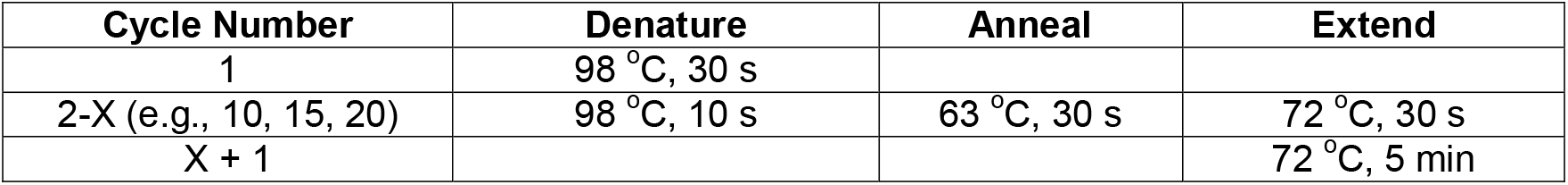

CRITICAL STEP: The number of PCR cycles should be limited to reduce PCR bias. It is difficult to predict the required number of cycles for a given library. Therefore, it is important to perform multiple different cycle numbers to empirically determine the optimal number of cycles. This consideration applies to lsPCR1 (Step 17) as well as lsPCR2 (Step 21).

PAUSE POINT: the PCR product can be stored at 4 ^o^C in the thermocycler at the end of the PCR program in the short term (days). Longer term storage should be at -20 ^o^C.

18) Run a fraction (1-5 µL of the 50 µL PCR reaction from Steps 16-17 is reasonable) of each lsPCR1 product with a different number of PCR cycles on a 2% (wt/vol) agarose gel. Analyze the gel to determine if the products occur at the expected size of ~100 bp (Supplementary Fig. 3a). If the bands run at the expected size, determine the minimal number of PCR cycles required to produce a visible band (Supplementary Fig. 3a). Utilize the remaining reaction volume (50 µL minus the volume run on the agarose gel) for the cycle number chosen based on the gel for Step 19.

CRITICAL STEP: The number of PCR cycles should be limited to reduce PCR bias. It is difficult to predict the required number of cycles for a given library. Therefore, it is important to perform multiple different cycle numbers to empirically determine the optimal number of cycles. The PCR reaction with the optimal cycle number as identified in Step 18 should be utilized for Step 19.

19) Dilute lsPCR1 reaction chosen from Step 17 1:10 using nuclease-free water.

20) Set up “Library Synthesis PCR2” (herein referred to as lsPCR2) with the following universal lsPCR2 primers (Table 6). lsPCR2 amplification removes the library-specific barcodes and adds sequence homologous to the lentiGuide-Puro plasmid for Gibson assembly-based cloning of the library.

**Table 6.**
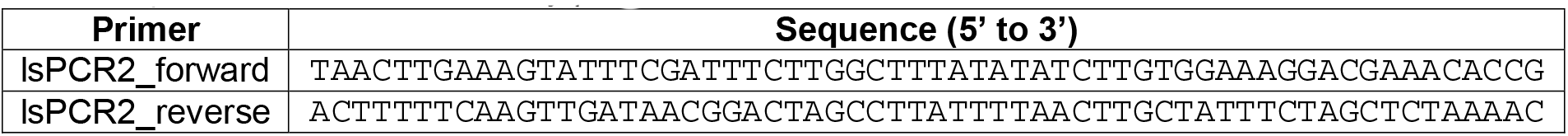
|Primers for lsPCR2 library preparation

CRITICAL STEP: It is important to use a proofreading polymerase to minimize introduction of PCR errors.

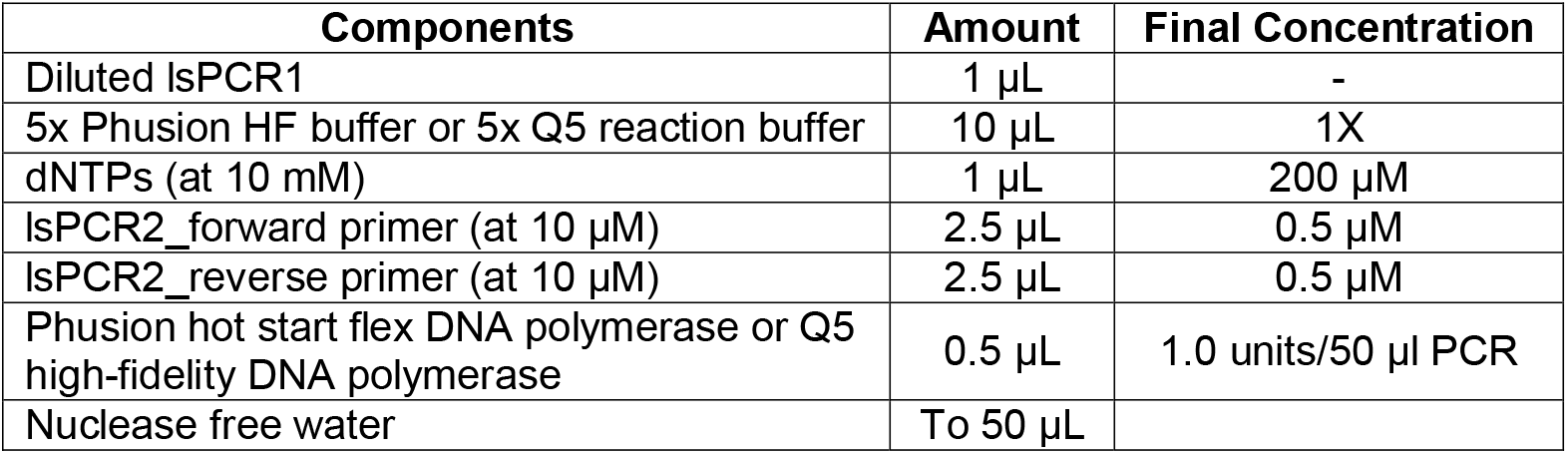

21) Perform multiple cycles (e.g., 10, 15, 20 cycles) of lsPCR2 in a thermocycler as follows (Supplementary Fig. 3b):

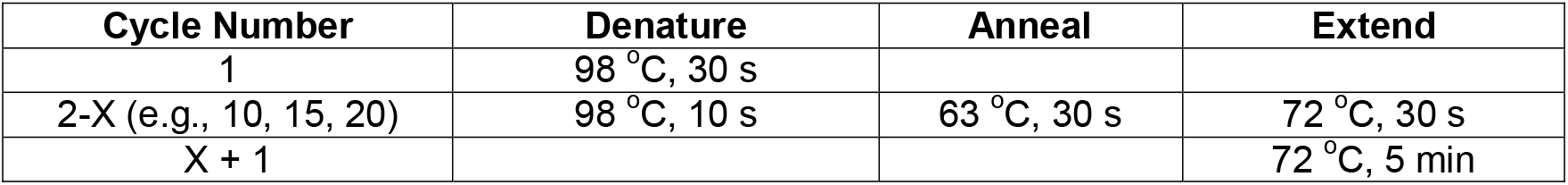

PAUSE POINT: PCR products can be stored at 4 ^o^C in the thermocycler at the end of the PCR program in the short term (days). Longer term storage should be at -20 ^o^C.

22) Run lsPCR2 products on a 2% (wt/vol) agarose gel to determine if the products occur at the expected size of ~140 bp (Supplementary Fig. 3b). Choose the sample with the minimal number of PCR cycles that produces a visible band in order to minimize PCR bias (Supplementary Fig. 3b). Gel purify the ~140 bp band.

CRITICAL STEP: If synthesizing multiple libraries simultaneously, leave empty lanes between different samples during gel electrophoresis to minimize risk of sample contamination during gel purification.

23) Set up Gibson assembly reaction using gel purified Esp3l-digested/dephosphorylated lentiGuide-Puro plasmid from Steps 5-7:

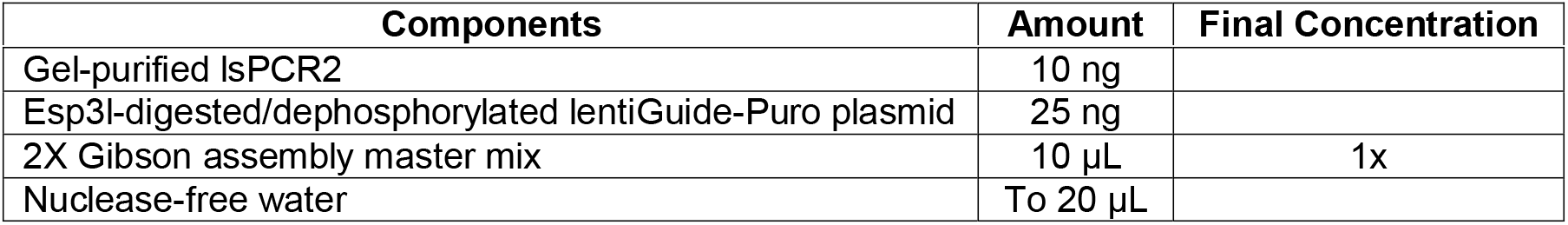

24) Incubate Gibson assembly reaction mix from Step 23 in a thermocycler at 50°C for 60 minutes

25) Thaw electrocompetent bacterial cells on ice until they thaw completely (~10-20 minutes).

26) Mix thawed electrocompetent bacterial cells by gently tapping on the side of the tube.

27) Aliquot 25 µL of cells to a pre-chilled microcentrifuge tube on ice.

28) Add 1 μL of the Gibson assembly reaction from Step 24 to the 25 µL of thawed electrocompetent bacterial cells from Step 27.

CRITICAL STEP: It is reasonable to aim for >50x representation of the sgRNA library. Depending on the achieved transformation efficiency as calculated in Step 37 and the number of sgRNA in the library, it may be necessary to set up >1 transformation reactions from Step 28. It is important to set up multiple reactions using 1 μL of Gibson assembly reaction and 25 μL of thawed electrocompetent bacterial cells. Increasing the amount of Gibson assembly reaction beyond 1 μL for a given electroporation can alter the chemistry of the electroporation and reduce transformation efficiency.

? TROUBLESHOOTING

29) Carefully pipet 25 µL of the electrocompetent bacterial cell/Gibson assembly reaction mixture from Step 28 into a chilled 0.1 cm gap electroporation cuvette without introducing bubbles. Quickly flick the cuvette downward to deposit the cells across the bottom of the cuvette’s well.

CRITICAL STEP: Minimize introduction of air bubbles as they can interfere with plasmid electroporation efficiency.

30) Electroporate sample at 25 μF, 200 Ohms, 1500 volts using an electroporator. Expected time constant during electroporation is ~4.7 (typical range of 4.2-4.8)

? TROUBLESHOOTING

31) Add 975 µL of recovery or SOC medium to the cuvette as soon as possible after electroporation (now final volume of 1 mL in cuvette). Pipet up and down enough times to resuspend the cells within the cuvette (likely a single pipetting up and down should be sufficient). Recovery medium is preferred; however, SOC medium can be used in its place.

CRITICAL STEP: Electroporation is toxic to bacterial cells. Addition of recovery or SOC medium as quickly as possible enhances bacterial cell survival post-electroporation.

32) Transfer the cell mixture (~1 mL) to a culture tube already containing 1 mL of SOC medium.

33) Shake each culture tube (containing a total of 2 mL of recovery/SOC medium) at 250 rpm for 1 hour at 37^o^C.

CRITICAL STEP: Growth at 32 ^o^C reduces recombination between the lentiviral long-terminal repeats; however, this is not essential and 37 ^o^C can be utilized.

CRITICAL STEP: If multiple transformations were set up in Step 28 using the same Gibson assembly reaction, the 2 mL culture mixes generated in Step 33 can be mixed together so that one set of dilution plates is sufficient for calculation of transformation efficiency (Step 34).

34) Remove 5 µL from the 2 mL mixture (or larger volume if samples were pooled/mixed) from Step 33 and add it to 1 mL of SOC medium. Mix well and plate 20 µL of the mixture (20,000x dilution) onto a pre-warmed 10-cm ampicillin-resistant (0.1 mg/mL ampicillin) agar plate and then plate the remaining 200 µL (2,000x dilution) onto a separate 10-cm ampicillin-resistant (0.1 mg/mL ampicillin) agar plate. These two dilution plates can be used to estimate transformation efficiency, which will help ensure full library representation of the sgRNA plasmid library. It is reasonable to aim for >50x representation of the sgRNA library.

CRITICAL STEP: Adjust the dilutions as necessary to obtain plates with a colony density that allows for accurate counting of the number of colonies on the 10-cm plates.

35) Plate the 2 mL transformation mixture (or 2-mL from a larger volume if samples were pooled/mixed) from Step 34 onto a pre-warmed 24.5 cm^2^ ampicillin-resistant (0.1 mg/mL ampicillin) agar plate using Pyrex beads to ensure even spreading. Spread the liquid culture until it is largely absorbed into the

agar and won’t drip when inverted for incubation in Step 36.

CRITICAL STEP: If multiple samples were mixed/pooled together in Step 34, it is important to still only plate 2-mL onto each pre-warmed 24.5 cm^2^ ampicillin-resistant agar plate as volumes >2-mL may not be fully absorbed into the agar.

36) Grow all three (or more) plates (2,000x dilution plate, 20,000x dilution plate, and 24cm^2^ non-dilution plate(s)) from Steps 34-35 inverted for 14-16 hours at 37^o^C.

CRITICAL STEP: Growth at 32 ^o^C reduces recombination between the lentiviral long-terminal repeats; however, this is not essential and 37 ^o^C can be utilized.

37) Count the number of colonies on the two dilution plates from Step 34. Multiple this number of colonies by the dilution factor (2,000x or 20,000x) and by the increased area of the 24.5 cm2 plate (~7.6-fold increase in area) for estimation of the total number of colonies on the 24.5 cm^2^ plate. It is reasonable to aim for >50x representation of the sgRNA library.

? TROUBLESHOOTING

38) Select ~10-20 colonies from the dilution plates (from Step 37) for screening by mini-scale plasmid preparation and subsequent Sanger sequencing (as in Steps 11-12) to determine if the sgRNA library has been PCR amplified. Given that the oligonucleotides for multiple libraries can be synthesized on the same programmable microarray (Steps 13-14), it is important to confirm that the correct library intended for amplification is obtained. This is also useful for preliminary evaluation of library representation. The expectation is to identify unique sgRNA sequences among all sequenced miniscale plasmid preparation given that probability of obtaining the same sgRNA from a full library of sgRNA should be low.

CRITICAL STEP: This step is intended to offer qualitative reassurance that library synthesis is proceeding in the expected manner by evaluating that the correct library has been PCR amplified and offering an initial assessment of library representation. Ultimately, the deep sequencing step (Step 42) provides a more comprehensive assessment of library composition.

? TROUBLESHOOTING

39) Pipette 10 mL of LB broth onto each 24.5 cm^2^ plate from Step 35 and scrape the colonies off with a cell scraper. The LB broth aides in removal of the colonies from the agar.

40) Pipette the LB broth/scraped bacterial colony mixture into a pre-weighed 50 mL tube and repeat the procedure a second time on the same plate with additional 5-10 mL of LB to maximize removal of bacterial colonies.

41) Centrifuge LB broth/scraped bacterial colony mixture at 400 x g for 5 minutes to pellet the bacteria and then discard the supernatant. Weigh the bacterial pellet in the tube (and subtract the pre-weighed tube to determine weight of the bacterial pellet) to determine the proper number of columns for maxi-scale plasmid preparation of the library. Each column can support ~0.45 g of bacterial pellet.

? TROUBLESHOOTING

42) Perform a sufficient number of maxi preps and combine eluted library plasmid DNA.

CRITICAL STEP: It is recommended to deep sequence the batch cloned plasmid library produced in Steps 13-42 to confirm successful library representation by following the deep sequencing procedure found in Steps 66-74 using library plasmid DNA as the template DNA for laPCR1 in Step 68.

### Lentivirus production from individual sgRNA plasmid or pooled sgRNA plasmid library | Timing 4 d

CAUTION: Take all necessary precautions for the handling lentivirus and disposal of lentiviral waste. Lentivirus is capable of integrating into the genome of human cells. This caution should be exercised during Steps 48-64.

43) Passage and maintain HEK293 cells with 16 mL of HEK293 medium as previously described^58^.

44) Perform transfection when HEK293 cells reach ~80% confluency in 15 cm round plate using polyethylenimine (PEI) as a transfection reagent. The plasmid:PEI ratio should be 1 μg of total transfected plasmid to 3 μg of PEI. Total plasmid consists of the sum of VSV-G μg + psPAX2 + library plasmid μg.

45) Mix PEI, VSV-G, psPAX2, and library plasmid in 1 mL of filtered DMEM without supplements in a sterile microcentrifuge tube:

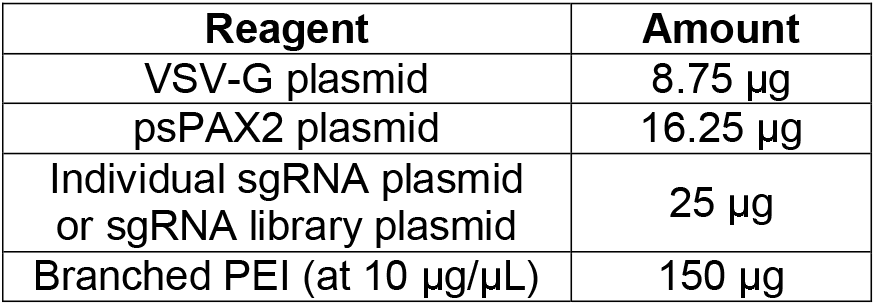

? TROUBLESHOOTING

46) Invert microcentrifuge tube several times to mix. Allow tube to incubate at room temperature (25 ^o^C) for 20-30 minutes.

CRITICAL STEP: the DMEM/plasmid DNA/PEI mixture should change from translucent to opaque during this incubation period.

47) Add the full volume (~1 mL) of DMEM/plasmid DNA/PEI mixture dropwise to the HEK293 cells and incubate at 37 ^o^C for 16-24 hours.

48) Replace the media with 16 mL of fresh HEK293 medium 16-24 hours after transfection in Step 47. The discarded media may contain lentivirus (albeit at low titer) so it should be disposed of as lentiviral waste.

49) Lentiviral supernatant harvest #1: 24 hours after replacing the medium in Step 47, collect the 16 mL media that now contains lentiviral particles (herein referred to as “viral supernatant”) in a 50 mL falcon tube. Replace with 16 mL of fresh HEK293 medium. Store the viral supernatant at 4^o^C.

CAUTION: Take all necessary precautions for the handling lentivirus and disposal of lentiviral waste. Lentivirus is capable of integrating into the genome of human cells.

50) Lentiviral supernatant harvest #2: 24 hours after lentiviral supernatant harvest #2, collect the viral supernatant and put it into the same 50 mL falcon tube from Step 49 (should now contain ~32 mL of viral supernatant).

51) Centrifuge collected viral supernatant from Step 50 at 500-700 x g for 5 minutes to pellet HEK293 cells and other debris.

52) Filter the viral supernatant through a 0.45 μm 50 mL filter.

? TROUBLESHOOTING

53) If sufficient viral titer is obtained with the viral supernatant from Step 52, the filtered viral supernatant can be stored at -80 ^o^C. It is recommended to freeze aliquots to minimize freeze-thaw cycles as they result in a reduction of viral titer. Aliquots also help to ensure consistent titer for each usage from the same batch of virus. If insufficient viral titer is achieved, it can be improved via ultracentrifugation (Steps 54-64). Viral titering can be performed through qPCR-based amplification of the HIV-1 genome (see Materials).

PAUSE POINT: The filtered viral supernatant can be stored at 4 ^o^C for up to 7 days prior to ultracentrifugation; however, storage at 4 ^o^C may result in reduction of viral titer. If ultracentrifugation is required (Steps 54-64), it is recommended to perform ultracentrifugation as soon as possible (minimize storage time at 4 ^o^C).

### Ultracentrifugation of viral supernatant (Optional) | Timing 2.5 hours

54) Transfer filtered viral supernatant to ultracentrifugation tubes.

55) Add 4-6 mL of 20% (wt/vol) sucrose solution to the bottom of the ultracentrifugation tube to create a sucrose layer (also known as a “sucrose cushion”) below the viral supernatant. This sucrose layer helps to remove any remaining debris from the viral supernatant as only the high-density viral

particles can pass through the sucrose layer to pellet while the low-density debris remain in the supernatant.

? TROUBLESHOOTING

56) Add sterile PBS to the top of the ultracentrifuge tube until the final liquid volume within the ultracentrifuge tube is ~2-3 mm from the top. The volume must be within 2-3 mm of the top of the tube to prevent the tube from collapsing due to the force of ultracentrifugation.

57) Place ultracentrifuge tubes into ultracentrifuge buckets.

58) Weigh all ultracentrifuge buckets containing ultracentrifuge tubes prior to ultracentrifugation to ensure appropriate weight-based balancing. Use sterile PBS to correct weight discrepancies.

59) Centrifuge at 100,000 x g for 2 hours in an ultracentrifuge

CAUTION: It is essential to weigh all samples prior to ultracentrifugation to ensure that the ultracentrifuge is appropriately balanced. Improper balancing can lead to spillage of lentiviral products, damage the ultracentrifuge, and/or harm to the user.

60) Remove ultracentrifuge tubes from ultracentrifuge buckets. The sucrose gradient should still be intact. ? TROUBLESHOOTING

61) Discard supernatant by inverting the ultracentrifugation tube. Allow inverted ultracentrifuge tube to stand on sterile paper towels to dry for 1-3 minutes.

? TROUBLESHOOTING

62) Revert ultracentrifugation tube to upright orientation and add 100 μL of sterile DMEM supplemented with 1% (vol/vol) FBS.

63) Incubate for 3-4 hours (or overnight) shaking at 4^o^C to resuspend the lentiviral pellet.

64) Store concentrated lentivirus in a microcentrifuge tube at -80^o^C. Due to loss of lentiviral titer with repeated freeze/thaw cycles, consider aliquoting at this point. Aliquots frozen at the same time can be assumed to have the same titer, which can be advantageous for multiple experiments using the same lentivirus.

### Execution of arrayed or pooled screen experiments | Timing 1-2 weeks

65) Experiments/screens using lentivirus should be performed as appropriate for the experimental design/objectives and the cell being utilized for study. Refer to Boxes 3 and 4 for discussion about and considerations for performing an arrayed or pooled screen experiment.

PAUSE POINT: After experiments/screens have been completed, cell pellets can be stored at -20 or - 80 ^o^C for weeks to months before processing for deep sequencing analysis of the samples (Steps 66-74).

### Deep sequencing of arrayed or pooled screen experiments | Timing 1 d

66) Extract genomic DNA (gDNA) from fresh cell pellets after completion of genome editing screens/experiments or from frozen pellets from previous experiments. See Materials for genomic DNA extraction kit and follow manufacturer instructions.

CRITICAL STEP: For arrayed experiments intended for indel analysis by CRISPResso (Step 76), it is recommended to include a non-edited sample from the same cell type. This sample will be a useful negative control when performing CRISPResso indel analysis/enumeration. Specifically, a non-edited sample is useful to help with optimization of CRISPResso analysis parameters summarized in Box 2. Furthermore, a non-edited sample can be useful in identifying single nucleotide polymorphisms (SNPs) or other variants present in the cells under study that can confound CRISPResso analysis. If a variant/indel is identified in the non-edited sample, CRISPResso settings can be altered to account for the variant/indel during CRISPR-mediated indel enumeration (Box 2).

67) Determine the concentration of each DNA sample using a UV-Vis spectrophotometer (e.g., NanoDrop) or other equivalent method.

68) Set up “Library Analysis PCR1” reaction (herein referred to as laPCR1). Use lentiGuide-Puro-specific primers (Table 7) to enumerate sgRNAs present for enrichment/dropout analysis or locus-specific primers to quantitate indels (Table 8).

**Table 7.**
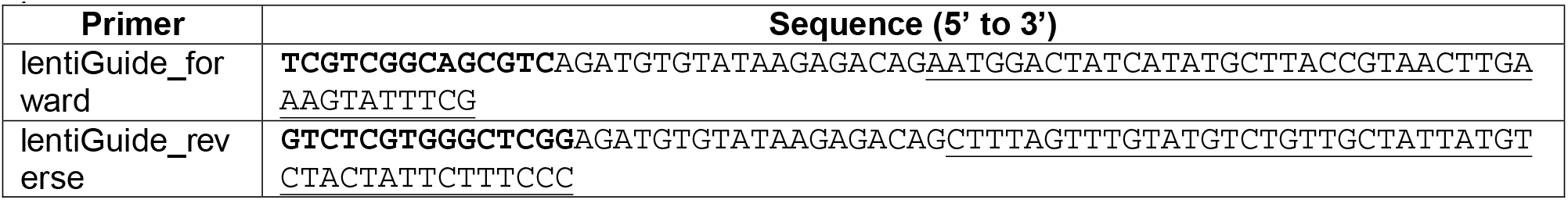
|Primers for laPCR1 for pLentiGuide-specific deep sequencing for sgRNA enumeration. Bold sequence is Illumina Nextera handle sequence. Underlined sequence is specific to the lentiGuide-Puro plasmid.

**Table 8.**
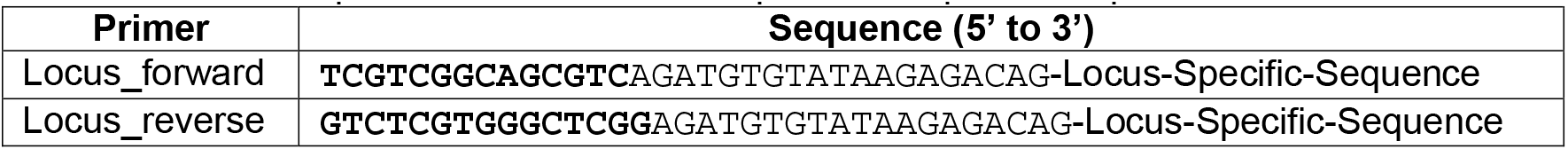
|Primers for laPCR1 for locus-specific deep sequencing. Bold sequence is Illumina Nextera handle sequence. Recommend 20 bp of locus-specific sequence.

CRITICAL STEP: It is important to use a proofreading polymerase to minimize introduction of PCR errors.

CRITICAL STEP: Optimization of DMSO concentration may be required (typically 1-10%, vol/vol). 8% (vol/vol) is used in the reaction below.

CRITICAL STEP: Amount of gDNA for laPCR1 can vary based on experimental needs for an arrayed experiment and based on the number of sgRNA in the library for a pooled experiment. On average, a genome from a single cell is approximately ~6.6 picograms^81^. Use adequate gDNA to represent the desired number of cells. For an arrayed experiment for indel analysis of a single locus using CRISPResso, the number of reads to adequately represent a locus may vary widely based on experimental goals; however, >1,000 cells at a given locus is reasonable. For a pooled sgRNA library experiment, 100-1,000x coverage of the sgRNAs in the library is reasonable. For example, 6.6 µg of gDNA is estimated to represent one million cells. For a library with 1,000 sgRNA, 6.6 µg of gDNA would provide 1000x coverage.

CRITICAL STEP: It is important to sequence the cloned plasmid library as generated in Steps 13-42. Sequencing of the plasmid library allows for confirmation of successful cloning of the library and can be used as an initial time point for a dropout (‘depletion’) screen (see Box 4 for details of this type of screen). Given that plasmids contain many fewer base pairs than a full genome, ≥50 picograms of plasmid DNA will be sufficient to represent a pooled plasmid library.

? TROUBLESHOOTING

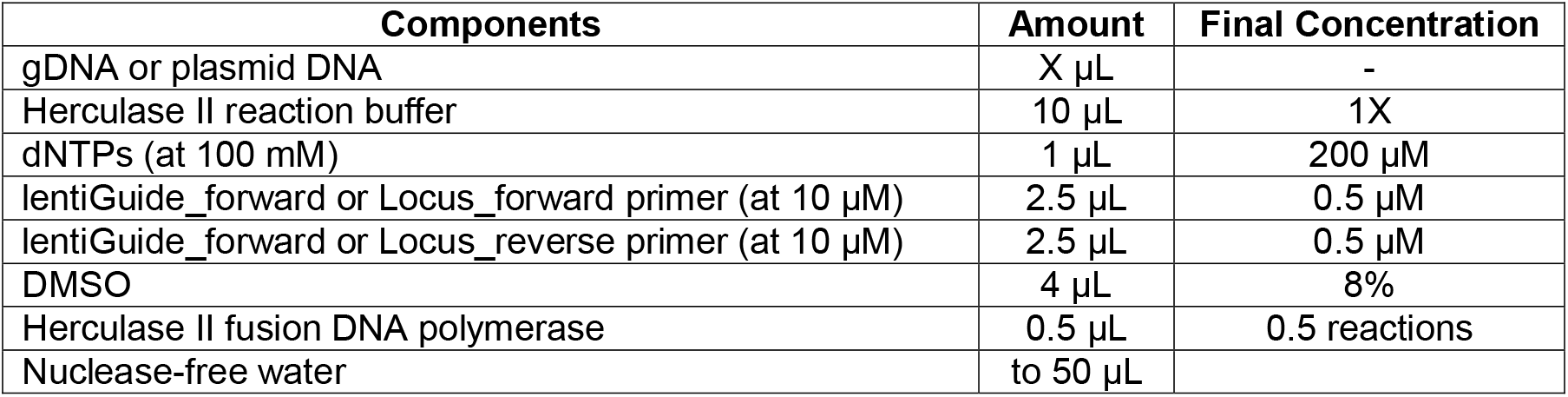

69) Perform multiple cycles (e.g., 10, 15, 20 cycles) of laPCR1 in a thermocycler as follows. Gradient PCR may be required to determine the optimal annealing temperature (Supplementary Fig. 3c).

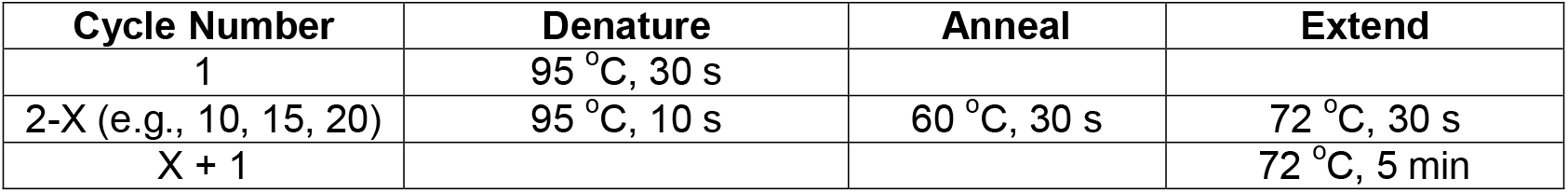

PAUSE POINT: the PCR product can be stored at 4 ^o^C in the thermocycler at the end of the PCR program in the short terms (days). Longer term storage should be at -20 ^o^C.

CRITICAL STEP: The number of PCR cycles should be limited to reduce PCR bias. It is difficult to predict the required number of cycles for a given library. Therefore, it is important to perform multiple different cycle numbers to empirically determine the optimal number of cycles.

? TROUBLESHOOTING

70) Set up “Library Analysis PCR2” (herein referred to as laPCR2) using barcode-specific primers (Tables 9-10). Each sample will have a unique Illumina Nextera index to allow demultiplexing of multiple samples sequenced together. Set up two PCRs for each sample.

**Table 9.**
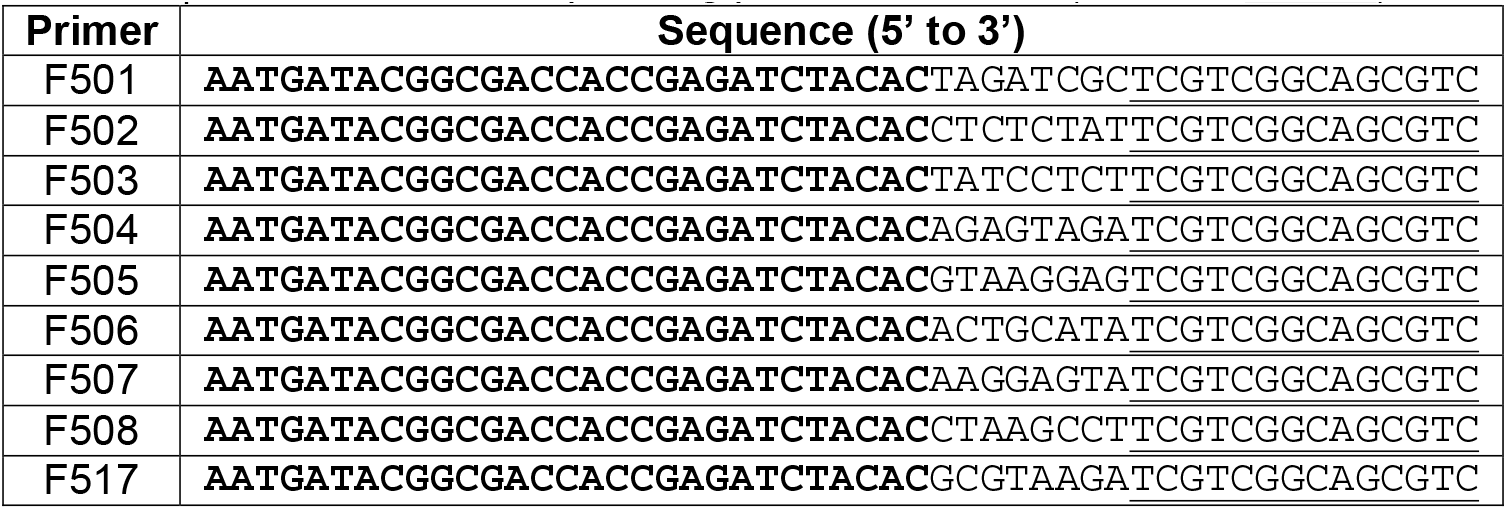
|Illumina forward sequencing primers for PCR2 (i5-Index-Handle)

**Table 10.**
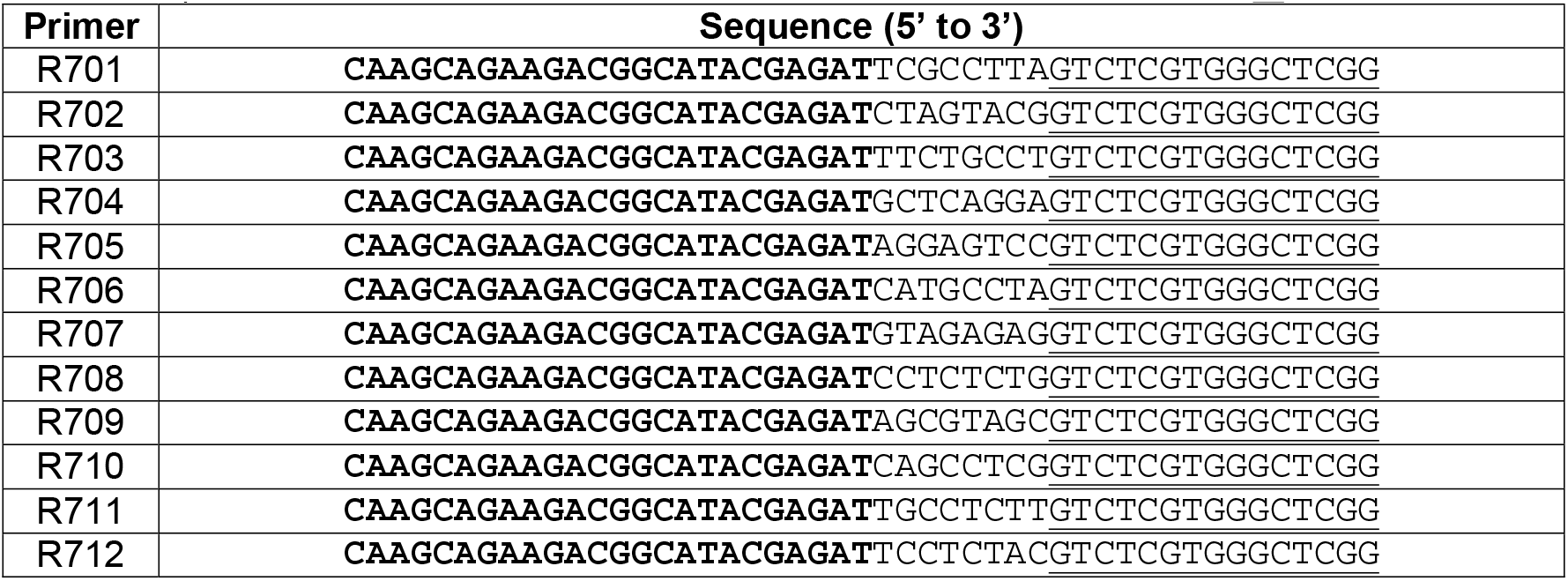
|Illumina reverse primers for deep sequencing (Reverse Primers (i7-Index-Handle)

CRITICAL STEP: Two separate 10 µL reactions are performed as opposed to a single 20 µL reaction to minimize PCR bias.

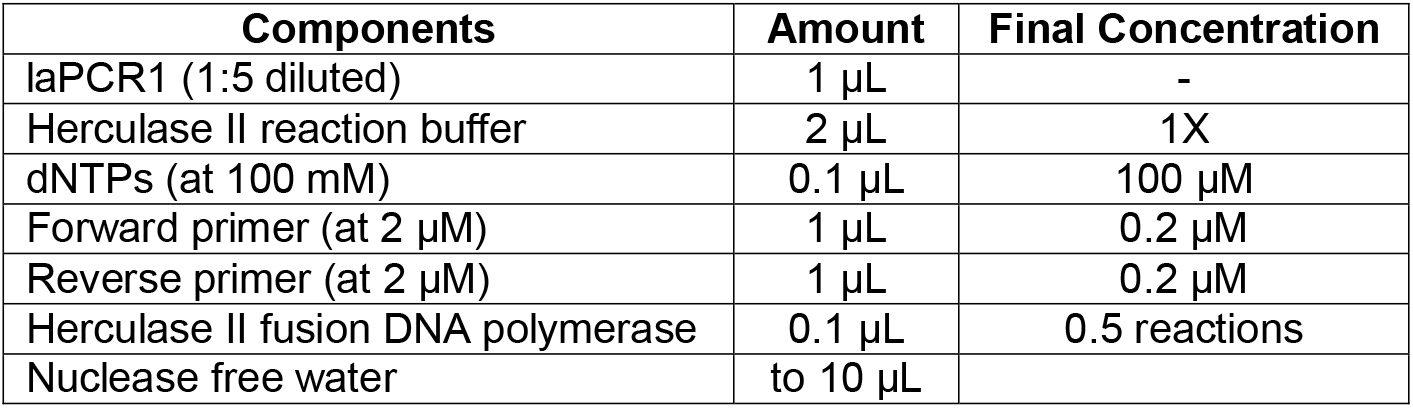

71) Perform multiple cycles (e.g., 10, 15, 20 cycles) of laPCR2 in a thermocycler as follows:

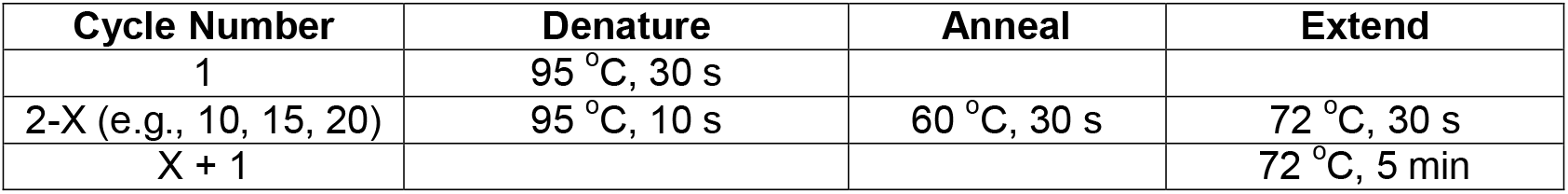

PAUSE POINT: the PCR product can be stored at 4 ^o^C in the thermocycler at the end of the PCR program in the short terms (days). Longer term storage should be at -20 ^o^C.

CRITICAL STEP: The number of PCR cycles should be limited to reduce PCR bias. It is difficult to predict the required number of cycles for a given library. Therefore, it is important to perform multiple different cycle numbers to empirically determine the optimal number of cycles.

72) Combine both 10 µL reaction for a total volume of 20 µL. Run laPCR2 products on a 2% (wt/vol) agarose gel to determine if the products occur at the expected size (expected size varies based on primers used in laPCR1). Choose the sample with the minimal number of PCR cycles that produces a visible band in order to minimize PCR bias (Supplementary Fig. 3d). Gel purify the relevant band of the expected size.

CRITICAL STEP: Leave empty lanes in between different samples to minimize risk of sample contamination during gel purification.

73) Quantitate DNA concentration by Qubit or other equivalent method.

74) The choice for appropriate deep sequencing platform (e.g., MiSeq or HiSeq) should include consideration for the required number of reads for the experiment as well as cost. For an arrayed experiment for indel enumeration/analysis by CRISPResso (Step 76), the number of reads to adequately represent a locus may vary widely based experimental goals; however, >1,000 reads at a given locus is reasonable. For a pooled sgRNA library experiment, 100-1,000x coverage of the number of sgRNA in the library is reasonable. The desired read length should be chosen based on the length of the amplicon generated by the primers used in Step 68 (refer to Step 1A(iv) for a discussion of amplicon length choice). Once the deep sequencing platform has been selected, submit barcoded samples for deep sequencing.

CRITICAL STEP: It can be useful to discuss experimental goals with the sequencing facility or individual performing the sequencing to help ensure the desired number of reads are obtained to adequately represent the sequenced locus or sgRNA library.

### CRISPResso installation and analysis of deep sequencing data | Timing 10-180 min

75) The CRISPResso utility can be used for analysis of deep sequencing data of a single locus/amplicon on a local machine or proprietary server, using a command line version or a webtool freely available at: http://crispresso.rocks (option A). The following steps will present detailed instructions for workflows using the webtool (option A), command line (option B), and Docker (option C). The webtool version of CRISPResso does not require installation. If using the webtool (option A), skip to the next step.

**(B) Installation of command line version of CRISPResso | Timing 15 min**

To install CRISPResso on a local machine, it is necessary first to install some dependencies before running the setup script:

i. Download and install Anaconda Python 2.7 following the instruction at this link: http://continuum.io/downloads.

ii. Open a terminal and type:

~~~
conda config–add channels r
conda config–add channels defaults
conda config–add channels conda-forge
conda config–add channels bio
conda install CRISPResso fastqc
~~~

iii. Close the terminal and open a new one, this will set the PATH variable. Now you are ready to use the command line version of CRISPResso.

iv. If you have installed an old version of CRISPResso, please remove it with the following command to avoid conflicts:

~~~
rm -Rf /home/user/CRISPResso_dependencies
~~~

? TROUBLESHOOTING

**(C) Installation of CRISPResso with Docker | Timing 15 min**

i. Check to have installed Docker (see Step 1C) and then type the command:

~~~
docker pull lucapinello/crispor_crispresso_nat_prot
~~~

ii. Check if the container was download successfully running the command:

~~~
docker run lucapinello/crispor_crispresso_nat_prot CRISPResso–help
~~~

If CRISPResso was properly installed, you will see the help of the command line version of CRISPResso.

76) Analysis of deep sequencing data with CRISPResso. The following step will present detailed instructions for workflows using the webtool (option A), command line (option B), and Docker (option C)):

**(A) Analysis of deep sequencing data using the CRISPResso webtool | Timing 15 min**

i) Open a web browser and point to the page: http://crispresso.rocks

ii) Select option for single-end (one FASTQ file) or paired end reads (two FASTQ files) based on the type of deep sequencing performed. Analysis of paired end reads requires overlapping sequence as the paired reads will be merged. Upload relevant FASTQ file(s) (as .fastq or .fastq.gz) for analysis. An example dataset to test the tool with previously generated FASTQ files is provided at http://crispresso.rocks/help (see the section: *Try it now*!).

? TROUBLESHOOTING

iii) Input amplicon sequence (required), sgRNA sequence (optional), and coding sequence(s) (optional input for when targeting exonic sequence). The reads uploaded in the previous step will be aligned to the provided amplicon sequence. If an sgRNA sequence is provided, its position will be indicated in all output analysis. If coding sequence is provided, it will allow for CRISPResso to perform frameshift analysis. The provided exonic sequence must be a subsequence of the amplicon sequence and not the sequence of the entire exon.

? TROUBLESHOOTING

iv) Adjust parameters for: window size (bp around each side of cleavage site) to quantify NHEJ edits (if sgRNA sequence provided), Minimum average read quality (Phred33 scale), Minimum single bp quality (Phred33 scale), Exclude bp from the left side of the amplicon sequence for the quantification of the mutations, Exclude bp from the right side of the amplicon sequence for the quantification of the mutations, and Trimming Adapter (see Box 2 for a discussion of these parameters).

v) Submit samples for analysis and download analysis reports. An illustrative video of the entire process is provided here: http://www.youtube.com/embed/dXblIIiAe00?autoplay=1

**(B) Analysis of deep sequencing data using command line CRISPResso | Timing 15 min**

i) Gather the required information and files (see Step 1A(iv) or 1C(xvi), the CRISPOR utility will output all the required information to run CRISPResso).

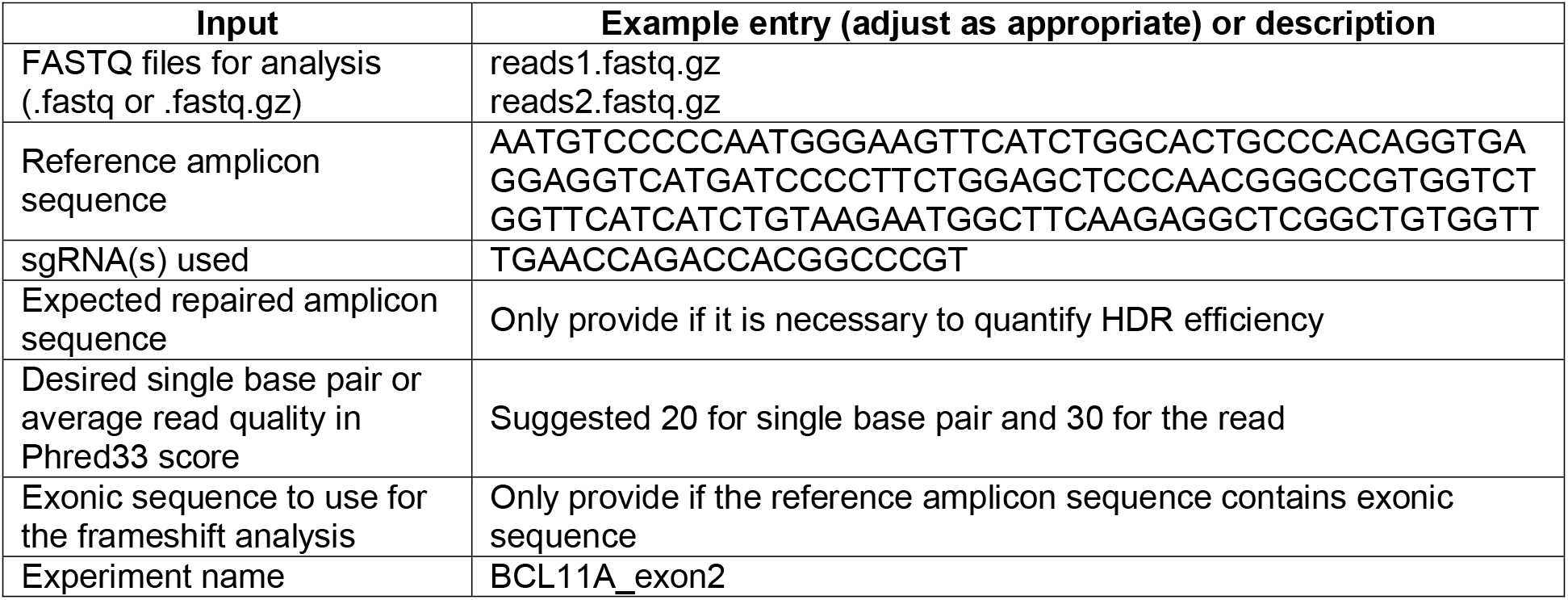

CRITICAL STEP: Be sure to check if the sequencing files were trimmed for adapters or not (refer to Box 2 for further discussion of the trimming process and options).

ii) Open a terminal and run the CRISPResso utility (the command below is based on the Step 76B(i)). Here we are assuming that the FASTQ files, reads1.fastq.gz and reads2.fastq.gz are stored in the folder: /home/user/amplicons_data/

~~~
CRISPResso -r1 /home/user/amplicons_data/reads1.fastq.gz -r2 /home/user/amplicons_data/reads2.fastq.gz -a AATGTCCCCCAATGGGAAGTTCATCTGGCACTGCCCACAGGTGAGGAGGTCATGATCCCCTTCTGGAGCTCCCAACGGGCCGTGGTCTGGTTCATCATCTGTAAGAATGGCTTCAAGAGGCTCGGCTGTGGTT -g TGAACCAGACCACGGCCCGT -s 20 -q 30 -n BCL11A_exon2
~~~

iii) After the execution of the command a new folder with all the results will be created, in this example the folder will be in:

`/home/user/amplicons_data/CRISPResso_on_BCL11A_exon2.` The summary of the different events discovered in the sequencing data is presented in the file `Alleles_frequency_table.txt`, and in several illustrative plots (.pdf files). Refer to the online documentation for the full description of the output:

https://github.com/lucapinello/CRISPResso

? TROUBLESHOOTING

**(C) Analysis of deep-sequencing data using CRISPResso with Docker | Timing 15 min**

i) Gather the required information and files as in option (B) (Step 76B(i)).

ii) Assuming that the FASTQ files, `reads1.fastq.gz` and `reads2.fastq.gz` are stored in the folder: `/home/user/amplicons_data/` run the command:

~~~
docker run -v /home/user/amplicons_data:/DATA -w /DATA
lucapinello/crispor_crispresso_nat_prot CRISPResso -r1
reads1.fastq.gz -r2 reads2.fastq.gz -a
AATGTCCCCCAATGGGAAGTTCATCTGGCACTGCCCACAGGTGAGGAGGTCATGATCCCCTT
CTGGAGCTCCCAACGGGCCGTGGTCTGGTTCATCATCTGTAAGAATGGCTTCAAGAGGCTCG
GCTGTGGTT -g TGAACCAGACCACGGCCCGT -s 20 -q 30 -n BCL11A_exon2
~~~

xviii) After the execution of the command a new folder with all the results will be created, in thi example the folder will be in:

`/home/user/amplicons_data/CRISPResso_on_BCL11A_exon2.` The summary of the different events discovered in the sequencing data is presented in the file `Alleles_frequency_table.txt`, and in several illustrative plots (.pdf files). Refer to the online documentation for the full description of the output:

https://github.com/lucapinello/CRISPResso?

TROUBLESHOOTING

### Analysis of a pooled sgRNA experiment using CRISPRessoCount | Timing 15 min

77) CRISPRessoCount is a utility for the enumeration of sgRNA (Fig. 2; Box 4). It is necessary to use the command line or Docker version of CRISPResso to run CRISPRessoCount (see Step 75B-C). CRISPRessoCount can enumerate sgRNAs from a user-generated list or can empirically identify all sgRNA present in the FASTQ file.

78) Gather the required data and information (adjust entries as appropriate for experiment):

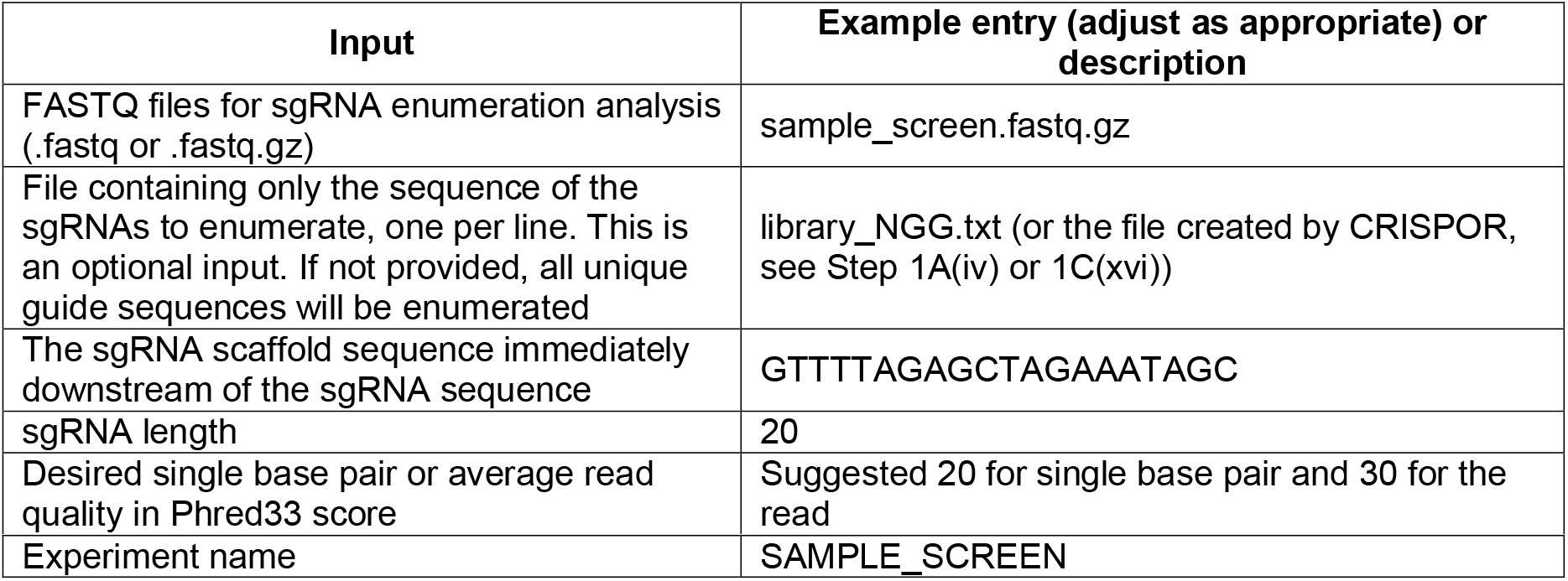

79) Run the CRISPRessoCount analysis. The following step will present detailed instructions for command line (option A) and Docker (option B). Notably, a webtool version of CRISPRessoCount is not available. Here we are assuming that the FASTQ file and the optional sgRNA file are stored in the folder: `/home/user/sgRNA _data/`

**(A) Using the command line version for CRISPRessoCount analysis (see Step 75B for installation) | Timing 15 min**

i. Execute the following commands

~~~
CRISPRessoCount -r /home/user/sgRNA_data/sample_screen.fastq.gz -s 20 -q 30 -f library_NGG.txt -t GTTTTAGAGCTAGAAATAGC -l 20–name SAMPLE_SCREEN
~~~

**(B) Using the Docker container for CRISPRessoCount analysis (see Step 75C for installation) | Timing 15 min**

i. Execute the following commands

~~~
docker run \
~~~

~~~
-v /home/user/sgRNA_data:/DATA \
~~~

~~~
-w /DATA \
~~~

~~~
lucapinello/crispor_crispresso_nat_prot \
~~~

~~~
CRISPRessoCount -r sample_screen.fastq.gz \
~~~

~~~
-s 20 -q 30 -f library_NGG.txt \
~~~

~~~
-t GTTTTAGAGCTAGAAATAGC -l 20–name SAMPLE_SCREEN
~~~

80) After the execution of the command a new folder with all the results will be created, in this example the folder will be in: `/home/user/amplicons_data/CRISPRessoCount_on_SAMPLE_SCREEN`. This folder contains two files, the execution log (`CRISPRessoCount_RUNNING_LOG.txt`) and a file (`CRISPRessoCount_only_ref_guides_on_sample_screen.fastq.gz.txt`) containing a table with the raw and normalized count for each sgRNA in the input FASTQ file:

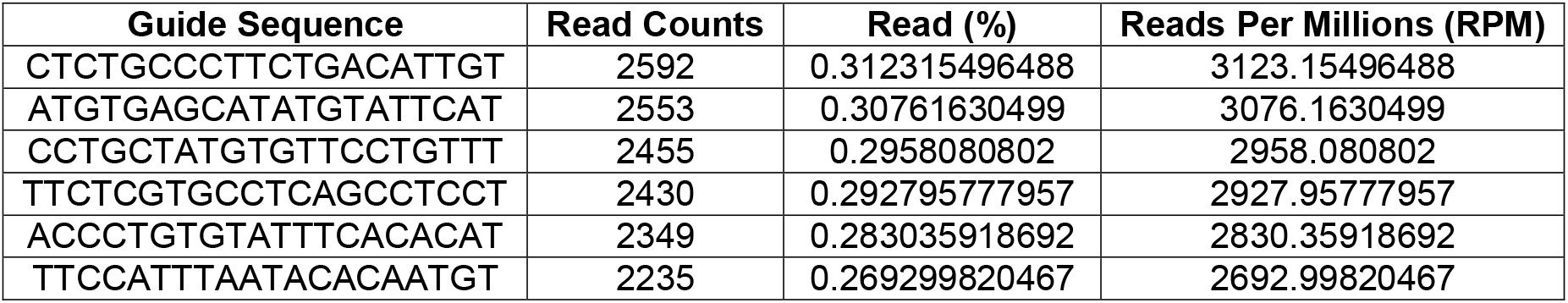

81) To calculate enrichment and/or dropout (‘depletion’) between two conditions, calculate the log2 ratio using the normalized sgRNA counts (that takes in account the total number of reads within each sequenced sample) using the column *Reads_Per_Millions_(RPM)* for the two conditions.

CRITICAL STEP: Read normalization is important to take into account the total number of reads when comparing two different deep sequencing samples. An example strategy to perform normalization involves normalizing all samples to one million reads. This can be accomplished by dividing the read count for each sgRNA by the total number of reads in that sample. Then this quotient is multiplied by one million. For example, (sgRNAn read count/total read count of sample)*1,000,000. This should be repeated for all samples so each sample has one million total reads. These normalized reads can then be used to calculate enrichment and/or dropout (‘depletion’) ratios. For example, log2(normalized read count of sgRNAn in sample X/normalized read count of sgRNAn in sample Y). The log2 is not necessary, but is often used when displaying the data since it will center the data around 0 for equal ratio, positive values will represent enrichment and negative values will represent depletion/dropout.

? TROUBLESHOOTING

### CRISPResso analysis using CRISPRessoPooled | Timing 20 min - 1 day

82) CRISPRessoPooled is a utility to analyze and quantify targeted sequencing CRISPR/Cas9 experiments involving sequencing libraries with pooled amplicons. To use CRISPRessoPooled is necessary to use the command line version of CRISPResso (see Step 75B). Although CRISPRessoPooled can be run in different modes (amplicon only, genome only, and mixed), in this protocol we use the most reliable mode for quantification called *mixed mode*. In this mode, the amplicon reads are aligned to both reference amplicons and genome to discard ambiguous or spurious reads. For more details regarding the different running modes, consult the online help: https://github.com/lucapinello/CRISPResso.

83) Gather the required data and information. FASTQ files to analyze containing reads for a pooled experiment with multiple sgRNA, for example:

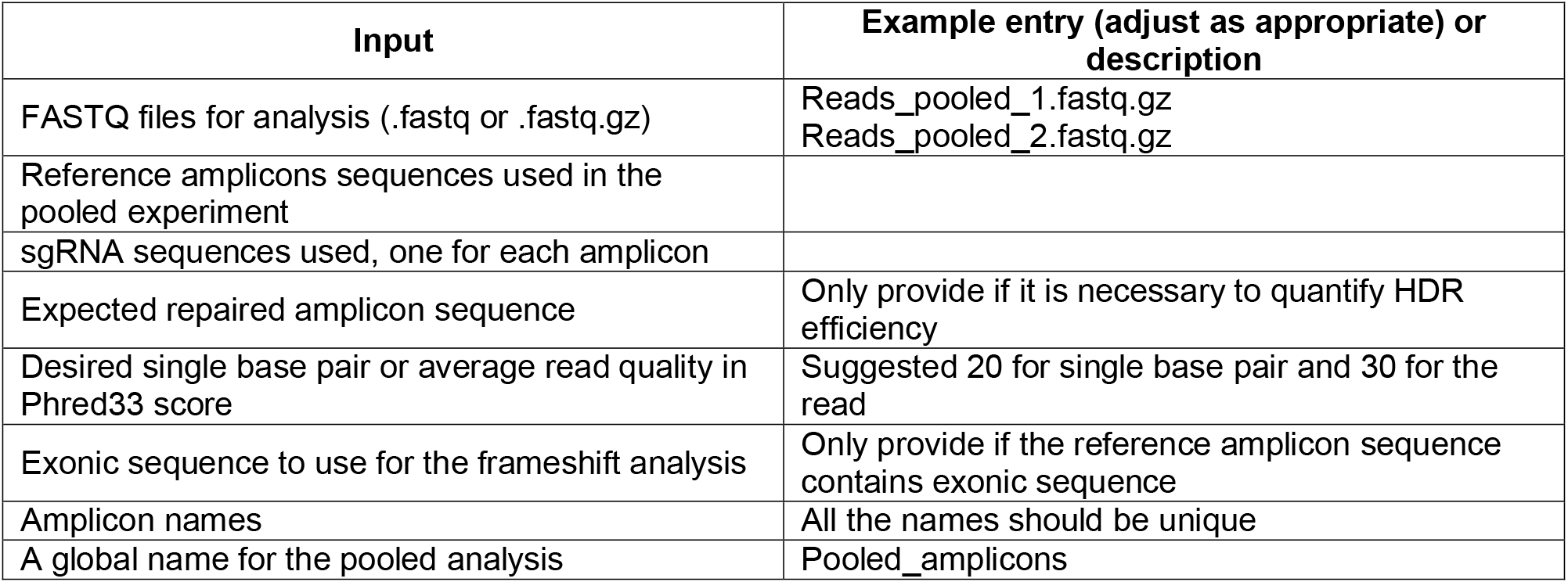

CRITICAL STEP: Be sure to check if the sequencing files were trimmed for adapters or not (refer to Box 2 for further discussion of the trimming options).

84) Create a folder for the reference genome and genomic annotations (this step is necessary only one time)

~~~
mkdir -p /home/user/crispresso_genomes/hg19
~~~

85) Download the reference genome: Download the desired genome sequence data and precomputed index from https://support.illumina.com/sequencing/sequencing_software/igenome.html. In this example, we download the hg19 assembly of the human genome.

86) Move to the folder from Step 84 that will be used to store the genome:

~~~
cd /home/user/crispresso_genomes/hg19
~~~

wget ftp://igenome:G3nom3s4u@ussd-

ftp.illumina.com/Homo_sapiens/UCSC/hg1938/Homo_sapiens_UCSC_hg19.tar.gz

87) Extract the data with the command:

~~~
tar -xvzf Homo_sapiens_UCSC_hg19.tar.gz \ Homo_sapiens/UCSC/hg19/Sequence/Bowtie2Index \–strip-components 5
~~~

88) Create an amplicon description file or use the amplicon file created by CRISPOR at Step 1A(iv) or 1C (xvi). This file, is a tab delimited text file with up to 5 columns (first 2 columns required):

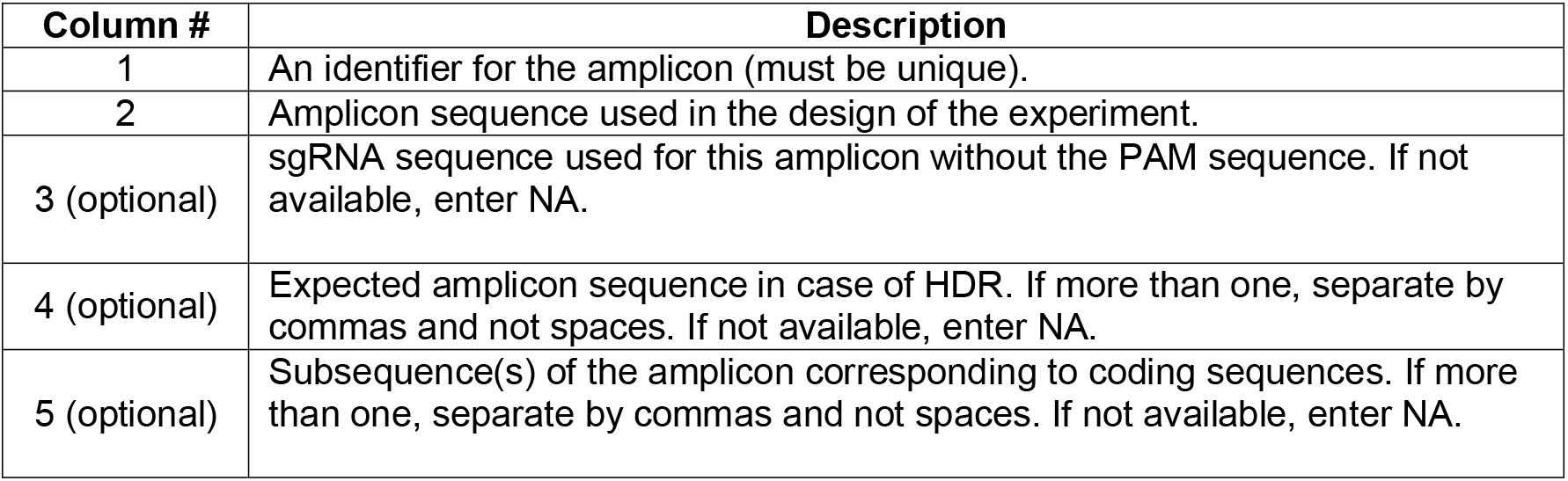

A properly formatted file should look like this:

~~~
Site1 CACACTGTGGCCCCTGTGCCCAGCCCTGGGCTCTCTGTACATGAAGCAAC CCCTGTGCCCAGCCC NA NA
~~~

~~~
Site2 GTCCTGGTTTTTGGTTTGGGAAATATAGTCATC NA GTCCTGGTTTTTGGTTTAAAAAAATATAGTCATC NA
~~~

~~~
Site3 TTTCTGGTTTTTGGTTTGGGAAATATAGTCATC NA NA GGAAATATA
~~~

No column heading is required.

Here we are assuming that the file is saved in the folder /home/user/pooled_amplicons_data/ and is called AMPLICONS_FILE.txt

89) Open a terminal and run the CRISPRessoPooled utility. The following step will present detailed instructions for command line (option A) and Docker (option B). Notably, a webtool version of CRISPRessoPooled is not available. Here we are assuming that the FASTQ files,

~~~
Reads_pooled_1.fastq.gz and Reads_pooled_1.fastq.gz are stored in the folder: /home/user/pooled_amplicons_data/
~~~

**(A) Using the command line version for CRISPRessoPooled analysis (see Step 75B for installation) | Timing 15 min**

i. Execute the following commands

~~~
CRISPRessoPooled \
~~~

~~~
-r1 /home/user/pooled_amplicons_data/Reads_pooled_1.fastq.gz \
~~~

~~~
-r2 /home/user/pooled_amplicons_data/Reads_pooled_2.fastq.gz \
~~~

~~~
-f /home/user/amplicons_data/AMPLICONS_FILE.txt \
~~~

~~~
-x /home/user/crispresso_genomes/hg19/genome \
~~~

~~~
-s 20 -q 30 \
~~~

~~~
–name Pooled_amplicons
~~~

**(B) Using the Docker container for CRISPRessoPooled analysis (see Step 75C for installation) | Timing 15 min**

i. Execute the following commands

~~~
docker run \
~~~

~~~
-v /home/user/amplicons_data/:/DATA -w /DATA \
~~~

~~~
-v /home/user/crispresso_genomes/:/GENOMES \ lucapinello/crispor_crispresso_nat_prot \
~~~

~~~
CRISPRessoPooled \
~~~

~~~
-r1 Reads_pooled_1.fastq.gz \
~~~

~~~
-r2 Reads_pooled_2.fastq.gz \
~~~

~~~
-f AMPLICONS_FILE.txt \
~~~

~~~
-x /GENOMES/hg19/genome \
~~~

~~~
-s 20 -q 30 \
~~~

~~~
–-name Pooled_amplicons
~~~

90) After the execution of the command in the previous step, a new folder with all the results will be created. In this example, the folder will be in:
`/home/user/amplicons_data/CRISPRessoPooled_on_Pooled_amplicons`. In this folder, the user can find these files:

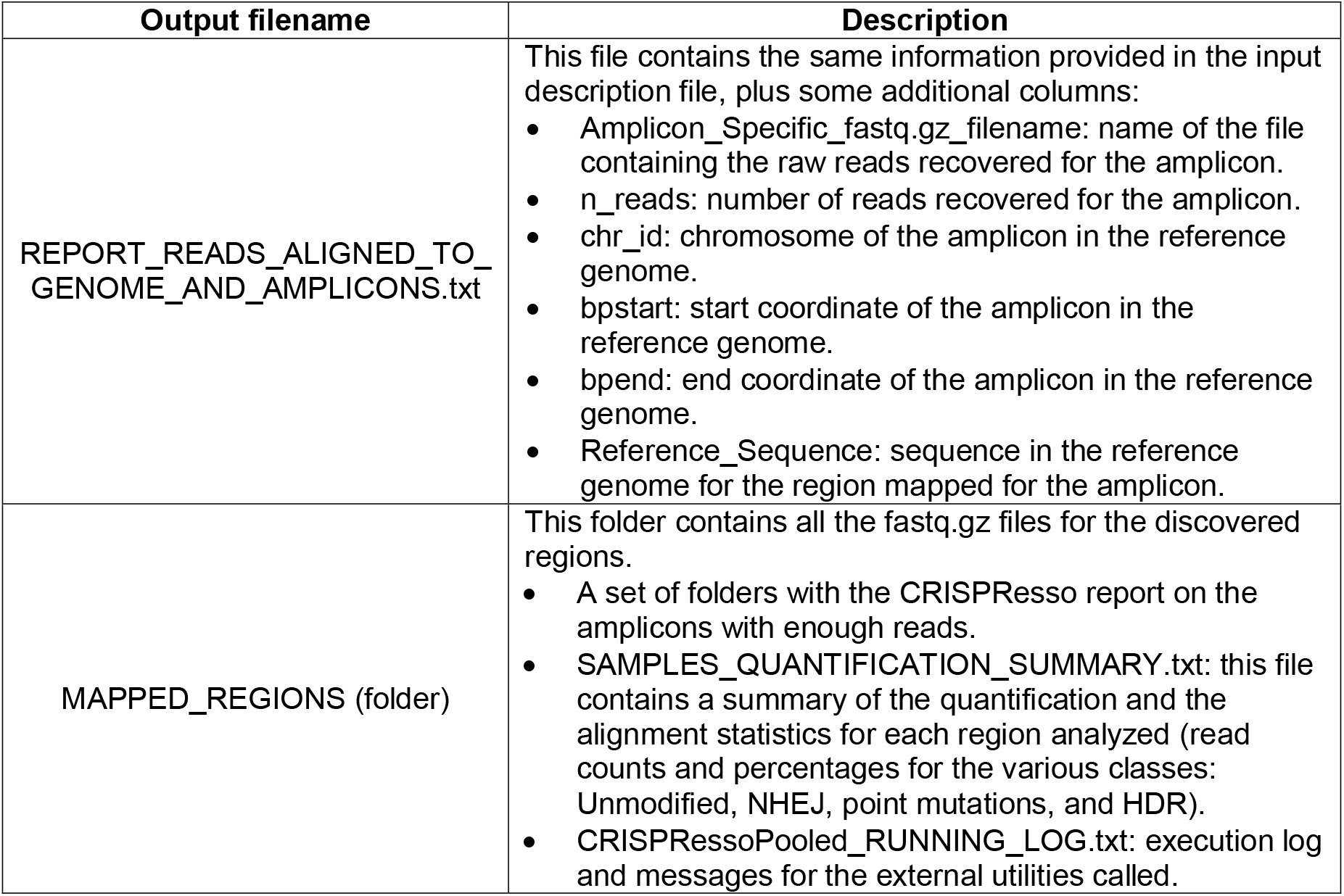

? TROUBLESHOOTING

### CRISPResso analysis using CRISPRessoWGS | Timing 1 h

91) CRISPRessoWGS is a utility for the analysis of genome editing experiment from whole genome sequencing (WGS) data. To use CRISPRessoWGS is necessary to use the command line version of CRISPResso (see Step 75B).

92) Gather the required data and information:

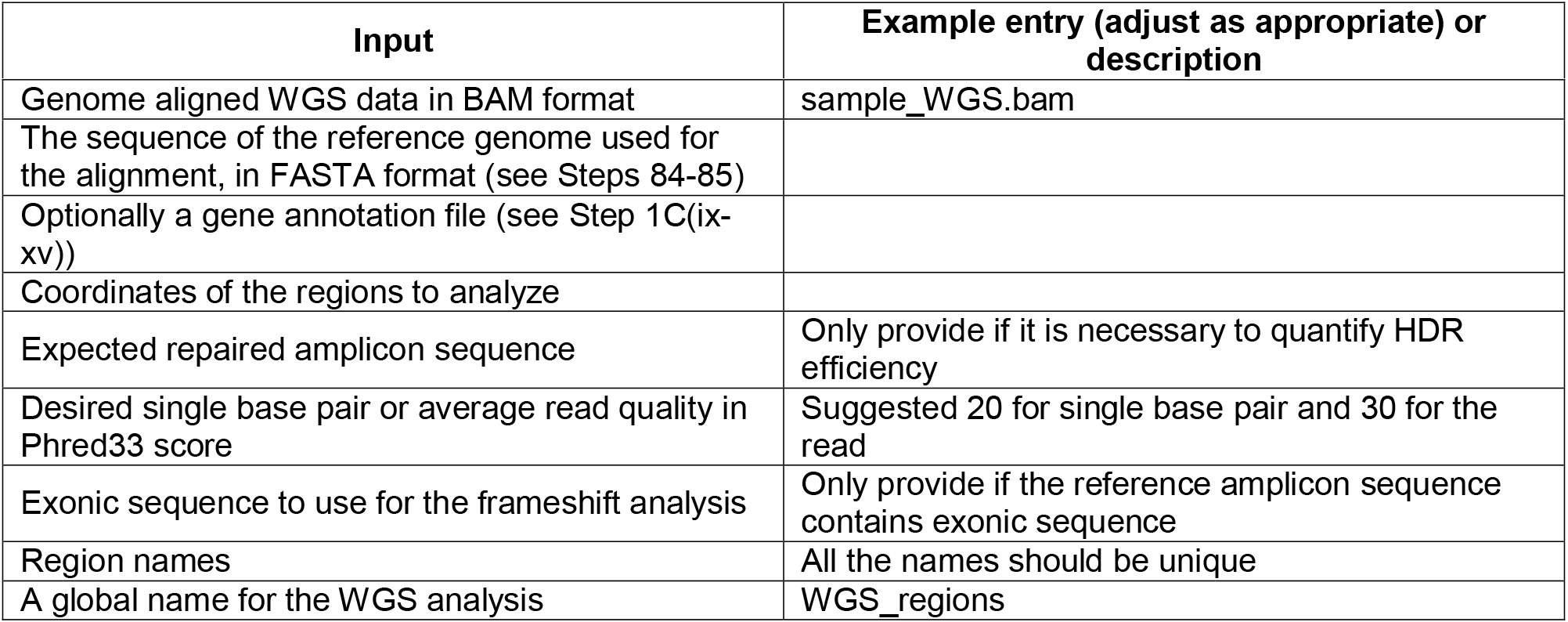

93) Create a regions description file. This file is a tab delimited text file with up to 7 columns (4 required) and contains the coordinates of the regions to analyze and some additional information:

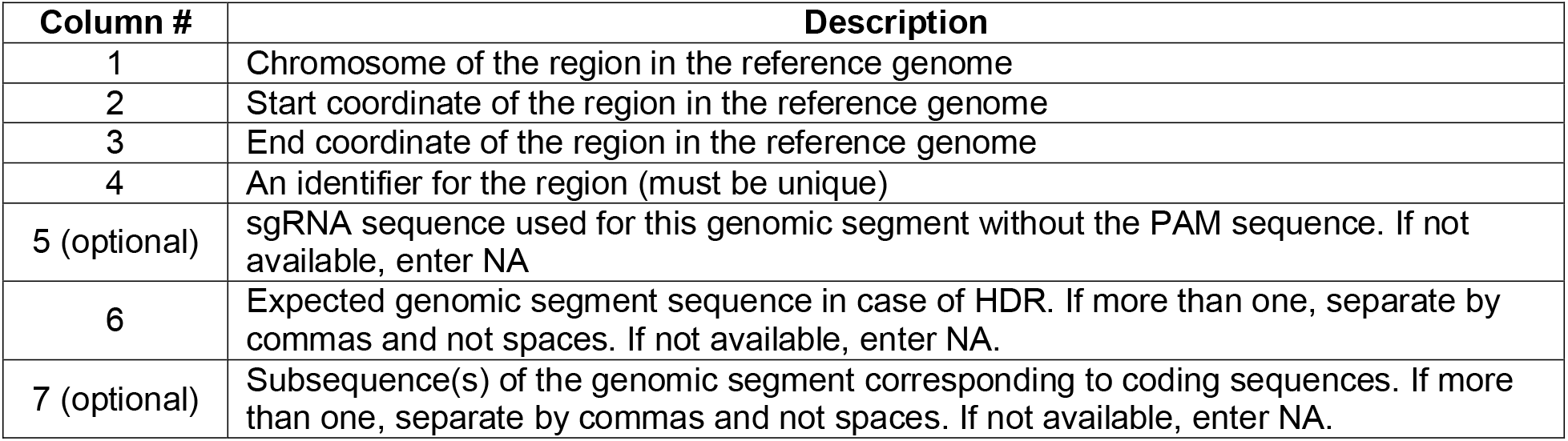

A properly formatted file should look like this:

~~~
chr1 65118211 65118261 R1 CTACAGAGCCCCAGTCCTGG NA NA
chr6 51002798 51002820 R2 NA NA NA
~~~

No columns heading is required.

Here we are assuming that the file is saved in the folder /home/user/wgs_data/ and it is called REGIONS_FILE.txt

94) Open a terminal and run the CRISPRessoWGS utility. The following step will present detailed instructions for command line (option A) and Docker (option B). Notably, a webtool version of CRISPRessoWGS is not available. Here we are assuming that the BAM is stored in the folder: /home/user/wgs_data/ and was aligned using the human hg19 reference genome.

**(A) Using the command line version for CRISPRessoWGS analysis | Timing 15 min**

i. Execute the following commands

~~~
CRISPRessoWGS\
~~~

~~~
-b /home/user/wgs_data/sample_WGS.bam\
~~~

~~~
-f /home/user/wgs_data/REGIONS_FILE.txt \
~~~

~~~
-r /home/user/crispresso_genomes/hg19/hg19.fa \
~~~

~~~
-s 20 -q 30 \
~~~

~~~
––name WGS_regions
~~~

**(B) Using the Docker container version for CRISPRessoWGS analysis | Timing 15 min**

i. Execute the following commands

~~~
docker run \
~~~

~~~
-v /home/user/wgs_data/:/DATA -w /DATA \
~~~

~~~
-v /home/user/crispresso_genomes/:/GENOMES \ lucapinello/crispor_crispresso_nat_prot \
~~~

~~~
CRISPRessoWGS \
~~~

~~~
-b sample_WGS.bam\
~~~

~~~
-f REGIONS_FILE.txt \
~~~

~~~
-r /GENOMES/hg19.fa \
~~~

~~~
-s 20 -q 30 \
~~~

~~~
–name WGS_regions
~~~

After the execution of the command a new folder with all the results will be created, in this example the folder will be in: /home/user/wgs_data/CRISPRessoWGS_on_WGS_regions. In this folder, the user can find these files:

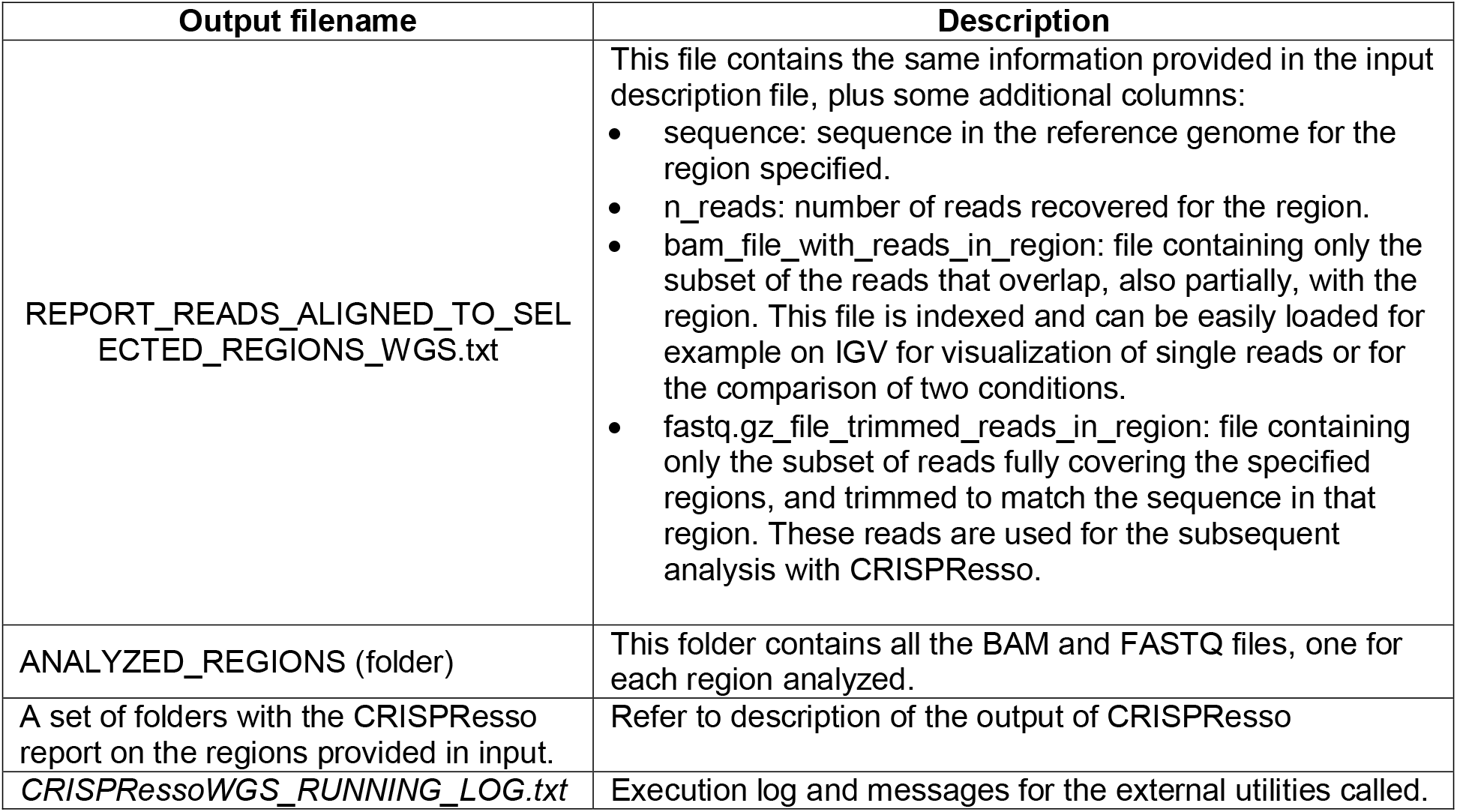

? TROUBLESHOOTING

### Direct comparison of CRISPResso analyses using CRISPRessoCompare and CRISPRessoPooledWGSCompare | Timing 30 min

95) CRISPRessoCompare and CRISPRessoPooledWGSCompare are two utilities allowing the user to directly compare two or several regions in two different conditions, as in the case of the CRISPResso, CRISPRessoPooled or CRISPRessoWGS. To use these utilities, it is necessary to use the command line version of CRISPResso (see Step 75B).

96) Gather the required data and information: Two completed CRISPResso, CRISPRessoPooled or CRISPRessoWGS analyses:

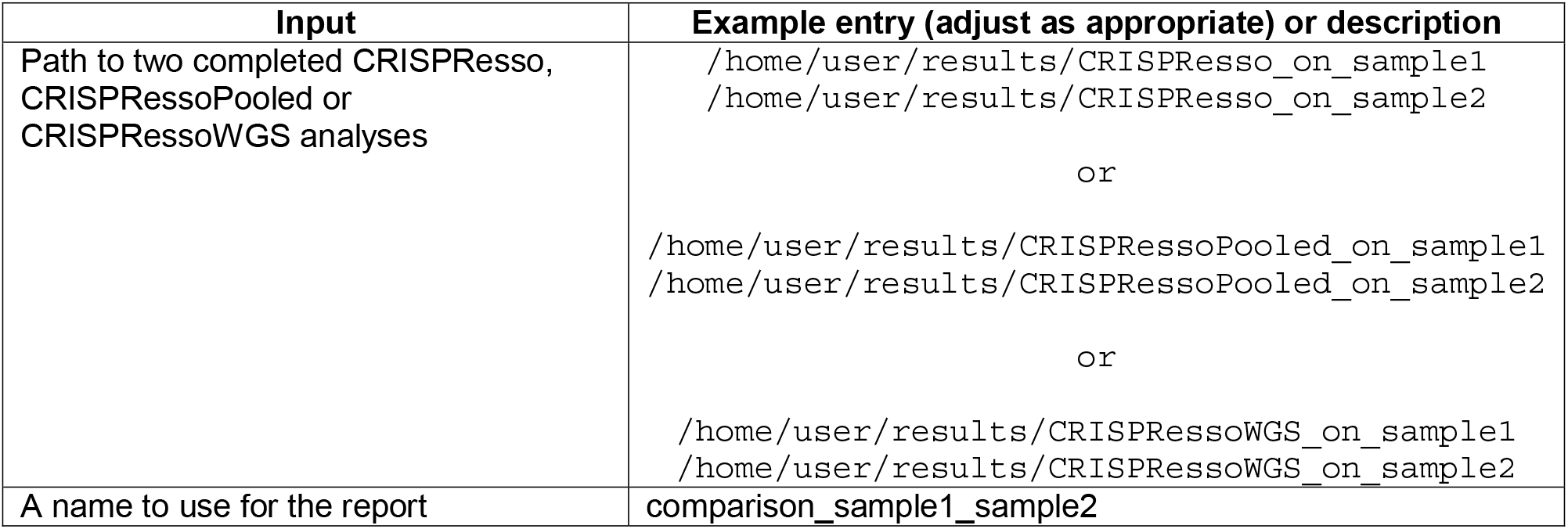

97) Open a terminal and run the CRISPRessoCompare or CRISPRessoPooledWGSCompare utility. The following step will present detailed instructions for command line (option A) and Docker (option B). Notably, a webtool version of CRISPRessoCompare or CRISPRessoPooledWGSCompare is not available. Here we are assuming that output folders from those utilities were saved in: `/home/user/results/`

**(A) Using the command line version (see Step 75B for installation) | Timing 15 min**

i. Execute the following command for CRISPRessoCompare

~~~
CRISPRessoCompare \
~~~

~~~
/home/user/results/CRISPResso_on_sample1 \ /home/user/results/CRISPResso_on_sample2 \
~~~

~~~
-n comparison_sample1_sample2
~~~

ii. Execute the following command for CRISPRessoPooledWGSCompare:

~~~
CRISPRessoPooledWGSCompare \
~~~

~~~
/home/user/results/CRISPRessoPooled_on_sample1 \
~~~

~~~
/home/user/results/CRISPRessoPooled_on_sample2 \
~~~

~~~
-n comparison_sample1_sample2
~~~

**(B) Using the Docker container (see Step 75C for installation) | Timing 15 min**

i. Execute the following command for CRISPRessoCompare:

~~~
docker run \
~~~

~~~
-v /home/user/results:/DATA -w /DATA \
~~~

~~~
lucapinello/crispor_crispresso_nat_prot \
~~~

~~~
CRISPRessoCompare \
~~~

~~~
CRISPResso_on_sample1 \
~~~

~~~
CRISPResso_on_sample2 \
~~~

~~~
-n comparison_sample1_sample2
~~~

ii. Execute the following command for CRISPRessoPooledWGSCompare:

~~~
docker run \
~~~

~~~
-v /home/user/results:/DATA -w /DATA \
~~~

~~~
lucapinello/crispor_crispresso_nat_prot \
~~~

~~~
CRISPRessoPooledWGSCompare \
~~~

~~~
CRISPRessoPooled_on_sample1 \
~~~

~~~
CRISPRessoPooled_on_sample2 \
~~~

~~~
-n comparison_sample1_sample2
~~~

The syntax for the utility CRISPRessoPooledWGSCompare is exactly the same for results obtained with CRISPRessoWGS.

After the execution of the command `CRISPRessoPooledWGSCompare`, a new folder with subfolders (one for each region) will be created. In this example, the folder will be in: `/home/user/results/CRISPRessoPooledWGSCompare_on_comparison_sample1_sample 2.`

In the folder created by CRISPRessoPooledWGSCompare, the user will see these files:

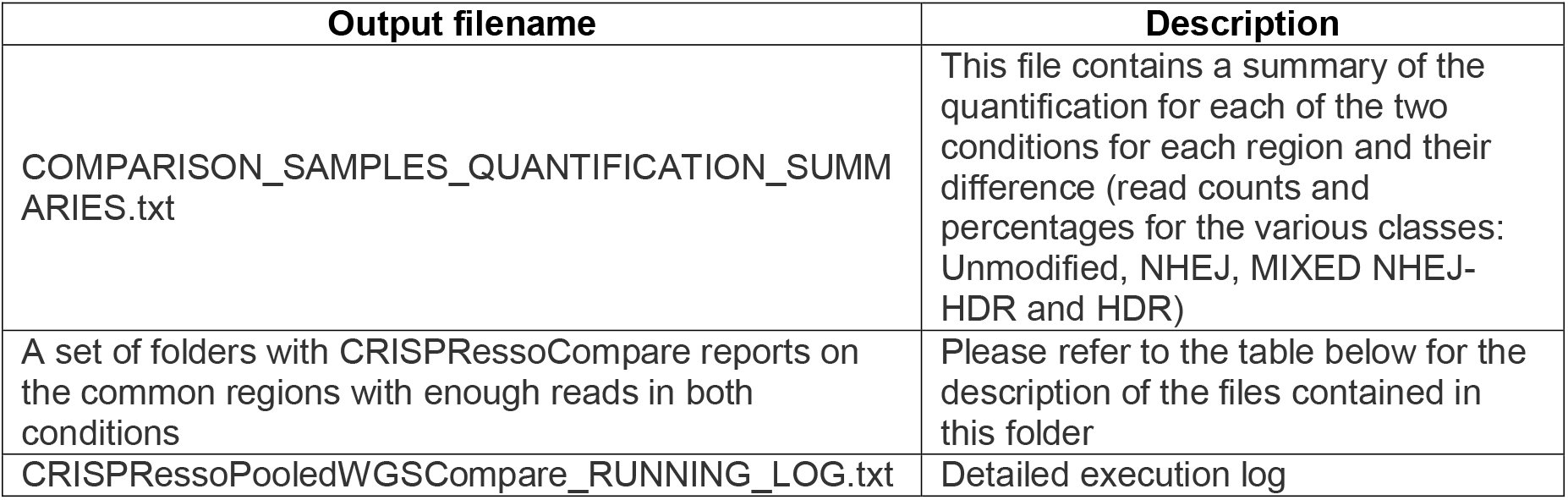

A single folder will be created if CRISPResso Compare was used instead. In the folder created by CRISPRessoCompare, the user will see these files:

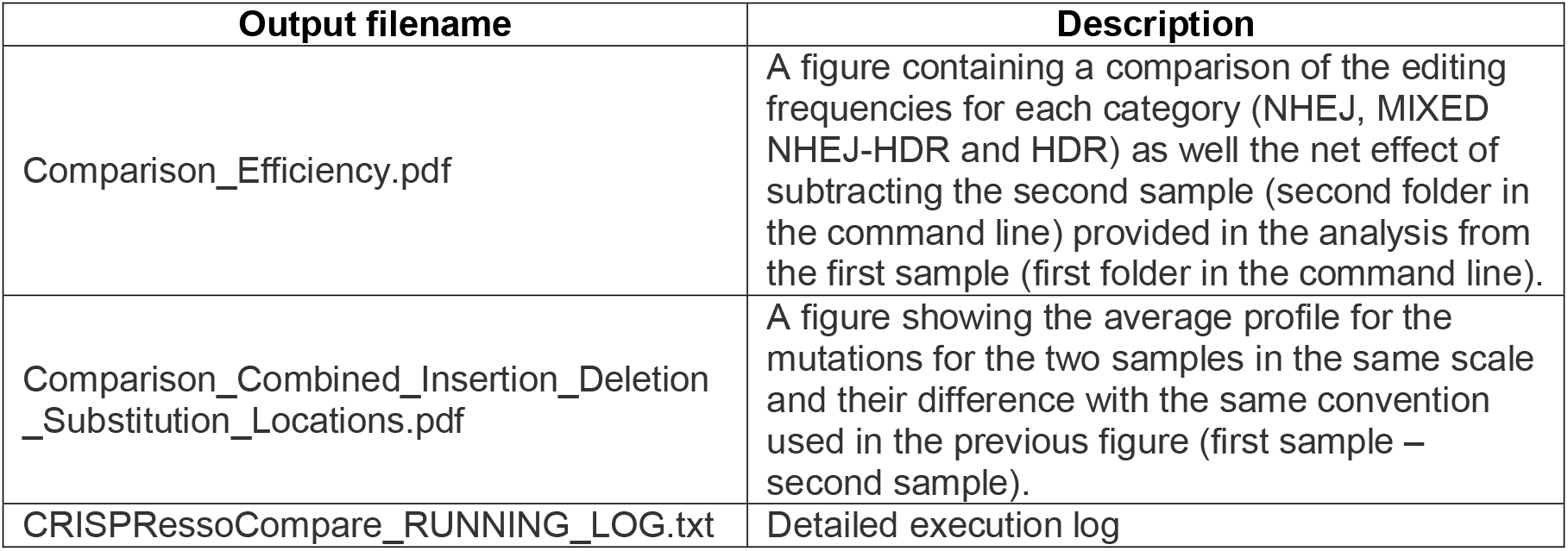

## TROUBLESHOOTING

**Table 11.**
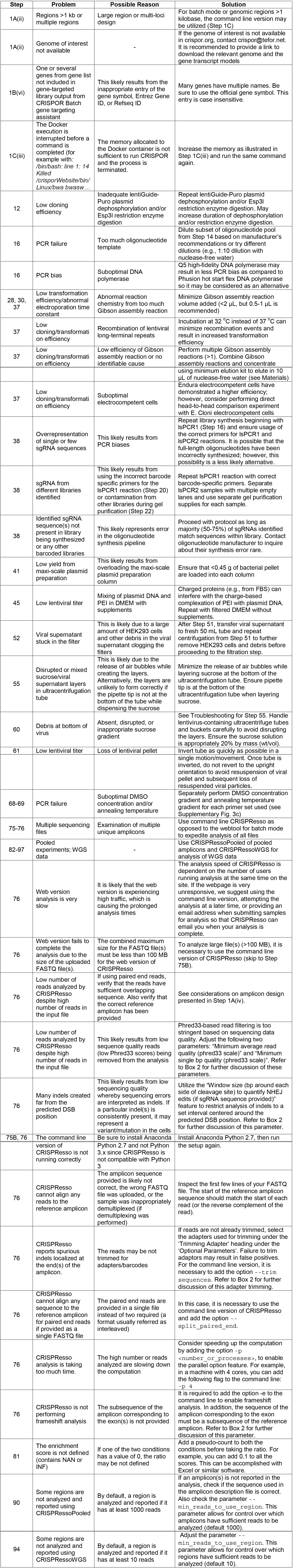
Troubleshooting

### Timing

Step 1, sgRNA design using CRISPOR: 1-4 h

Step 1A, sgRNA design for arrayed and pooled experiments using the CRISPOR webtool: 1-4 h

Step 1B, Gene-targeted library design generated from a gene list using the CRISPOR Batch gene targeting assistant webtool: 1-4 h

Step 1C, sgRNA design using command line CRISPOR: 1-4 h

Steps 1C(ix-xv), Generate FASTA file with exonic sequences (optional): 15 min

Step 1C(xiii), Run CRISPOR analysis on single- or multi-FASTA input file: 15 min

Step 1C(xvii), Selecting a subset of sgRNAs and primers for analysis: 15 min

Steps 2-12, Synthesis and cloning of individual sgRNA into a lentiviral vector: 3 d to obtain sequence-confirmed, cloned sgRNA lentiviral plasmid

Steps 13-42, Synthesis and cloning of pooled sgRNA libraries into a lentiviral vector: 2-4 d to obtain cloned sgRNA lentiviral plasmid library

Steps 43-64, Lentivirus production from individual sgRNA plasmid or pooled sgRNA plasmid library: 4 d

Step 65, Execution of arrayed or pooled screen experiments: 1-2 weeks

Steps 66-74, Deep sequencing of arrayed or pooled screen experiments: 1 d

Steps 75-76, CRISPResso installation and analysis of deep sequencing data: 10-180 min

Step 75B, Installation of command line version of CRISPResso: 15 min

Step 75C, Installation of CRISPResso with Docker: 15 min

Step 76A, Analysis of deep-sequencing data using the CRISPResso webtool: 15 min

Step 76B, Analysis of deep-sequencing data using command line CRISPResso: 15 min

Step 76C, Analysis of deep-sequencing data using CRISPResso with Docker: 15 min

Steps 77-81, Analysis of a pooled sgRNA experiment using CRISPRessoCount: 15 min

Step 79A, Using the command line version for CRISPRessoCount analysis (see Step 75B for installation): 15 min

Step 79B, Using the Docker container for CRISPRessoCount analysis (see Step 75C for installation): 15 min

Steps 82-90, CRISPResso analysis using CRISPRessoPooled: 20 min - 1 day

Step 89A, Using the command line version for CRISPRessoPooled analysis (see Step 75B for installation): 15 min

Step 89B, Using the Docker container for CRISPRessoPooled analysis (see Step 75C for installation): 15 min

Steps 91-94, CRISPResso analysis using CRISPRessoWGS: 1 h

Step 94A, Using the command line version for CRISPRessoWGS analysis: 15 min

Step 94B: Using the Docker container version for CRISPRessoWGS analysis: 15 min

Steps 95-97, Direct comparison of CRISPResso analyses using CRISPRessoCompare and CRISPRessoPooledWGSCompare: 30 min

## ANTICIPATED RESULTS

Indel enumeration/analysis of a deep sequencing amplicon generated by locus-specific PCR primers can be performed by CRISPResso for an arrayed experiment (Fig. 1, 3, 4a-d). Quantification of editing (Fig. 4a) and distribution of combined insertions/deletions/substitutions demonstrates successful editing of *BCL11A* exon 2 with indels flanking the double-strand break site (Fig. 4b-c). Frameshift analysis reveals a predominance of frameshift mutations as compared to in-frame mutations (Fig. 4d). The distribution of all identified alleles can be directly visualized (Supplementary Fig. 4). Direct comparison of the sample treated with CRISPR/Cas9 to a non-edited control using CRISPRessoCompared can offer reassurance that the identified mutations are likely to have resulted from CRISPR-mediated mutagenesis (Supplementary Fig. 5).

A pooled screen experiment can also be analyzed by CRISPRessoCount (Figs. 2-3). A saturating mutagenesis experiment was performed to identify functional sequence within the *BCL11A* enhancer. All sgRNA were designed within the core of the *BCL11A* enhancer^11,12^. All sgRNA were designed within *BCL11A* exon 2 to serve as positive controls and non-targeting controls were included as negative controls. sgRNAs were batched cloned into pLentiGuide-puro and lentivirus was produced. HUDEP-2 erythroid cells with stable Cas9 expression were transduced at low multiplicity. After puromycin selection and allowing 1-2 weeks for editing to occur, phenotypic selection occurred by fluorescence activated cell sorting (FACS). Specifically, cells with high-and low-fetal hemoglobin (HbF) were sorted by FACS. Genomic DNA was extracted from the high-and low-fetal hemoglobin sorted cell pellets, respectively. PCR was performed with primers specific to the pLentiGuide-Puro construct. After deep sequencing, FASTQ files were analyzed by CRISPRessoCount to determine read counts for each sgRNA in the library. Read counts were subsequently normalized for each sample. Normalized read counts from the high-HbF sample were divided by the normalized read counts from the low-HbF sample for each sgRNA followed by a log2 transformation to provide an enrichment score. The enrichment score for each sgRNA was plotted at the genomic position for the predicted double-strand break (Fig. 4e). Notably, this analysis shows pooled experiments using NGG-and NGA-restricted sgRNA (Table 1) and demonstrates reproducibility of the experimental results using this procedure across multiple studies. Based on the analysis presented in Figure 4e, enrichment of sgRNA based on HbF levels can be observed within the *BCL11A* enhancer comparable to targeting BCL11A coding sequence suggesting functional sequence within this region.

**Box 1** | Selection of sgRNA based on on-target and off-target prediction scores.

CRISPOR provides two separate types of predictions: on-target and off-target scores. Off-target scores try to estimate the off-target cleavage potential, which depends on the sequence similarity between the on-target sequence and predicted off-target sequences. Predictions are first calculated on the level of the individual putative off-target site using the cutting frequency determination (CFD) off-target score. An off-target with a high off-target score is more likely to be cleaved than one with a low score. The scores of all off-target sites of a guide are then summarized into the “guide specificity score”. It ranges from 0 to 100; a guide with a high specificity score is expected to have fewer off-targets and/or off-targets with a lower cleavage frequencies than one with a lower score. It was previously shown that guides with specificity scores >50 never lead to more than around a dozen off-targets and none had a total off-target cleavage >10% in whole-genome assays^9^. These guides are shown in green by CRISPOR.

On the level of the individual site, off-targets with a CFD score <0.02 are very unlikely to be cleaved based on previous analysis that demonstrated only a few true off-target events with such a score (Haeussler et al^9^, Fig. 2). For off-target sites that are cleaved, the CFD score is well-correlated with editing frequency but predicted off-target sites are known to be primarily false positives. As such, for most guides, more than 95% of predicted off-targets will not show any cleavage detectable with the sequencing depths in use today. In addition, as summarized in Haeussler et al.^9^, a few guides with low specificity scores (in the range 10-30) lead to fewer than 3-5 detected off-targets, so low specificity scores do not always translate to strong off-target effects.

The other type of prediction is on-target scoring, which tries to estimate on-target cleavage efficiency and depends only on the target sequence. CRISPOR offers eight algorithms in total with two scores identified as most predictive when compared on independent datasets^9^. The algorithm by Doench et al^21^ (herein referred to as the “Doench 2016 score”) is optimal for cell-culture based assays where the guide is expressed from a U6 promoter like as described in this protocol. The score by Moreno-Mateos et al^48^ (CRISPRscan) is best for guides expressed *in vitro* with a T7 promoter (e.g. for mouse/rat/zebrafish oocyte injections, not discussed further here). On-target scoring is not very reliable as the Pearson correlation between cleavage and the Doench 2016 score is only around 0.4. Despite this, there is an enrichment for top-scoring guides. For example, for guides in the top 20% of Doench 2016 scores, 40% of them are in the top quartile for efficiency, which is a 1.5-fold enrichment of top quartile guides compared to random guide selection.

By default, guides in CRISPOR are sorted by guide specificity score and off-targets are sorted by CFD score. The relative importance of off-target specificity versus on-target efficiency depends on the experiment. In some cases, such as when targeting a narrow region, one has little choice and a guide with low specificity (i.e., many/high-probability off-targets) is the only one available. In this case, more time and effort may be needed to screen for off-targets. To provide an estimate of the number of off-targets to test, previous work showed that most off-targets of guides with a specificity of > 50 are detected by screening the 96 predicted off-targets with the highest CFD scores^9^.

Both off-and on-target scores have been pre-calculated for most common model organisms and are shown when hovering with the computer mouse over a guide in the UCSC Genome Browser track “Genes” - “CRISPR” at http://genome.ucsc.edu. The sequence currently shown in the UCSC Genome Browser can be sent directly to CRISPOR by selecting “View– In external tools” in the Genome Browser menu. In the UCSC track, guides are colored by predicted on-target efficiency whereas they are colored by off-target specificity on Crispor.org.

It can be particularly important to use off-target scoring prediction to aid in saturating mutagenesis experiments for sgRNA depletion or dropout^12^. sgRNA with highly probable off-target sites may deplete/dropout from screens due to cellular toxicity resulting from multiple cleavages as opposed to a biological effect from mutagenesis of the on-target site.

**Box 2** | Optimizing parameters for CRISPResso analysis.

CRISPResso allows many parameters to be set/adjusted based on the desired analysis to be performed. Details on all the parameters are available in the online help section (http://crispresso.rocks/help). Here we discuss the key parameters that can significantly affect the quantification.

**Amplicon sequence** (-a): This is the sequence expected to be observed without any edits. The sequence must be reported without adapters and barcodes.

**sgRNA sequence** (*-g*): This is the sequence of the sgRNA used and it should be a subsequence of the amplicon sequence (or its reverse complement). Although this parameter is optional, it is required to specify it to enable the window mode (see the parameter–window_size in order to reduce false positives in the quantification). It is important to remember that the sgRNA needs to be input as the guide RNA sequence (usually 20 nt) immediately upstream of the PAM sequence for Cas9 species (Table 1). For other nucleases, such as Cpf1, enter the sequence (usually 20 nt) immediately downstream of the PAM sequence and explicitly set the cleavage offset (see the parameter:–cleavage_offset).

**Coding sequence** (*-c*): The subsequence/s of the amplicon sequence covering one or more coding sequences. Without this sequence, frameshift analysis cannot be performed.

**Window size** (-w): This parameter allows for the specification of a window(s) in bp around each sgRNA to quantify indels. This can help limit sequencing/amplification errors and/or non-editing polymorphisms (e.g., SNPs) from being inappropriately quantified by CRISPResso’s analysis. The window is centered on the predicted cleavage site specified by each sgRNA. Any indels that do not overlap and/or substitutions that are not adjacent to the window are excluded from analysis. A value of 0 will disable this filter (default: 1).

**Cleavage offset** (––cleavage offset): This parameter allows for the specification of the cleavage offset to use with respect to the provided sgRNA sequence. The default is -3 and is suitable for *S. pyogenes* Cas9. For alternate nucleases (Table 1), other cleavage offsets may be appropriate. For example, set this parameter to 1 if using Cpf1.

**Average read and single bp quality** (–min_average_read_quality and–min_single_bp_quality): These parameters allow for the specification of the minimum average quality score or the minimum single bp score for inclusion of a read in subsequent analysis. The scale used is Phred33 (default: 0, minimum: 0, maximum: 40). The PHRED score represents the confidence in the assignment of a particular nucleotide in a read. The maximum score of 40 corresponds to an error rate of 0.01%. This average quality of a read is useful to filter out low quality reads. A reasonable value for this parameter is 30. More stringent filtering can be performed by using the single bp quality; any read with a single bp quality below the threshold will be discarded. A reasonable value for this parameter is greater than 20.

**Identity score** (–min_identity_score): This parameter allows for the specification of the minimum identity score for the alignment (default: 60.0). In order for a read to be considered properly aligned, it should pass this threshold. We suggest lowering this threshold only if really large insertions or deletions are expected in the experiment (>40% of the amplicon length).

**Exclude ends of the read** (–exclude_bp_from_left and–exclude_bp_from_right): Artifacts are sometimes present at the ends of the reads due to imperfect trimming or due to a drop in quality scores. To exclude these regions at the ends of reads, these parameters allow for the exclusion of a few bp from the left and/or right of the amplicon sequence during indel quantification (default: 15).

**Trimming of adapters** (–trim_sequences): This parameter enables the trimming of Illumina adapters with Trimmomatic (default: False). For custom adapters, the user can customize the Trimmomatic execution using the parameter–trimmomatic_options_string (check the Trimmomatic manual for more information on the different flags and options: http://www.usadellab.org/cms/uploads/supplementary/Trimmomatic/TrimmomaticManual_V0.32.pdf). It is important to check with your sequencing facility to determine if the reads were trimmed for adapters or not. If this is not possible, we suggest using the software FASTQC (http://www.bioinformatics.babraham.ac.uk/projects/fastqc/). In order to perform this analysis, open a terminal and type: fastqc. This command will open the main software window. After opening the FASTQ file(s) to analyze with FASTQC, a graphical report will be automatically generated. In the graphical report, find the section “Adapter content”. If the reads are not properly trimmed, the adapter used will be reported. If a custom adapter was used, find the section “Overrepresented sequences” instead where all the sequences that are present in more than 0.1% reads are reported. A subsequence present in more than 10% of the reads is usually an indication that the reads were not properly trimmed. An example of FASTQC report is available here: http://www.bioinformatics.babraham.ac.uk/projects/fastqc/RNA-Seq_fastqc.html#M9

**Box 3** | Technical and experimental considerations for performing an arrayed CRISPR genome editing experiment.

Here are some practical guidelines for executing an arrayed CRISPR experiment. An experimental schematic can be found in Figure 1 and an example workflow in Figure 3:

**Generation of cells with stable CRISPR nuclease expression**: For cell lines, it is convenient to generate lines with stable CRISPR nuclease expression for arrayed experiments. This can be accomplished by transducing cells with a lentiviral CRISPR nuclease with subsequent selection for transduced cells, such as the usage of lentiviral Cas9 with blasticidin resistance (the focus of the remainder of the discussion in Box 3 will focus on the usage of *S. pyogenes* Cas9; however, the same principles apply to other CRISPR nucleases including Cpf1). It is recommended that a “kill curve” is created to determine the optimal concentration of blasticidin for the cells used prior to beginning the experiment. It is also important to determine the duration of selection required to complete cell death by blasticidin. Stable expression of Cas9 can be confirmed via Western blot. Alternatively, stable expression can be confirmed by assessing Cas9 function using a reporter system, such as the previously described constructs that provide GFP and an sgRNA targeting GFP to assess for functional Cas9 through flow cytometry (see Materials)^12,46^. It is possible to screen for clones with high Cas9 expression and/or high Cas9 activity as assessed by a reporter; however, it is not required. If stable Cas9 expression cannot be generated, such as usage of primary cells with limited culture duration, co-transduction of Cas9 and sgRNAs can be performed with double selection (blasticidin for Cas9 and puromycin for sgRNA). Co-transduction can occur simultaneously or can occur on back-to-back days. Selection by blasticidin and/or puromycin is typically added 24-48 hours after transduction.

**Arrayed sgRNA experiment execution**: Unlike a pooled screen experiment as described in Box 4, copy number (i.e., number of viral integrants per cell) may not be an important consideration for many experiments since only one sgRNA is being used. Therefore, it may only be important for certain applications for cells to have one copy of the sgRNA or have a mixture of ≥1 copy. Therefore, lentiviral transduction of Cas9-expressing cells can occur at low multiplicity (goal transduction rate of 30-50%) to ensure single integrants (i.e., one sgRNA per cell)^82–84^. Alternatively, lentiviral transduction of Cas9-expressing cells can occur at high multiplicity (transduction rate >50%) to obtain a higher rate of transduced cells with a resulting heterogeneous mix of sgRNA copy number. For an arrayed experiment with one sgRNA, multiple integrants per cell do not result in “passenger” sgRNAs and associated false positives. Lentiviral transduction efficiency is affected by cell density, volume, incubation time, and multiplicity of infection (MOI). Therefore, it is important to keep all of these factors constant to ensure consistent lentiviral transduction rates. Notably, this underscores the importance of aliquoting lentivirus to ensure consistent lentiviral titer since titer is decreased by freeze/thaw cycles (see Steps 53, 64). A decrease in cell density, decrease in media volume, and increase in MOI will all increase transduction rates. It is recommended that a “kill curve” is created to determine the optimal concentration of puromycin for the cells used prior to beginning the experiment. It is also important to determine the duration of selection required to complete cell death by puromycin. Longer exposure to the CRISPR/Cas9 reagents results in increased editing rates^14^. A reasonable experiment duration is 1-2 weeks^14^; however, this may vary based on the experimental system used. It is possible to use a Cas9 nuclease activity reporter as described above to more accurately determine when editing has plateaued in the experimental system. At the end of the experiment, cell pellets can be made to proceed with deep sequencing (Steps 66-74).

**Enhancing lentiviral transduction**: Multiple methods exist to enhance lentiviral transduction. Reagents such as polybrene, rapamycin, protamine sulfate, and prostaglandin E2 have been shown to enhance lentiviral transduction rates^85^. In addition, lentiviral spin-infection (centrifugation during transduction) can enhance transduction rates. These types of methodologies to enhance lentiviral transduction can be useful to reduce the amount of lentivirus required for experiments, can help achieve the desired transduction rates in the setting of low lentiviral titer, and can increase efficiency for difficult to transduce cells. It is important to determine if any of these reagents/methods lead to cellular toxicity.

**Box 4** | Technical and experimental considerations for performing a pooled CRISPR genome editing experiment.

Here are some practical guidelines for executing a pooled screen using an sgRNA library with 1,000 sgRNAs with a goal of 1,000x representation of the sgRNA library. A experimental schematic can be found in Figure 2 and an example workflow in Figure 3:

**Generation of cells with stable CRISPR nuclease expression**: For cell lines, it is convenient to generate lines with stable CRISPR nuclease expression for arrayed experiments. This can be accomplished by transducing cells with a lentiviral CRISPR nuclease with subsequent selection for transduced cells, such as the usage of lentiviral Cas9 with blasticidin resistance (the focus of the remainder of the discussion in Box 4 will focus on the usage of *S. pyogenes* Cas9; however, the same principles apply to other CRISPR nucleases including Cpf1). It is recommended that a “kill curve” is created to determine the optimal concentration of blasticidin for the cells used prior to beginning the experiment. It is also important to determine the duration of selection required to complete cell death by blasticidin. Stable expression of Cas9 can be confirmed via Western blot. Alternatively, stable expression can be confirmed by assessing Cas9 function using a reporter system, such as the previously described constructs that provide GFP and an sgRNA targeting GFP to assess for functional Cas9 through flow cytometry (see Materials)^12,46^. It is possible to screen for clones with high Cas9 expression and/or high Cas9 activity as assessed by a reporter; however, it is not required. If stable Cas9 expression cannot be generated, such as usage of primary cells with limited culture duration, co-transduction of Cas9 and sgRNAs can be performed with double selection (blasticidin for Cas9 and puromycin for sgRNA). Co-transduction can occur simultaneously or can occur on back-to-back days. Selection by blasticidin and/or puromycin is typically added 24-48 hours after transduction.

**sgRNA library representation**: Library representation refers to estimating how frequently each sgRNA in the library is included in the experiment. 1000x representation of a library suggestions that 1,000 cells were transduced by the median sgRNA in the library at the beginning of the experiment. Suboptimal representation may lead to the absence of certain sgRNAs from the screen experiment and thus can cause sgRNAs to appear falsely depleted from the experiment. The desired level of representation may reflect the expected uniformity of genome editing outcome and biologic phenotype as well as the distribution of sgRNA abundance within the library. For example, true positive sgRNAs expected to have uniform genetic and biological effect and narrowly distributed abundance may require lower representation to be accurately identified as hits. To represent a 1,000 sgRNA library at 1000x, one million cells must be transduced (1,000 sgRNAs * 1,000 cells each = 1,000,000 cells).

**Transduction at low multiplicity**: Lentiviral transduction of Cas9-expressing cells at low multiplicity ensures single integrants (i.e., one sgRNA per cell)^82–84^. As such, the goal transduction rate is 30-50%. Transduction rates >50% increase the risk of multiple integrants per cell, which can result in “passenger” sgRNAs and associated false positives. Lentiviral transduction efficiency is affected by cell density, volume, incubation time, and multiplicity of infection (MOI). Therefore, it is important to keep all of these factors constant to ensure consistent lentiviral transduction rates. Notably, this underscores the importance of aliquoting lentivirus to ensure consistent lentiviral titer since titer is decreased by freeze/thaw cycles (see Steps 53, 64). The example screen requires transduction of 1,000,000 cells to represent the sgRNA library at 1,000x; however, this is referring to the number of transduced cells. Since the transduction rate must be between 30-50%, this requires using 2,000,000-3,333,333 cells at the beginning of the experiment, which will be reduced to 1,000,000 cells upon selection by puromycin for successful transductants.

**Determination of screen conditions**: For this example, the goal will be a 40% transduction rate. Therefore, the experiment will require 2,500,000 cells (2,500,000*0.4 = 1,000,000). As previously described, lentiviral transduction efficiency is affected by cell density, volume, incubation time, lentiviral titer, and MOI. Constant titer will be assumed due to appropriate lentivirus aliquoting (see Steps 53, 64). Increasing media volume results in lower transduction. Therefore, it is useful to limit the amount of media. Also, it can be simpler to perform the transduction in “parts” (e.g., 10 separate transductions of 250,000 cells). A reasonable approach would include transducing 10 wells in a 24 well plate of 250,000 cells in 500 µL of medium (cell density of 500,000 cells/mL). The use of a 24 well plate and the cell density of 500,000 cells/mL are reasonable approaches; however, these may need to be optimized for the cells used for study and/or experimental goals. Transductions should be performed at different MOI to determine the MOI to achieve the goal transduction rate of 40%.

**Enhancing lentiviral transduction**: Multiple methods exist to enhance lentiviral transduction. Reagents such as polybrene, rapamycin, protamine sulfate, and prostaglandin E2 have been shown to enhance transduction rates^85^. In addition, lentiviral spin-infection (centrifugation during transduction) can enhance transduction rates. These types of methodologies to enhance lentiviral transduction can be useful to reduce the amount of lentivirus required for experiments, can help achieve the desired transduction rates in the setting of low lentiviral titer, and can increase efficiency for difficult to transduce cells. It is important to determine if any of these reagents/methods lead to cellular toxicity. If a reagent/method is used, it should be used for all transduction samples. In addition, the reagent/method should be used when determining the MOI to achieve the goal transduction rate (40% in this case).

**Pooled sgRNA screen experiment execution**: For this example, it will be assumed that an MOI of 1 results in a 40% transduction rate. To perform the screen, 10 wells in a 24 well plate with 250,000 cells in 500 µL of medium (cell density of 500,000 cells/mL) are transduced at an MOI of 1. 24 hours after transduction, the 10 wells can be pooled into one larger well or flask. Prior to selection for successful transduction, it is useful to remove a subset of cells from the pooled transductions to determine transduction efficiency to ensure ~40% transduction was empirically achieved. A reasonable approach would include pooling all 10 wells together for a total of 2,500,000 cells in 5 mL of medium 24 hours after transduction. ~25,000-50,000 cells can be removed from this 5 mL cell mixture to be used to determine transduction rate by splitting the cells into with and without puromycin conditions to confirm the transduction rate by cell counts post-selection. After removal of ~25,000-50,000 cells, puromycin selection should be begun on the all cells in the well/flask. It is recommended that a “kill curve” is created to determine the optimal concentration of puromycin for the cells used prior to beginning the screen experiment. It is also important to determine the duration of selection required to complete cell death by puromycin. Longer exposure to the CRISPR/Cas9 reagents results in increased editing rates^14^. A reasonable screen duration is 1-2 weeks^14^; however, this may vary based on the experimental system used. It is possible to use a Cas9 nuclease activity reporter as described above to more accurately determine when editing has plateaued. At the end of the experiment, cell pellets can be made to proceed with deep sequencing (Steps 66-74).

**Types of Screens:** Screens typically rely on either positive or negative selection. Common screen strategies involve determination of enrichment or dropout (‘depletion’) of sgRNAs. This can be achieved by deep sequencing either the plasmid pool or cells at an early time point in the experiment to serve as the initial time point for comparison. It has been previously shown that there is no difference between using the plasmid pool versus cells from an early time point in the experiment for this purpose^14,86^. Deep sequencing of samples at the end of the experiment can then be compared to this initial time point. The sgRNA presence can be determined by enumerating sgRNAs using CRISPRessoCount (Fig. 2-3, Steps 77-81). Using enrichment/dropout strategies, screens can be performed for such applications as drug/toxin resistance/susceptibility or fluorescence-activated cell sorting based selection (using antibody or cell reporter). A summary of representative published screens is summarized in Joung et al^20^.

## ACKNOWLEDGMENTS

M.C.C. is supported by a National Institute of Diabetes and Digestive and Kidney Diseases (NIDDK) Award (F30DK103359-01A1). M.H. is funded by grants NIH/NHGRI 5U41HG002371-15 and NIH/NCI 5U54HG007990-02 and by a grant from the California Institute of Regenerative Medicine, CIRM GC1R-06673C.). D.E.B. is supported by NIDDK (K08DK093705, R03DK109232), NHLBI (DP2OD022716), Burroughs Wellcome Fund, Doris Duke Charitable Foundation Innovations in Clinical Research Award, ASH Scholar Award, Charles H. Hood Foundation Child Health Research Award, and Cooley’s Anemia Foundation fellowship. S.H.O. is supported by an award from the NHLBI award (P01HL032262) and an award from the NIDDK (P30DK049216, Center of Excellence in Molecular Hematology). N.E.S. is supported by the NIH through NHGRI (R00-HG008171). L.P. is supported by a National Human Genome Research Institute (NHGRI) Career Development Award (R00HG008399) and the Defense Advanced Research Projects Agency (HR0011-17-2-0042).

## AUTHOR CONTRIBUTIONS

M.C.C., M.H., and L.P. conceived this project. M.H. and J-P.C. created CRISPOR. L.P., D.B. and G-C.Y created CRISPResso. M.C.C. and D.B. performed the experiments. M.C.C., D.B., S.H.O., N.E.S., O.S., G-C.Y., F.Z. and L.P. analyzed the experimental data. M.C.C., M.H., and L.P. wrote the manuscript with input from all authors.

---

**Supplementary Fig. 1: Screenshot for sharing disk volumes with Docker.** Screenshot using Docker to ensure that the drive(s) you want to be available to the container is(are) checked (under Settings…/Shared Drives).

**Supplementary Fig. 2: Screenshot for allocation of memory for Docker containers.** Screenshot using Docker to allocate enough memory to the container (under Settings…/Advanced).

**Supplementary Fig. 3: Pooled sgRNA library preparation and analysis. a**, Representative results for lsPCR1. b, Representative results for lsPCR2. c, Gradient PCR for locus-specific primers for laPCR1. d, Representative results for laPCR2.

**Supplementary Fig. 4: Visualization of the distribution of identified alleles generated from targeting *BCL11A* exon 2.** Nucleotides are indicated by unique colors (A = green; C = red; G = yellow; T = purple). Substitutions are shown in bold font. Red rectangles highlight inserted sequences. Horizontal dashed lines indicate deleted sequences. The vertical dash line indicates the predicted double-strand break position.

**Supplementary Fig. 5: Direct comparison of *BCL11A* exon 2 sequence between a *BCL11A* exon 2 targeted sgRNA sample (“edited”) and a non-edited control sample (“non-edited”). a**, Distribution of editing outcomes (unmodified, NHEJ, HDR, and mixed alleles) for treated (edited) and control (non-edited) samples. b, Comparison of the percent different editing outcomes (unmodified, NHEJ, HDR, and mixed alleles) between the treated (edited) and control (non-edited) samples. c, Combined (substitutions/deletions/insertions) mutation position distribution for treated (edited) and control (non-edited) samples. The vertical dashed line indicates the position of predicted Cas9 cleavage. The position of the sgRNA is shown in gray. d, Comparison of the percent different combined mutations (substitutions/deletions/insertions) between the treated (edited) and control (non-edited) samples. The vertical dashed line indicates the position of predicted Cas9 cleavage. The position of the sgRNA is shown in gray.

